# Noise analysis of derivative-action biomolecular topologies

**DOI:** 10.64898/2026.05.06.723344

**Authors:** Emmanouil Alexis, Sebastián Espinel-Ríos, Luca Laurenti, Luca Cardelli, Ioannis G. Kevrekidis, Clarence W. Rowley, José L. Avalos

**Affiliations:** Omenn-Darling Bioengineering Institute, Princeton University, New Jersey, USA; Department of Chemical and Biological Engineering, Princeton University, New Jersey, USA; School of Chemical and Bioprocess Engineering, University College Dublin, Dublin, Ireland; Delft Center for Systems and Control, Delft University of Technology, Delft, Netherlands; The Italian Institute of Artificial Intelligence for Industry (AI4I), Turin, Italy; Department of Computer Science, University of Oxford, Oxford, UK; Department of Chemical and Biomolecular Engineering, Johns Hopkins University, Baltimore, USA; Department of Applied Mathematics and Statistics, Johns Hopkins University, Baltimore, USA; Department of Mechanical and Aerospace Engineering, Princeton University, New Jersey, USA; High Meadows Environmental Institute, Princeton University, New Jersey, USA; The Andlinger Center for Energy and the Environment, Princeton University, New Jersey, USA; Department of Molecular Biology, Princeton University, New Jersey, USA

## Abstract

Temporal gradient sensing is a fundamental capability observed across diverse natural biological systems, contributing to the coordination of their functions. Harnessing this ability is also of significant interest in synthetic biology, particularly for sensing and control applications. In this work, we focus on a biomolecular topology that exemplifies a broader class of signal-differentiating architectures, while introducing a structural variant of it. We examine their behavior under both nominal and non-ideal conditions, accounting for stochastic noise arising from different sources. Our investigation includes scenarios where these topologies operate independently, as well as when embedded within minimal regulatory architectures based on negative as well as positive feedback. We analyze the stability of the resulting macroscopic dynamics—a prerequisite for practical deployment—and quantify stochastic fluctuations in system output, providing comparisons with the corresponding input/unregulated process. Importantly, our results demonstrate that signal differentiation can be effectively implemented in a biomolecular setting without incurring deleterious noise amplification—a major concern in the utilization of derivative action across disciplines.

## 1 Introduction

In recent years, the design of biomolecular topologies capable of temporal gradient sensing has gained increasing attention within synthetic biology. This comes as no surprise, as accurate estimation of time derivatives of a system’s output can reveal important aspects of its properties and can be leveraged for tight regulation. This concept is well-established in traditional engineering approaches used for the design and control of electromechanical systems, some of which might inspire analogous biomolecular implementations. Interestingly, temporal gradient sensing is also abundant in natural biological systems. For instance, small motile bacteria (e.g., *Escherichia coli*) compute temporal gradients of nutrient concentrations to guide their swimming behaviour—a defining feature of a mechanism known as *chemotaxis* [1–3].

A promising synthetic biology application of derivative-action biomolecular topologies is the development of rate-of-change biosensors (e.g., speedometers or accelerometers) enabling real-time monitoring of, for example, uptake or secretion rates of biomolecules of interest [4]. Another promising application is the realization of derivative action-based biomolecular control architectures. Notably, Proportional-Integral-Derivative (PID) control laws are the “bread and butter” of feedback control engineering practice, where derivative action is typically used to improve closed-loop characteristics such as stability and transient dynamics [5, 6].

Several theoretical approaches for implementing topologies capable of signal differentiation and derivative control for diverse *in vitro* and/or *in vivo* applications have recently been proposed [7– 21]. These approaches include various types of biomolecular interactions following mass-action and/or Hill-type kinetics. In parallel, the useful information—corresponding to the computed temporal gradients—is represented in different ways, namely as a biomolecular species, the non-physical difference between two such species, or a chemical reaction rate. A comparative discussion of the above methodologies can be found in [22]. Most of the aforementioned studies rely on deterministic analysis; in the few cases where the inherent stochasticity of chemical reactions is considered, the investigation is limited to numerical simulations within derivative-control-based settings. An exception is the study in [21], where, among other results, the authors derive analytical expressions quantifying the stationary stochastic noise levels of a negative-feedback derivative-control scheme tailored to a gene expression process.

In this work, we focus on a specific family of such biomolecular motifs, known as *BioSD* (*Biomolecular Signal Differentiators*), originally introduced in [7]. Specifically, we are concerned with a biomolecular topology, called *BioSD*-*II*. This topology can function as a general purpose signal differentiator bio-device with low-pass filtering action and can be utilized for the design of derivative control regulatory strategies [7, 9]. Our choice to work with this particular topology is motivated by the fact that, compared to other members of its family, *BioSD*-*II* has a simpler structure and appears more appealing for experimental implementations. Potential *in vivo* realizations have been explored in [7], while a suitable *in vitro* implementation methodology based on DNA strand displacement can be found in [20, 23]. Note that the temporal gradient computations here are encoded as biomolecular species concentrations, thereby facilitating direct experimental measurements. In addition, we introduce and analyze a variant topology, termed *AC*-*BioSD*-*II* (*Autocatalytic*-*BioSD*-*II*), in which the signal to be differentiated is applied via an autocatalytic reaction [10].

The main aim of this paper is to investigate the behaviour of *BioSD*-*II* and *AC*-*BioSD*-*II* in the presence of noise, which manifests itself as stochastic fluctuations. There are two sources of these fluctuations, both of which are taken into account: the probabilistic nature of chemical reactions firing within the topology under consideration itself (intrinsic noise) and the randomizing effects from the interaction of this topology with its external environment (extrinsic noise) [24–28]. We examine both *BioSD*-*II* and *AC*-*BioSD*-*II* across various scenarios: as standalone modules operating under nominal conditions, under commonly encountered non-idealities, and in both the presence and absence of an input stimulus. Furthermore, we explore different regulatory architectures based on negative and positive feedback, in which these modules play the role of bio-controllers. Our analysis offers valuable insights into how noise propagates within these settings, particularly concerning the biomolecular species responsible for carrying the information of interest. These findings are highly relevant, especially for future experimental implementations, as undesired noise sensitivity is often a critical concern in the practical deployment of derivative action, regardless of the application context [6, 29].

Except for some very simple cases, the exact description of a set of chemical reactions in the stochastic setting—e.g., via the Chemical Master Equation (CME)—results in mathematical models that are analytically intractable [30, 31]. To overcome this obstacle, we employ a widely used approach, namely the Linear Noise Approximation (LNA) of the CME, to evaluate stochastic fluctuations around macroscopically stable trajectories of interest, given by (deterministic) reaction rate equations (RRE) [32–36]. To quantify noise, we adopt the Fano factor, defined as the variance-to-mean ratio [37, 38], which also enables direct comparison with Poisson noise, for which this metric equals unity (since the values of the mean and variance are equal). To validate our LNA-based results, we perform computational analysis using Gillespie’s stochastic simulation algorithm (SSA) [39], which faithfully approximates the aforementioned CME. In addition, through RRE-based models, we investigate practically important properties of the mean behaviour, such as the capacity for signal differentiation/attenuation, regulatory action and stability. Further details on our mathematical and computational methods are provided in Sections S1 and S2 of the Supplementary Material. Moreover, an overview of the types and representations of the chemical reactions used in this study is presented in Fig. 1a.

**Figure 1:**
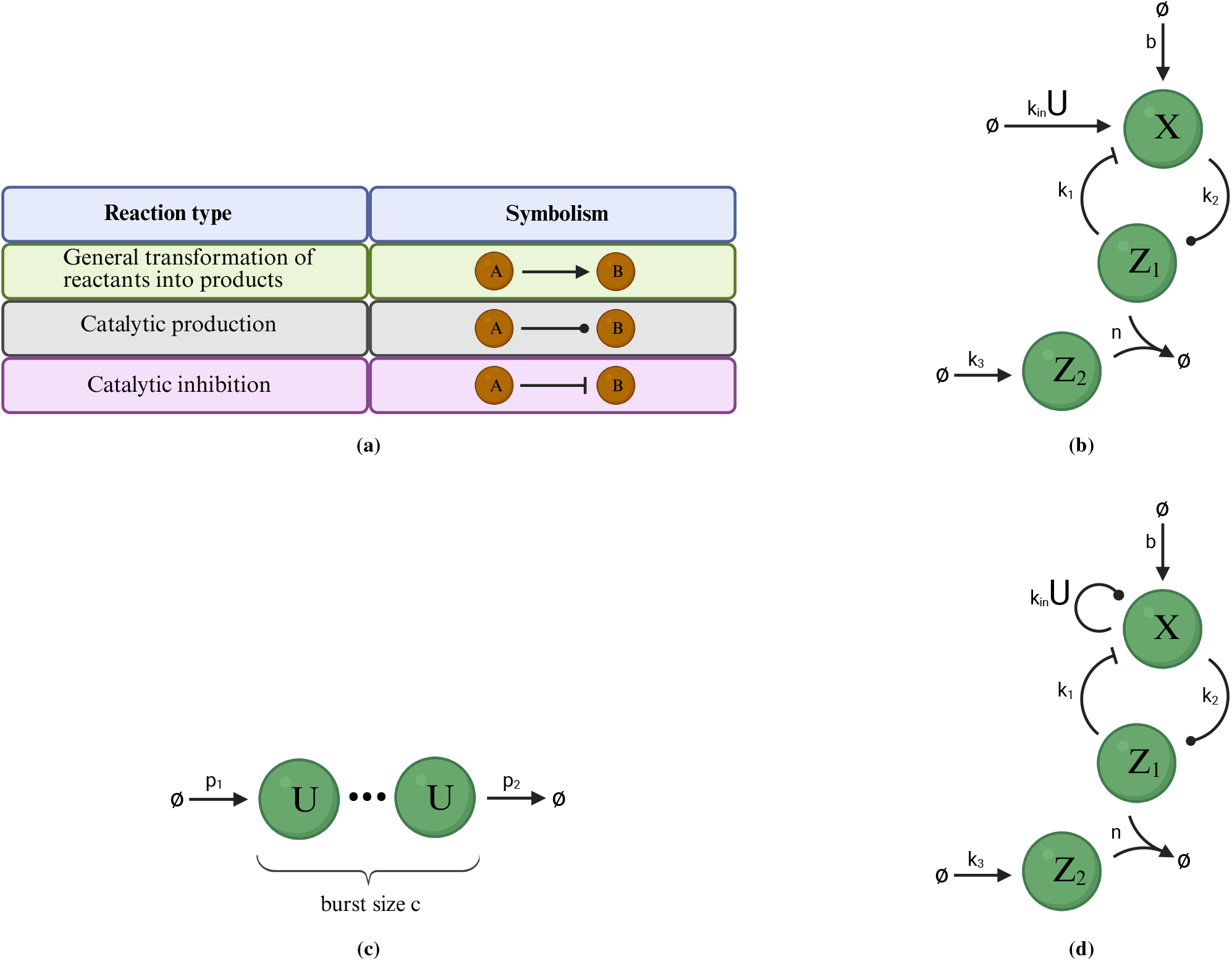
Biomolecular topologies under investigation. Schematic of (a) the different types of chemical reactions considered in this study; (b) the *BioSD*-*II* module; (c) a bursty birth–death biomolecular process; (d) the *AC*-*BioSD*-*II* module.

## 2 Behaviour under nominal conditions

### 2.1 The *BioSD-II* topology

The chemical reaction network (CRN) describing the *BioSD*-*II* topology (Fig. 1b) is:

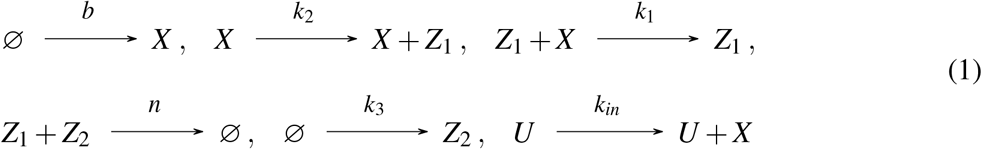

where *U, X* represent the input and the output of *BioSD*-*II*, respectively. By focusing on the corresponding RREs, one can clearly observe the filtered derivative action at play. In particular, it can be shown [7, 9] that around an equilibrium of interest, 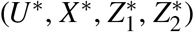, where ^∗^ denotes the steady state of a variable, the input/output relation can be described in the Laplace domain as:

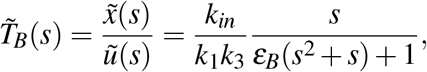

where 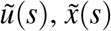 are the Laplace transform of *u* = *U* − *U* ^∗^, *x* = *X* − *X* ^∗^, respectively, *s* is the Laplace variable (complex frequency) and 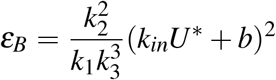. The transfer function 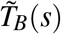 embodies a signal differentiation process, accompanied by the action of a second-order low-pass filter. Note that 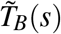 holds for *n* → ∞, which results in the practical constraint 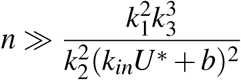. Experimental mechanisms for implementing (sufficiently fast) annihilation reactions, such as 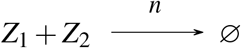, include, among others, split intein-based [40–42] and DNA strand displacement-based [20, 23, 43] reactions.

We now turn our focus to the stochastic fluctuations and examine the special case in which no input signal is present, i.e. *U* = 0. Assuming *n* → ∞, we calculate the Fano factor of the output, *X*, at steady state as:

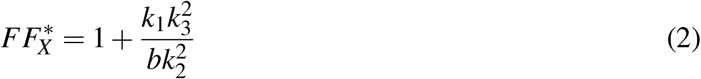

which indicates the presence of super-Poisson noise, as 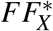 is always greater than unity. A system of ordinary differential equations (ODEs) approximating the time evolution of *FF*_*X*_ (*t*) is provided in Section S3 of the Supplementary Material.

Next, to study the behaviour of *BioSD*-*II* with respect to both intrinsic and extrinsic noise, we consider an input signal modelled as a general bursty birth–death biomolecular process, comprising the chemical reactions:

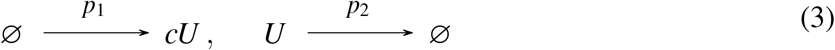

where the constant *c* ∈ ℕ^+^ denotes the burst size (Fig. 1c). The Fano factor of *U* at steady state can be (exactly) calculated as:

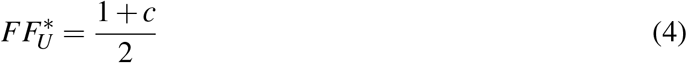

It is clear that the value of 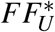 and, by extension, the type of the input noise is completely determined by the burst size. For instance, *c* = 1 results in 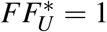 entailing Poisson noise. Note that (bursty) birth-death processes are commonly employed to model important biomolecular phenomena such as protein production [44–47].

The topology described by the CRNs in Eqs. (1), (3) results in the following steady-state output Fano factor (assuming *n* → ∞):

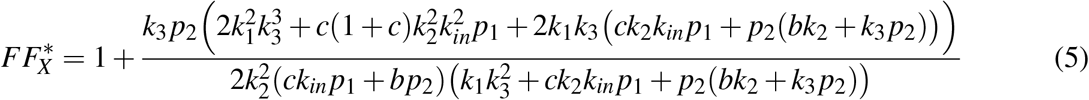

which shows that the noise remains above Poisson levels 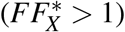. An ODE system for *FF*_*X*_ (*t*) can be found in Section S3 of the Supplementary Material.

### 2.2 The *AC-BioSD-II* topology

We introduce the *AC*-*BioSD*-*II* topology (Fig. 1d) whose CRN is:

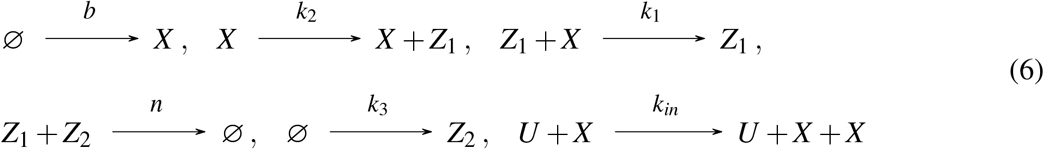

In *AC*-*BioSD*-*II* the input signal is applied through an autocatalytic excitation process 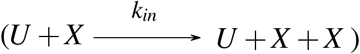 which constitutes the only structural difference between the CRNs in Eq. (1) and (6). This modification is motivated by [10], where this concept was studied for the first time within a different *BioSD* architecture.

Using the corresponding RREs, we analyze the main characteristics of this new topology in Section S4 of the Supplementary Material. As shown, the input/output relation around an equilibrium of interest 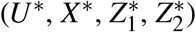, can be described in the Laplace domain by the transfer function:

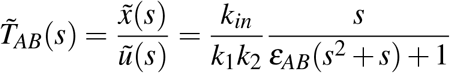

Where 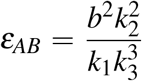. Observe that, although 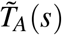 and 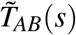 have the same form, in contrast to *ε*_*B*_, *ε*_*AB*_ depends exclusively on the rates of the internal reactions and is independent of the input excitation process. This property can considerably facilitate the tuning of the embedded low-pass filter, providing a significant practical implementation advantage. 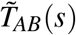 has also been calculated assuming *n* → ∞, which results in the practical constraint 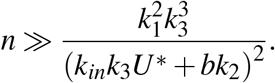.

In the absence of an input signal, Eq. (2) holds, since the CRNs in Eqs. (1) and (6) coincide. If we now consider the input signal given by the CRN in Eq. (3), we can obtain for the Fano factor of *X* (see the CRN given by Eq. (6)) at steady state (in the limit as *n* → ∞):

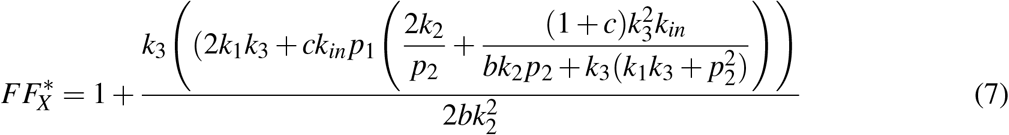

which indicates the presence of super-Poisson noise 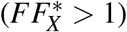. An ODE system for *FF*_*X*_ (*t*) is provided in Section S4 of the Supplementary Material.

In Figs. 2, S1, we computationally demonstrate the results presented in Sections 2.1 and 2.2 using a simulation example. We applied the LNA, validated against the SSA, to examine both the noise levels and the signal differentiation capacity with respect to the biomolecular topologies under investigation. A noteworthy point is that, under external stimulation, it is possible for the output of our topologies to approximate the input’s first derivative with high fidelity while exhibiting markedly lower noise levels (than those of the input). Finally, in Section S5 of the Supplementary Material, we discuss an approach to approximate the output Fano factors, *FF*_*X*_ (*t*), with explicit formulas, which can provide further insight into the transient dynamics.

**Figure 2:**
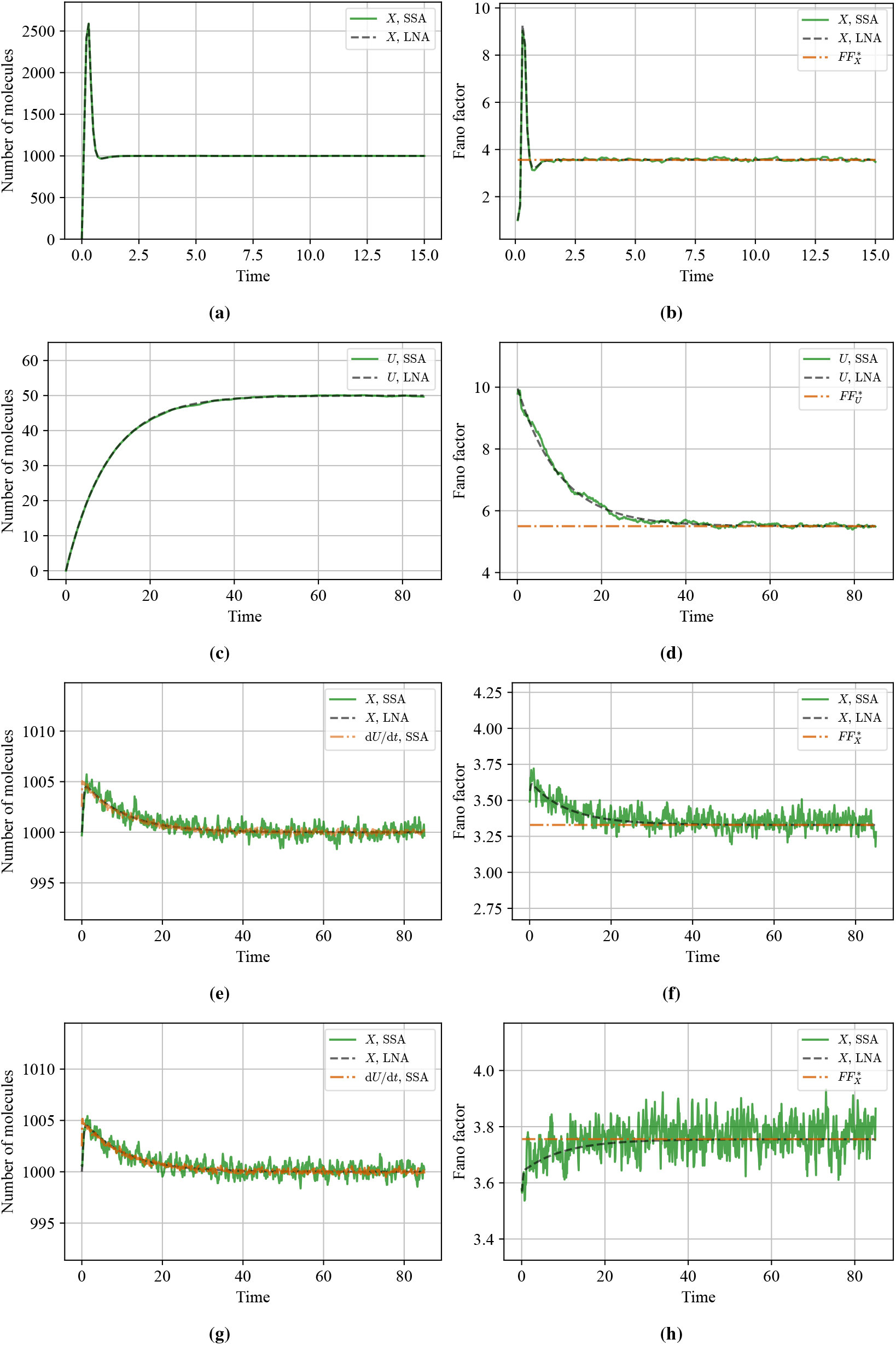
Nominal behavior. Time evolution of the mean molecular copy number and the Fano factor (together with the corresponding analytically derived steady-state approximation), obtained using SSA and LNA for: (a,b) the output of the *AC*-*BioSD*-*II* module in the absence of input stimulation (CRN in Eq. (1) with *U* = 0), using [7] *b* = 250, *k*_1_ = 1.6, *k*_2_ = 1, *k*_3_ = 20, and *n* = 1000 (moderately large); (c,d) a general bursty birth–death biomolecular process (CRN in Eq. (3)) with *c* = 10, *p*_1_ = 0.01, and *p*_2_ = 0.1; (e,f) the output response of the *BioSD*-*II* module to an input signal given by (c,d), with *k*_in_ = 32 and all remaining parameters unchanged; (g,h) the output response of the *AC*-*BioSD*-*II* module to the same input signal, with *k*_in_ = 1.6 and all remaining parameters unchanged. In panels (e) and (g), the first derivative of the input signal (shifted by the output steady-state value) is also shown while the differentiators’ output gain is equal to 1, facilitating visual comparisons. The initial conditions in (a,b) and (c,d) are set to zero, whereas those in (e,f) and (g,h) are set to the corresponding steady-state values from (a,b). Individual stochastic trajectories for the above simulation scenarios are presented in Fig. S1 of the Supplementary Material.

## 3 Behaviour under non-idealities

### 3.1 Effects of external degradation sources

First, we consider the scenario in which the output of our topologies is affected by a degradation mechanism, modeled as 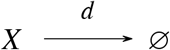. For instance, if the output corresponds to a protein within the cell, such degradation can occur due to the action of a protease [48, 49]. This alters the conditions under which the dynamics governed by the RREs remain stable. Interestingly, however, assuming stability is maintained, both the signal differentiation ability (measured in terms of mean behaviour) and the steady-state output Fano factor, in the limit as *n* → ∞, remain unchanged (see Figs. 3-5 and S3-S5).

**Figure 3:**
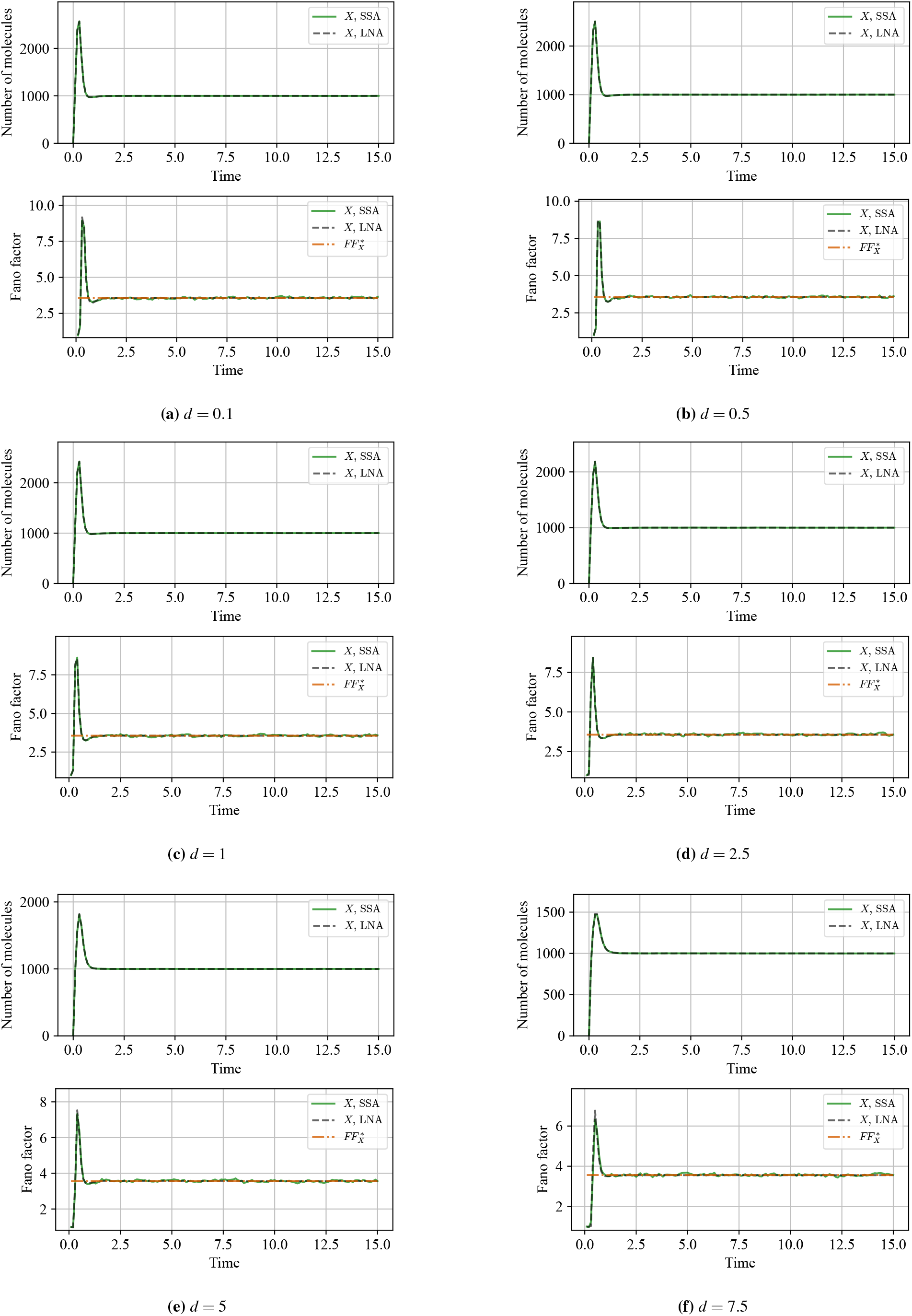
Effect of output degradation in the absence of input stimulation. The simulation scenario of Fig. 2(a,b) is repeated for different values of the output degradation rate constant *d*. Individual stochastic trajectories are presented in Fig. S3 of the Supplementary Material.

**Figure 4:**
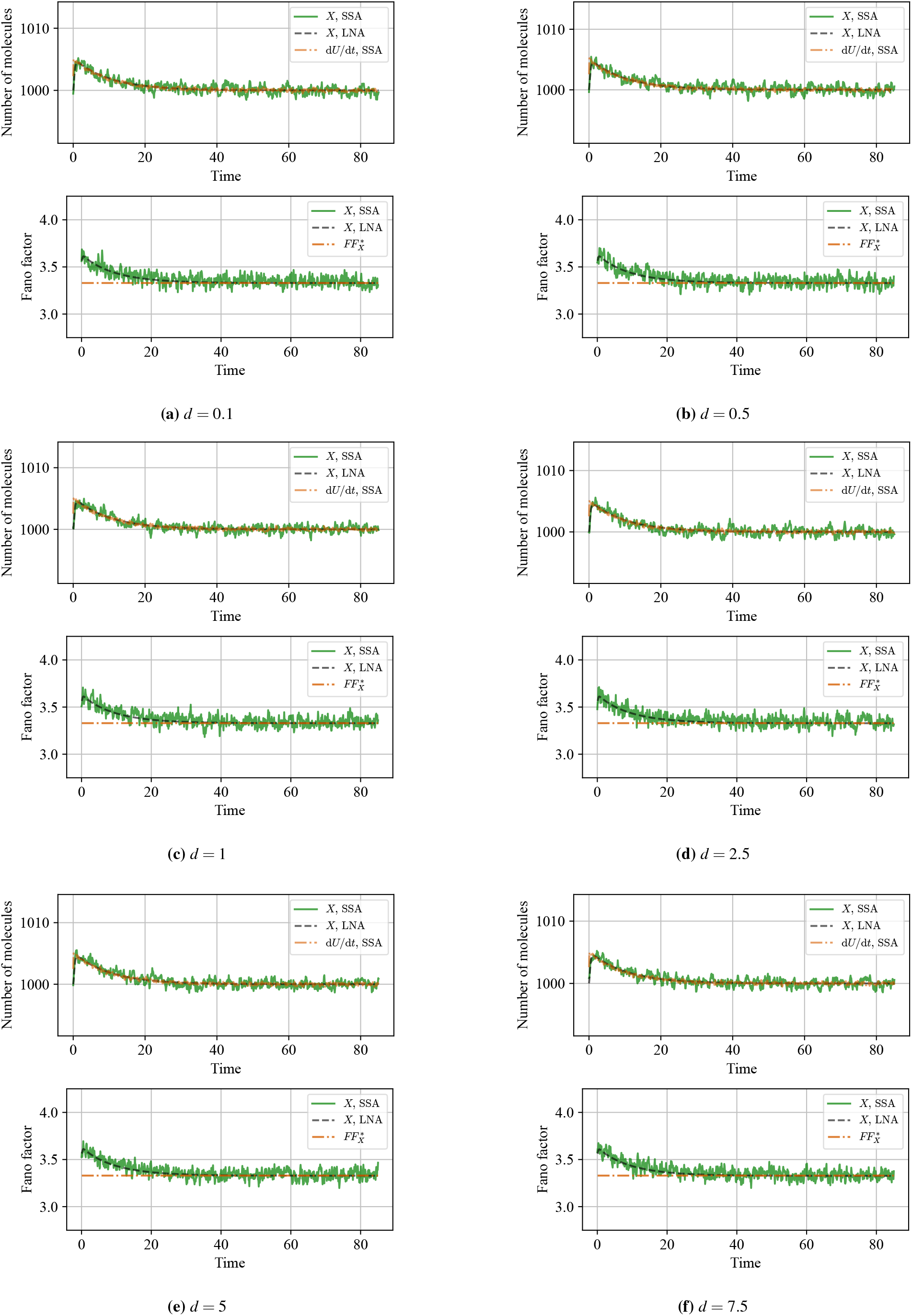
*BioSD-II* - Effect of output degradation under input stimulation. The simulation scenario of Fig. 2(e,f) is repeated for different values of the output degradation rate constant *d*. Individual stochastic trajectories are presented in Fig. S4 of the Supplementary Material.

**Figure 5:**
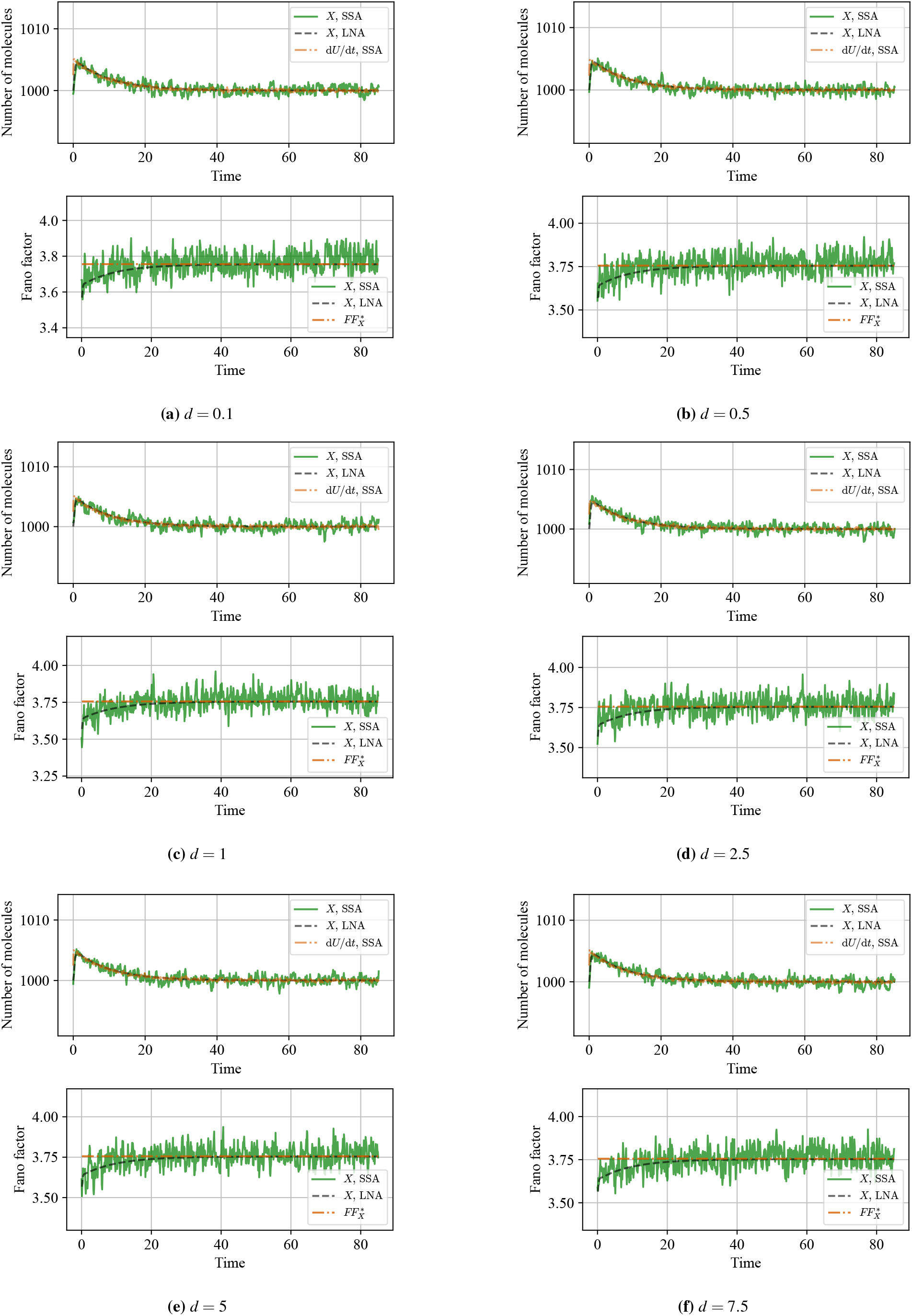
*AC-BioSD-II* - Effect of output degradation under input stimulation. The simulation scenario of Fig. 2(g,h) is repeated for different values of the output degradation rate constant *d*. Individual stochastic trajectories are presented in Fig. S5 of the Supplementary Material.

In particular, for the *BioSD*-*II* topology, defined now by the CRN in Eq. (1) and the aforementioned degradation reaction, it is shown in [7] that, for any 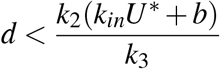, the system is stable and 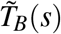 is valid. As for the *AC*-*BioSD*-*II* topology, defined now by the CRN in Eq. (6) and the aforementioned degradation reaction, we show in Section S6 of the Supplementary Material that the system stability and the validity of 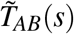 are maintained for any 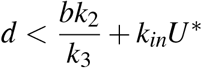. As can be seen, when no input signal is applied, the above cases coincide, and the corresponding stability condition becomes 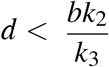. In the same Section, we also provide ODE systems for the corresponding *FF*_*X*_ (*t*), from which we can exactly recover Eqs. (2), (5), (7).

Another interesting and more complex scenario arises when all species within our topologies are subject to degradation. This situation, which may represent, for instance, the phenomenon of dilution due to cell growth [48, 49], involves three additional degradation reactions instead of one, namely 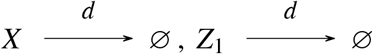, and 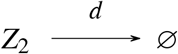. In Figs. 6-8 and S6-S8, we computationally investigate this scenario with respect to the output noise levels and we demonstrate the distortion of the signal differentiation process, which becomes increasingly corrupted as *d* grows (for further details on the corresponding *FF*_*X*_ (*t*), see Section S6 of the Supplementary Material).

**Figure 6:**
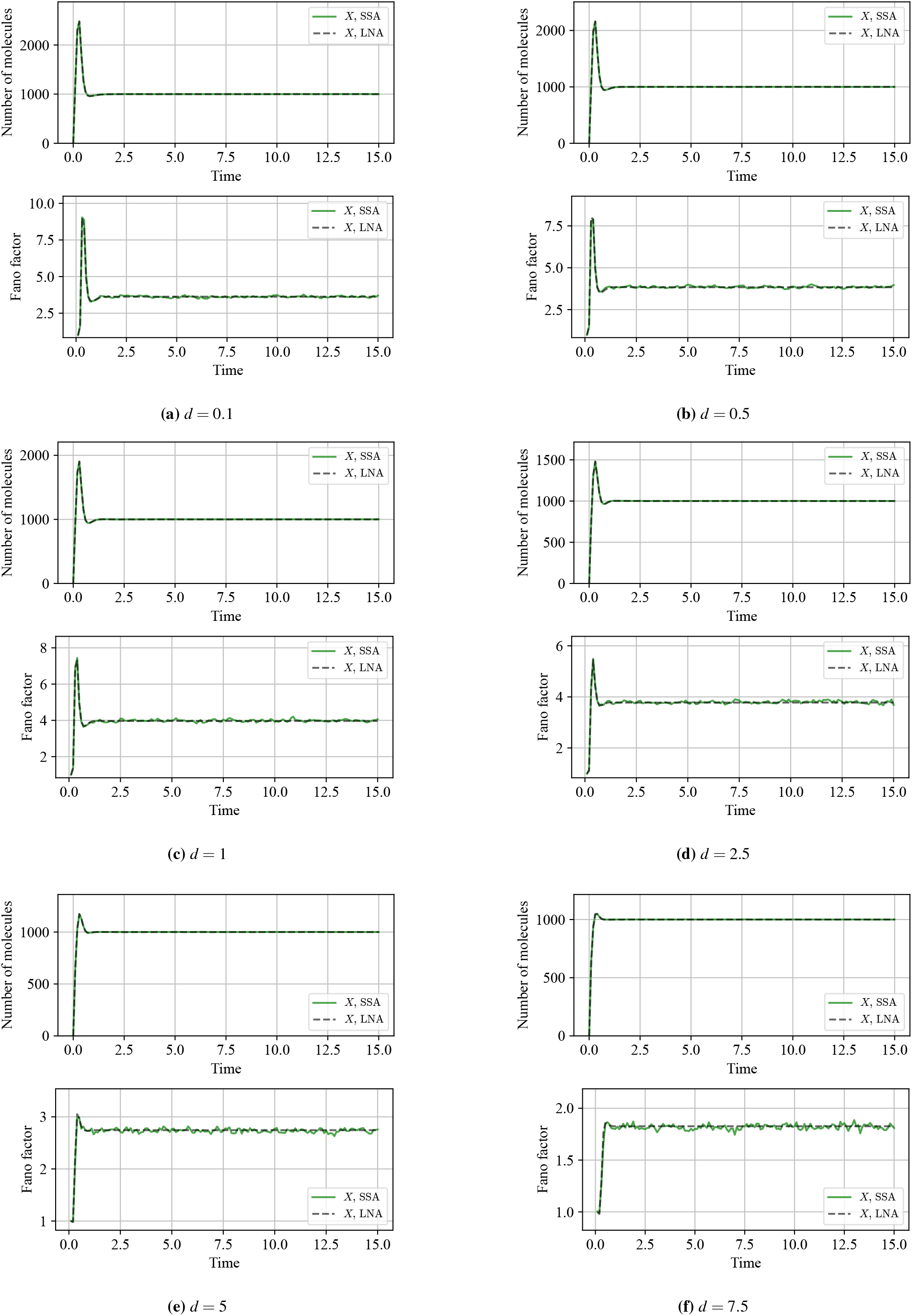
Effect of degradation of all species in the absence of input stimulation. The simulation scenario of Fig. 2(a,b) is repeated for different values of the common degradation rate constant *d*. Individual stochastic trajectories are shown in Fig. S6 of the Supplementary Material.

**Figure 7:**
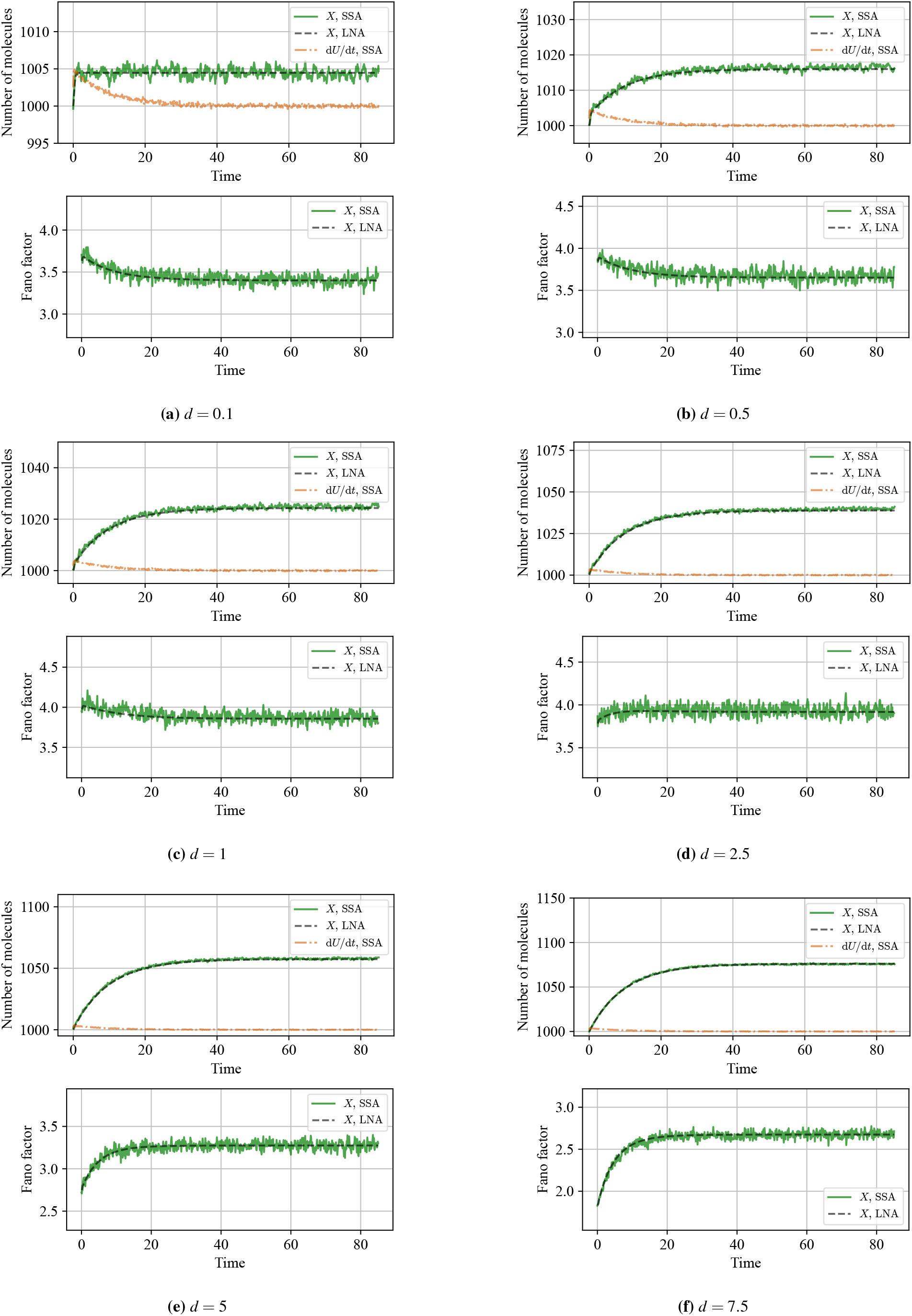
*BioSD-II* - Effect of degradation of all species under input stimulation. The simulation scenario of Fig. 2(e,f) is repeated for different values of the common degradation rate constant *d*. Individual stochastic trajectories are presented in Fig. S7 of the Supplementary Material.

**Figure 8:**
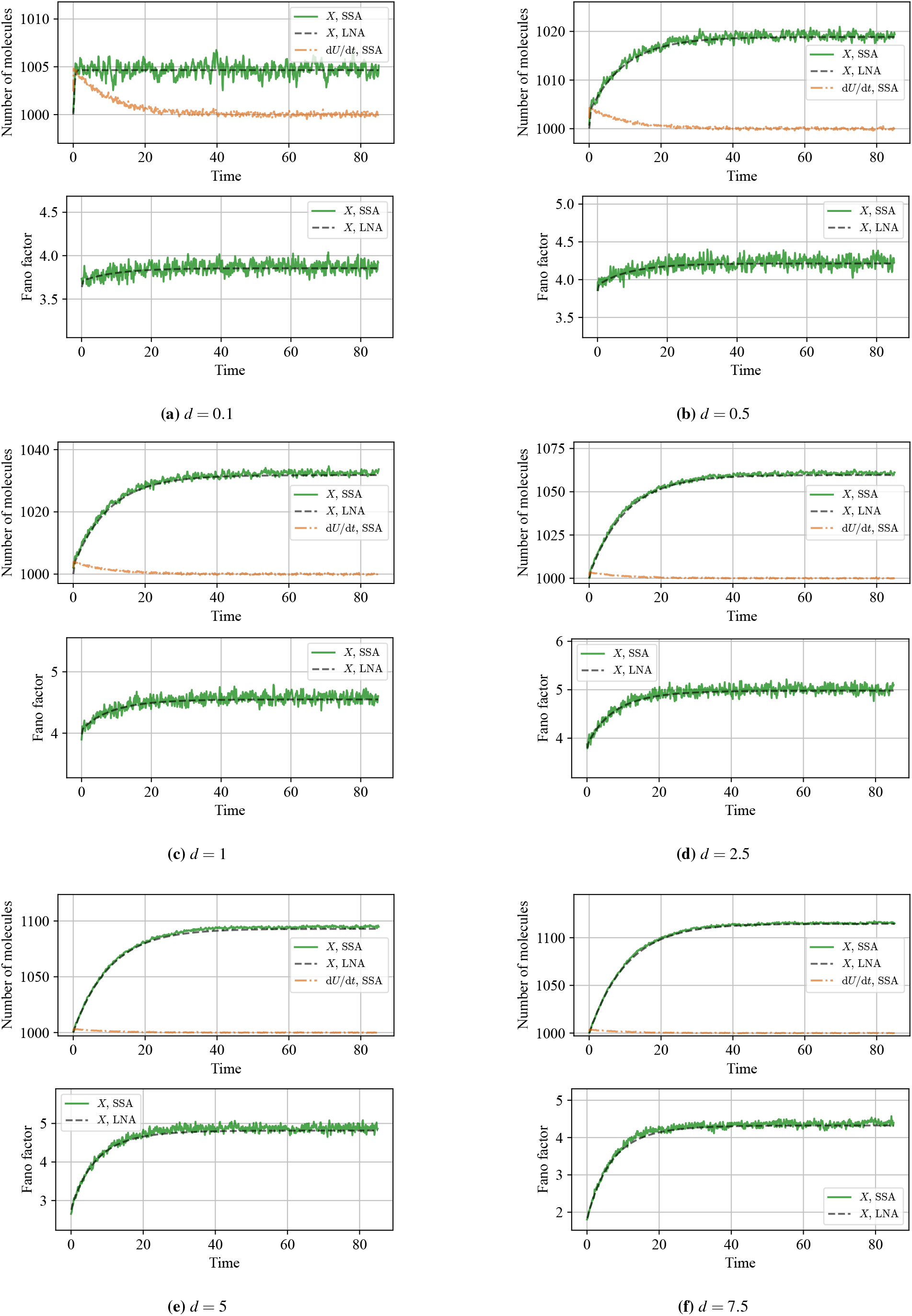
*AC-BioSD-II* - Effect of degradation of all species under input stimulation. The simulation scenario of Fig. 2(g,h) is repeated for different values of the common degradation rate constant *d*. Individual stochastic trajectories are presented in Fig. S8 of the Supplementary Material.

Finally, in the computational analysis presented in the figures, the parameter *d* is varied over a range for which all of the above cases yield stable dynamics.

### 3.2 Effects of slow annihilation reactions

As illustrated in Figs. 9-10 and S9-S10, when the annihilation reaction between *Z*_1_ and *Z*_2_ slows down (i.e., as *n* decreases), the output noise levels decrease. At the same time, the accuracy of signal differentiation gradually deteriorates, a trend that becomes evident for sufficiently slow reactions. This observation underscores an important performance trade-off that should be considered in experimental implementations.

**Figure 9:**
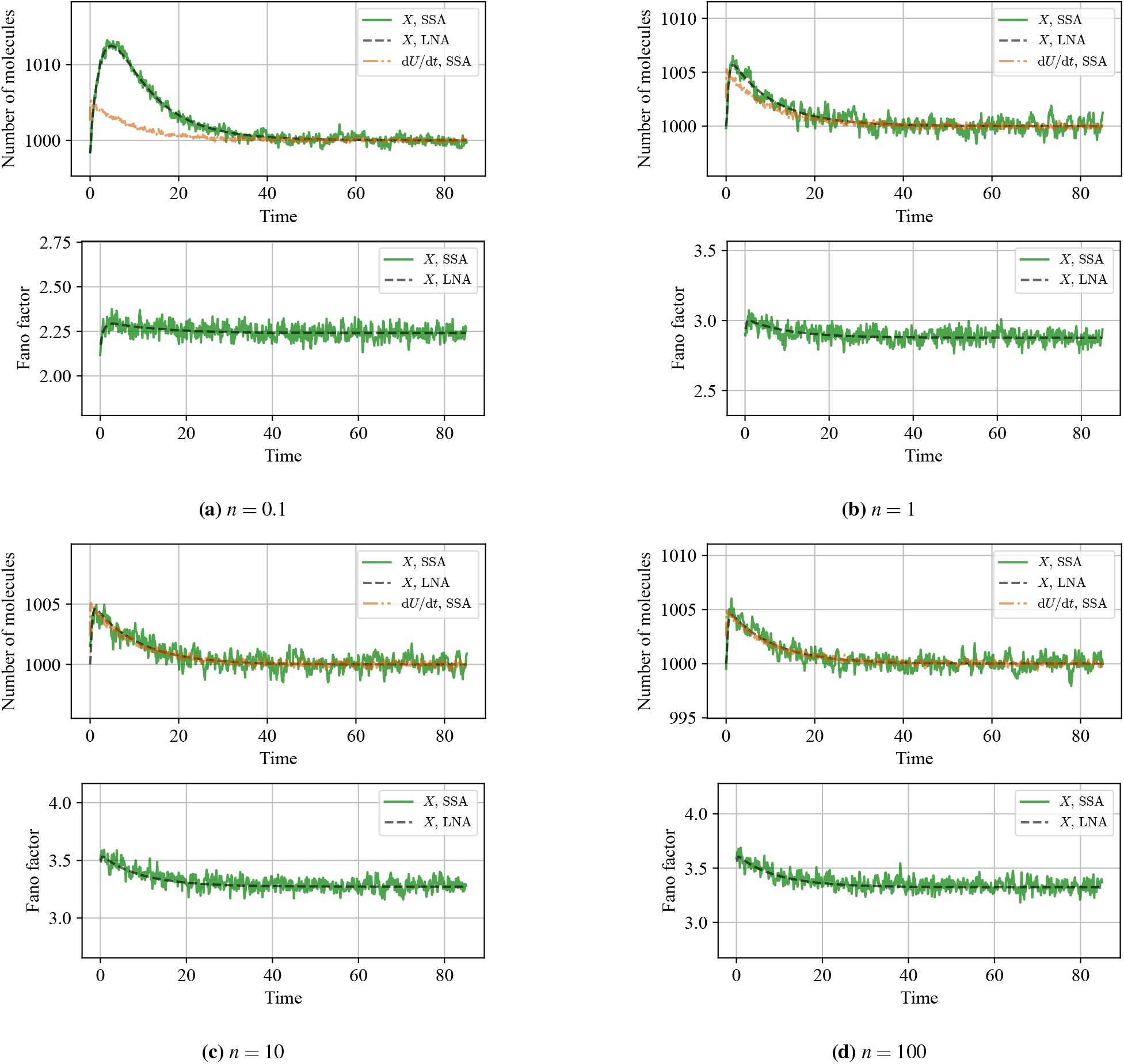
*BioSD-II*: Effect of slow annihilation of species *Z*_1_ and *Z*_2_. The simulation scenario of Fig. 2(e,f) is repeated for different values of the annihilation rate constant *n* (*n <* 1000). Individual stochastic trajectories are presented in Fig. S9 of the Supplementary Material.

**Figure 10:**
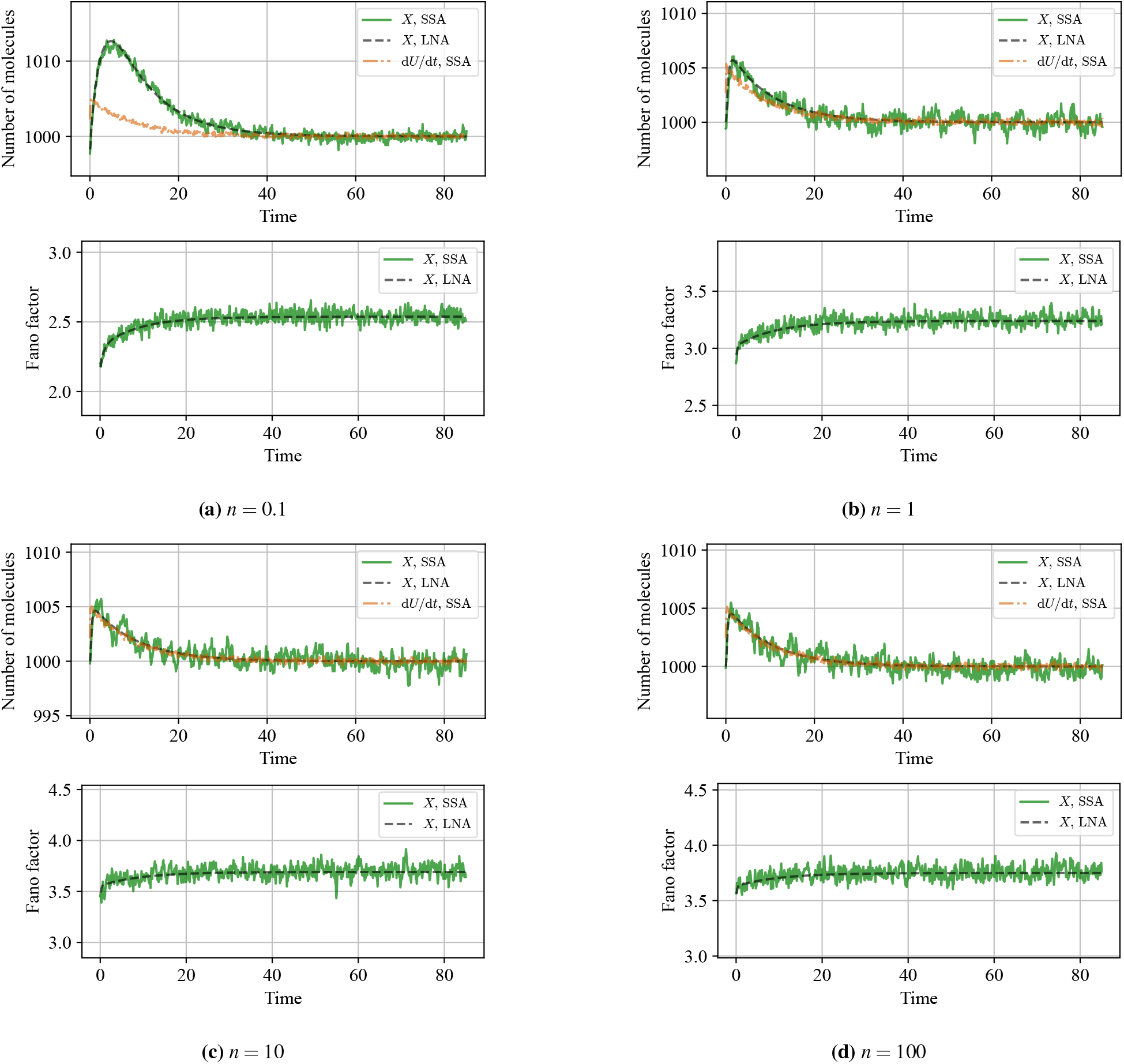
*AC-BioSD-II*: Effect of slow annihilation of species *Z*_1_ and *Z*_2_. The simulation scenario of Fig. 2(g,h) is repeated for different values of the annihilation rate constant *n* (*n <* 1000). Individual stochastic trajectories are presented in Fig. S10 of the Supplementary Material.

## 4 Derivative control through negative feedback

### 4.1 *BioSD-II*-based control

Fig. 11a illustrates an abstract biomolecular process to be controlled (open-loop process), represented as a cloud network. The process includes a target species (the output species of interest), *Y*_*t*_, which interacts with *BioSD*-*II* through a negative feedback loop. This interaction gives rise to a *BioSD*-*II*-based controller defined by the CRN:

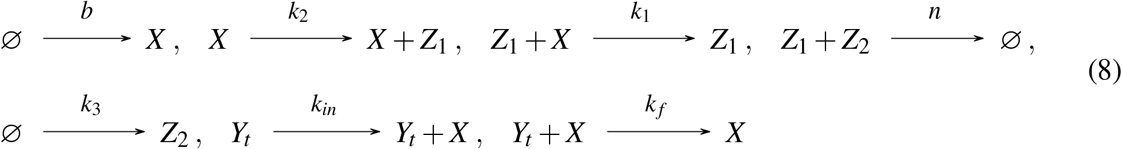

where *Y*_*t*_ is allowed to participate in an arbitrary number of reactions within the cloud network.

**Figure 11:**
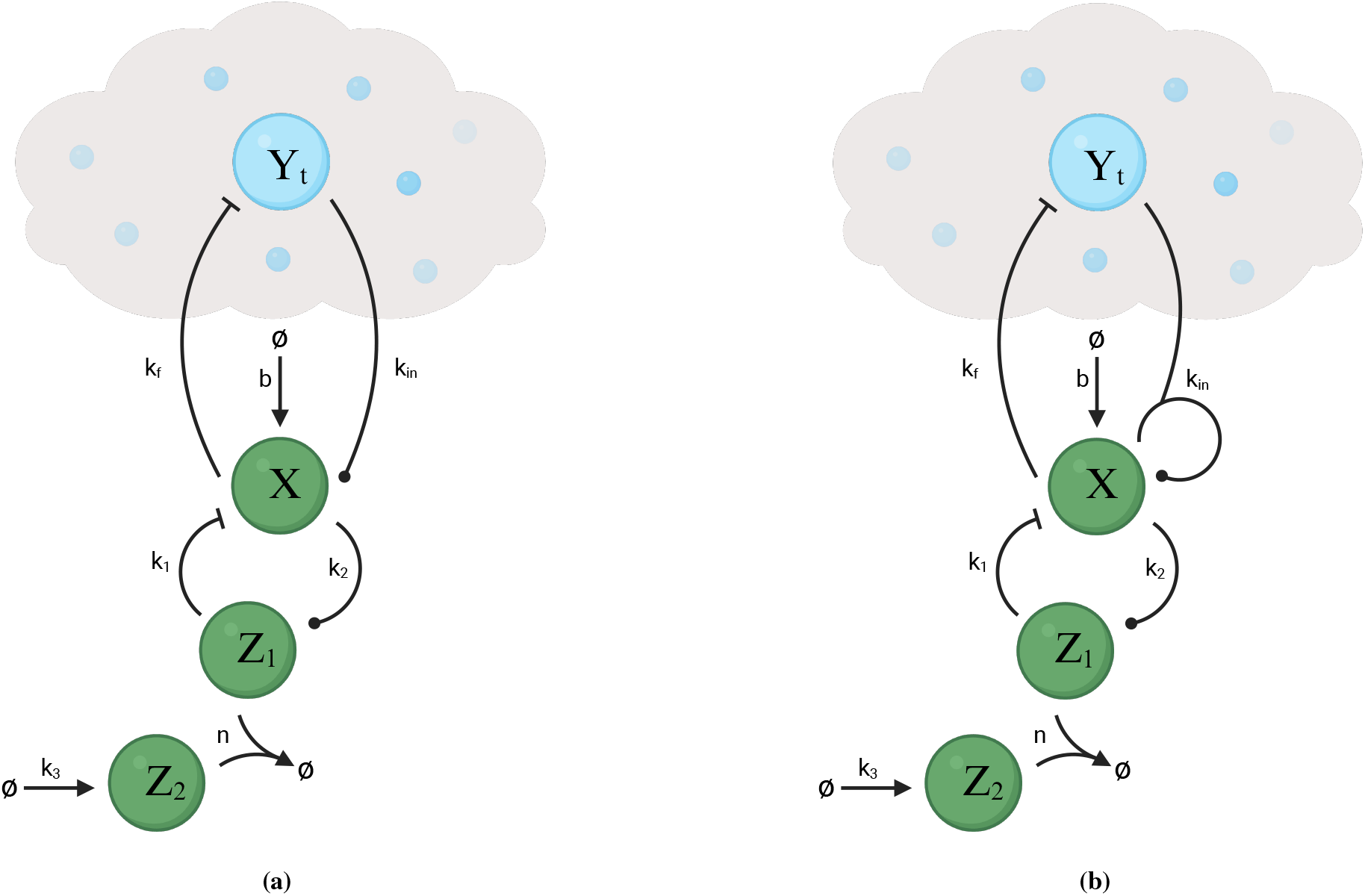
Biomolecular topologies exhibiting negative-feedback, derivative-control action. Schematic of (a) a *BioSD*-*II*-based controller; (b) an *AC*-*BioSD*-*II*-based controller. The required measurement and actuation reactions are implemented via the same species of the process being controlled

Focusing on the corresponding RREs, we can show (see Section S7 of the Supplementary Material) that, near an equilibrium of interest of the resulting closed-loop topology, the CRN in Eq. (8) realizes a control law expressed in the Laplace domain as:

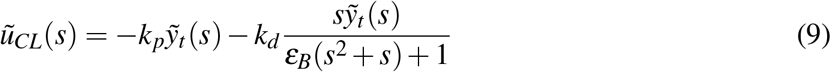

with 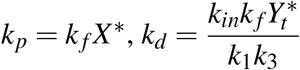 and 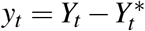.

Eq. (9) represents a Proportional-Derivative (PD) negative feedback control strategy, commonly used in technological control applications [50]. Note that, in the more general case (see Section S7 of the Supplementary Material), where the feedback control action is applied on a different species of the cloud network (interacting with *Y*_*t*_), the proportional control component vanishes, resulting in pure (filtered) derivative control. In addition, we prove that local asymptotic stability of the afore-mentioned equilibrium point can be guaranteed for any cloud network (locally) corresponding to a stabilizable and detectable process whose input–output relation is given by a weakly strictly positive real (WSPR) transfer function (the associated definitions and proofs can be found in Section S7 of the Supplementary Material). This constitutes a powerful result, as it provides sufficient conditions for achieving (local) closed-loop stability based solely on the (linearized) process to be controlled.

We now consider the scenario where the abstract process to be controlled consists of a general bursty birth-death process, i.e. the former coincides with the CRN given by Eq. (3) where *U* is replaced by *Y*_*t*_. The corresponding transfer function (here we have minimal realization) around the resulting closed-loop steady state is WSPR and, thus, the latter is locally asymptotically stable (see Section S7 of the Supplementary Material).

Assuming that the parameter values of the open-loop process are known and, for simplicity, set to unity, in the limit as *n* → ∞, the Fano factor of the output, *Y*_*t*_, at steady state becomes:

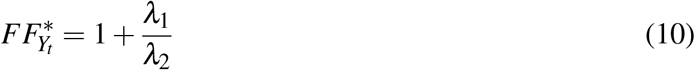

where

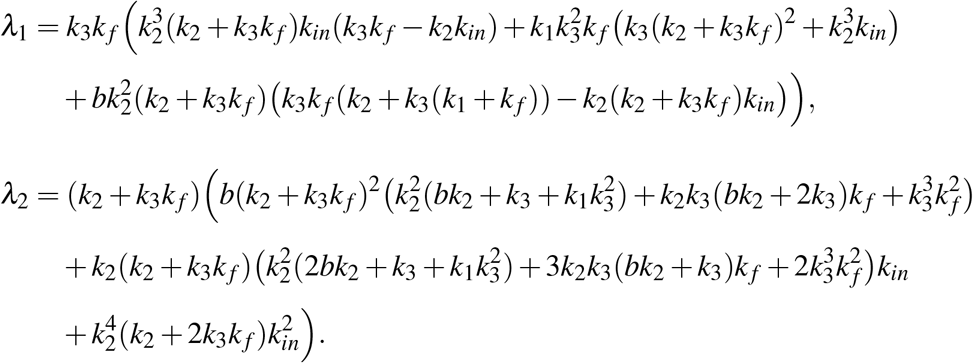

Fixing the parameter values of the open-loop process allows for a substantial simplification of the expression for 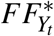, thereby highlighting the influence of the controller-associated parameters. The corresponding expression for arbitrary open-loop parameters, as well as an ODE system for 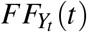, can be found in Section S7 of the Supplementary Material. Interestingly, the *BioSD*-*II*-based negative feedback action can lead to sub-Poisson steady-state output noise - that is, 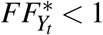 for *λ*_1_ *<* 0 (*λ*_2_ *>* 0 always).

### 4.2 *AC-BioSD-II*-based control

Fig. 11b illustrates the same concept as Fig. 6(a) with the difference that the *BioSD*-*II* has now been replaced by *AC*-*BioSD*-*II*. The resulting *AC*-*BioSD*-*II*-based controller is defined by the CRN:

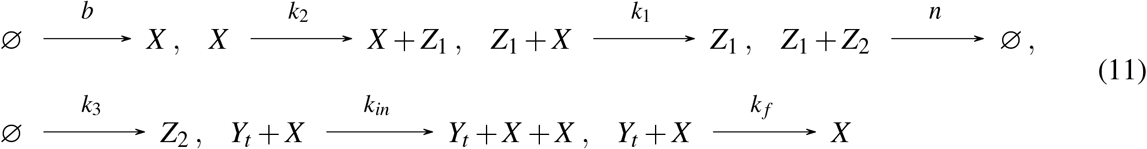

which, similarly to the *BioSD*-*II*–based controller, locally realizes a PD control law expressed in the Laplace domain as

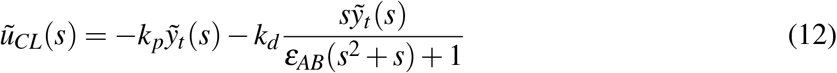

with 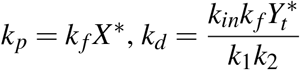 and 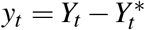.

As before, the proportional control term in Eq. (12) vanishes if the feedback control action is exerted on a different species within the cloud network that interacts with *Y*_*t*_. A detailed analysis on the above is provided in Section S8 of the Supplementary Material. There, we also show that the sufficient conditions for local closed-loop asymptotic stability presented in Subsection 4.1 remain valid for this topology, as well. This entails that, if we replace the abstract process to be controlled with a general bursty birth-death process (see Subsection 4.1), the resulting closed-loop steady state is locally asymptotically stable.

Finally, as in the previous case, we calculate the steady-state output Fano factor for unit open-loop process parameter values and *n* → ∞, as:

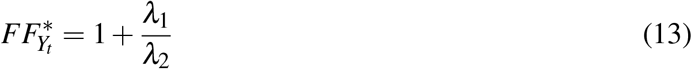

where

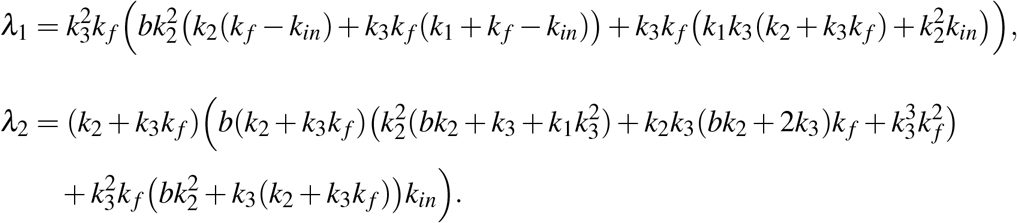

Note that 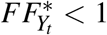 for *λ*_1_ *<* 0 (*λ*_2_ *>* 0 always) and, thus, the *AC*-*BioSD*-*II*-based negative feedback action is able to drive the output noise below Poisson levels. The expression for 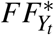 assuming arbitrary parameters, along with an ODE system for 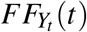 can be found in Section S8 of the Supplementary Material.

Figs. 13 and S13 present a simulation scenario using the LNA, validated against SSA. The setup is based on the controllers described in Sections 4.1 and 4.2, with the target species participating in a bursty birth-death process (assuming unit parameter values). In both cases, the closed-loop dynamics are stable, achieving sub-Poisson output noise at steady state. In parallel, this reflects a mitigation of output stochastic fluctuations, given that the corresponding noise of the open-loop process is Poisson.

**Figure 12:**
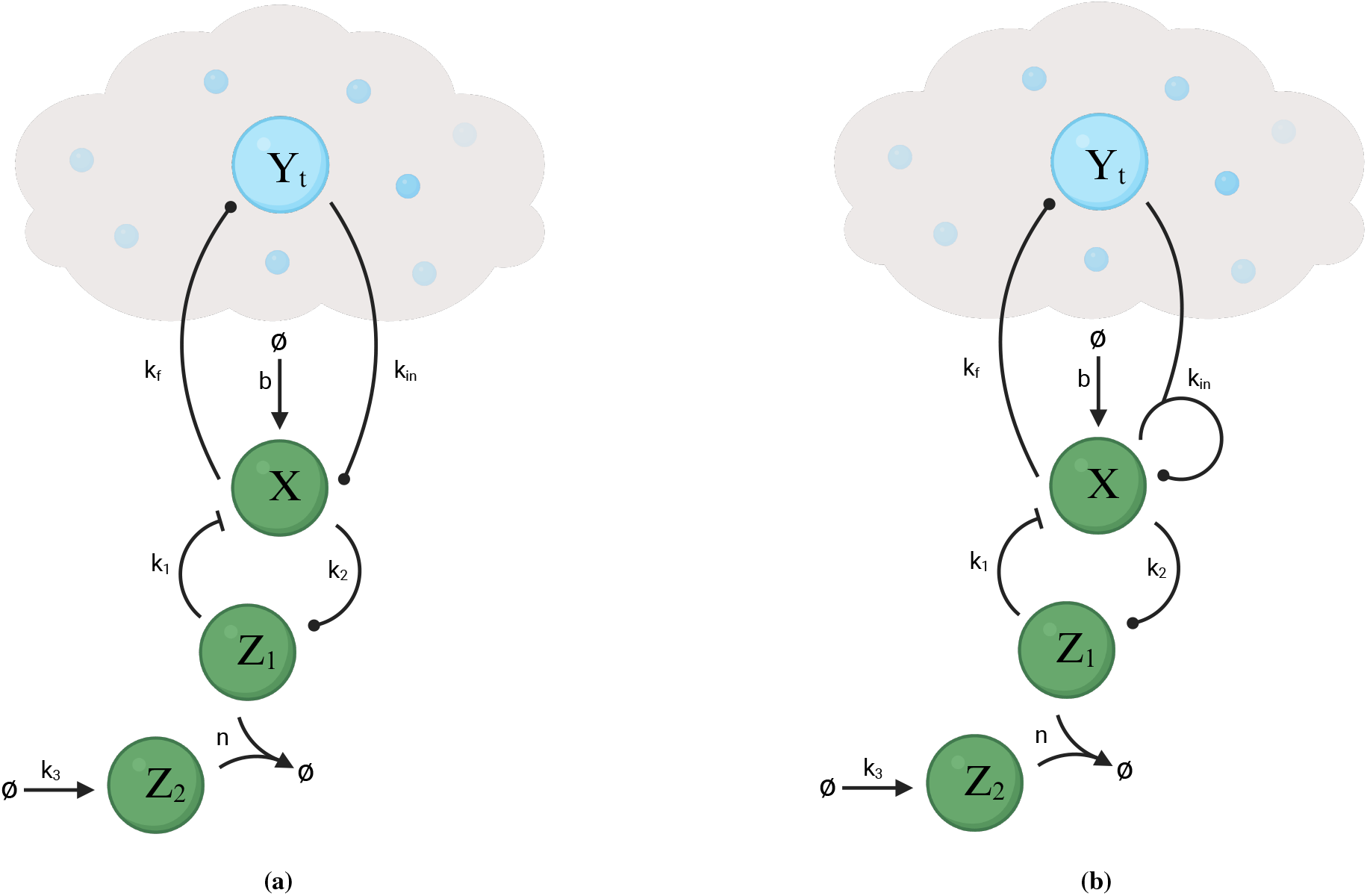
Biomolecular topologies exhibiting positive-feedback, derivative-control action. Schematic of (a) a *BioSD*-*II*-based controller; (b) an *AC*-*BioSD*-*II*-based controller. The required measurement and actuation reactions are implemented via the same species of the process being controlled.

**Figure 13:**
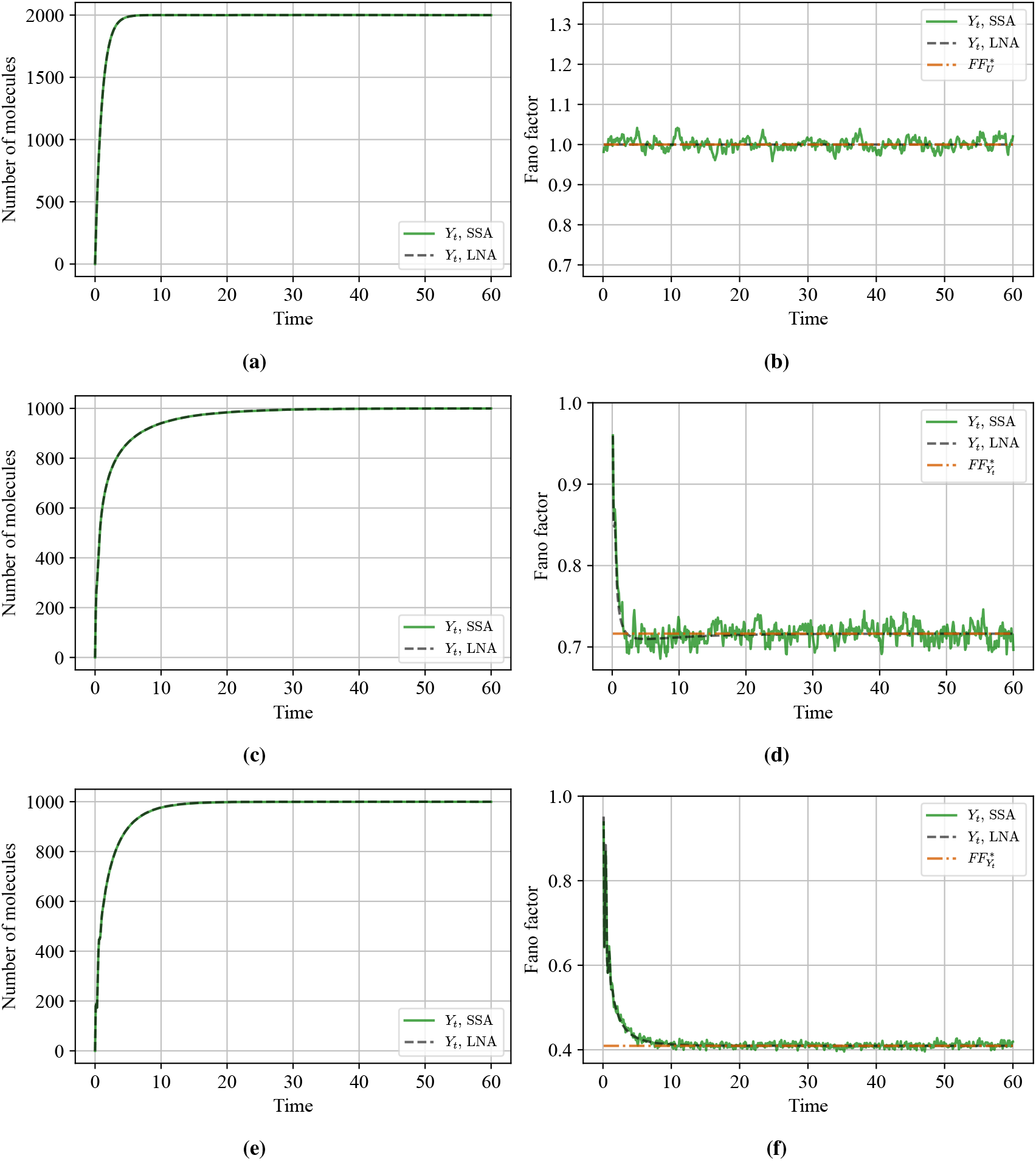
Negative-feedback, derivative control. Time evolution of the mean molecular copy number and the Fano factor (together with the corresponding analytically derived steady-state approximation), obtained using SSA and LNA for: (a,b) a target species (*Y*_*t*_) participating in a general bursty birth–death biomolecular process (CRN in Eq. (3)) with *c* = 1, *p*_1_ = 1, and *p*_2_ = 1 (process to be controlled); (c,d) the target species in (a,b) regulated by a *BioSD*-*II*-based controller (CRNs in Eq. (8)) with *k* _*f*_ = 0.5, *b* = 25, *k*_1_ = 1, *k*_2_ = 10, *k*_3_ = 20, *n* = 1000 and *k*_*in*_ = 400. (e,f) the target species in (a,b) regulated by an *AC*-*BioSD*-*II*-based controller (CRNs in Eq. (11)) with *k*_*in*_ = 200 and all remaining parameters unchanged. The initial conditions in all cases are set to zero. Individual stochastic trajectories for the above simulation scenarios are presented in Fig. S13 of the Supplementary Material.

## 5 Derivative control through positive feedback

### 5.1 *BioSD-II*-based control

Fig. 12a illustrates a positive feedback interconnection of *BioSD*-*II* with a cloud network. The corresponding *BioSD*-*II*-based controller is described by the CRN:

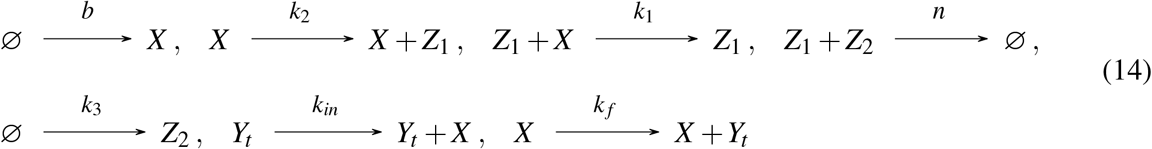

Around a closed-loop system equilibrium of interest, the resulting RRE-based control law can be expressed in the Laplace domain as (see Section S9 of the Supplementary Material):

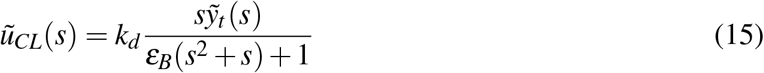

With 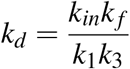, which represents a positive feedback (filtered) derivative control scheme. Note that Eq. (15) remains the same regardless of the cloud-network species on which the feedback action is applied. Subsequently, if the abstract process to be controlled is replaced with a bursty birth-death process, as in Section 4, we can show that closed-loop asymptotic stability is guaranteed for any chosen parameters (see Section S9 of the Supplementary Material).

Assuming now unit parameter values for the bursty birth-death process and *n* → ∞ (as in Section 4), we obtain the steady-state output Fano factor:

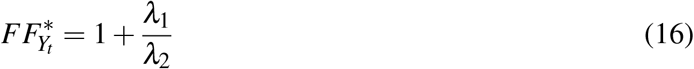

where

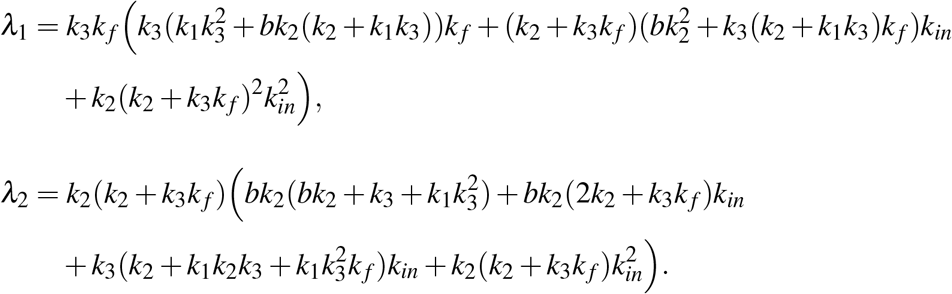

In Section S9 of the Supplementary Material, we provide an expression for 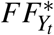 (in the same form as Eq. (16)) assuming arbitrary parameters as well as an ODE system for 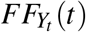. As can be seen there, *λ*_1_, *λ*_2_ *>* 0 for any chosen parameters and, thus, the steady-state output noise is always super-Poisson 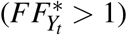.

### 5.2 *AC-BioSD-II*-based control

Fig. 12b illustrates the same concept as Fig. 8(a), except that *BioSD*-*II* has now been substituted with *AC*-*BioSD*-*II*. The resulting *AC*-*BioSD*-*II*-based controller is defined by the CRN:

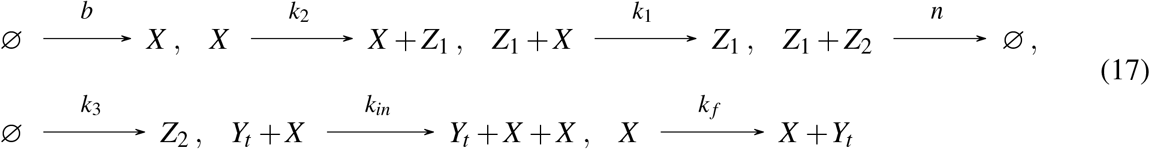

Similarly to the corresponding *BioSD*-II–based control configuration, this topology implements a positive-feedback (filtered) derivative control law, expressed in the Laplace domain as (see Section S10 of the Supplementary Material):

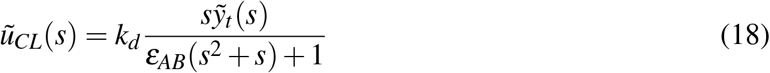

With 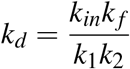. As before, this expression remains unchanged in the more general case where the feedback control action is applied to a cloud-network species different from the output one. If we now consider a bursty birth–death process to be controlled (see Section 4), then, contrary to the *BioSD*-II–based control configuration, closed-loop asymptotic stability is not always guaranteed; a discussion on identifying parameter regimes that do ensure such stability is provided in Section S10 of the Supplementary Material.

By setting all parameters of the bursty birth–death process to unity and taking the limit *n* → ∞ (as in Section 4), the steady-state output Fano factor can be computed as:

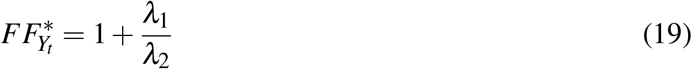

where

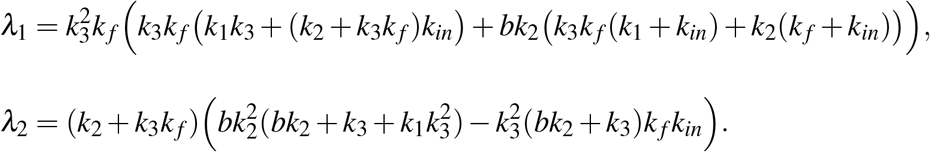

An expression for 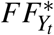 (in the same form as Eq. (19)), together with an ODE system for 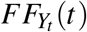 under arbitrary parameter values, is provided in Section S10 of the Supplementary Material. As demonstrated therein, *λ*_1_, *λ*_2_ *>* 0 for any realistic parameter set of interest and, thus, the steady-state output noise is always above Poisson levels 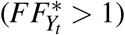.

In Figs. 14 and S14, we present a simulation scenario based on the control architectures described in Sections 5.1 and 5.2, assuming a bursty birth–death process as the open-loop process with unit parameter values (as in Fig. 7). Results from both the LNA and SSA are depicted. Note that here we show an instance in which the steady-state output noise levels of the closed-loop topology exceed those of the open-loop process—an outcome that, as demonstrated in Figs. S15, S16, does not always occur.

**Figure 14:**
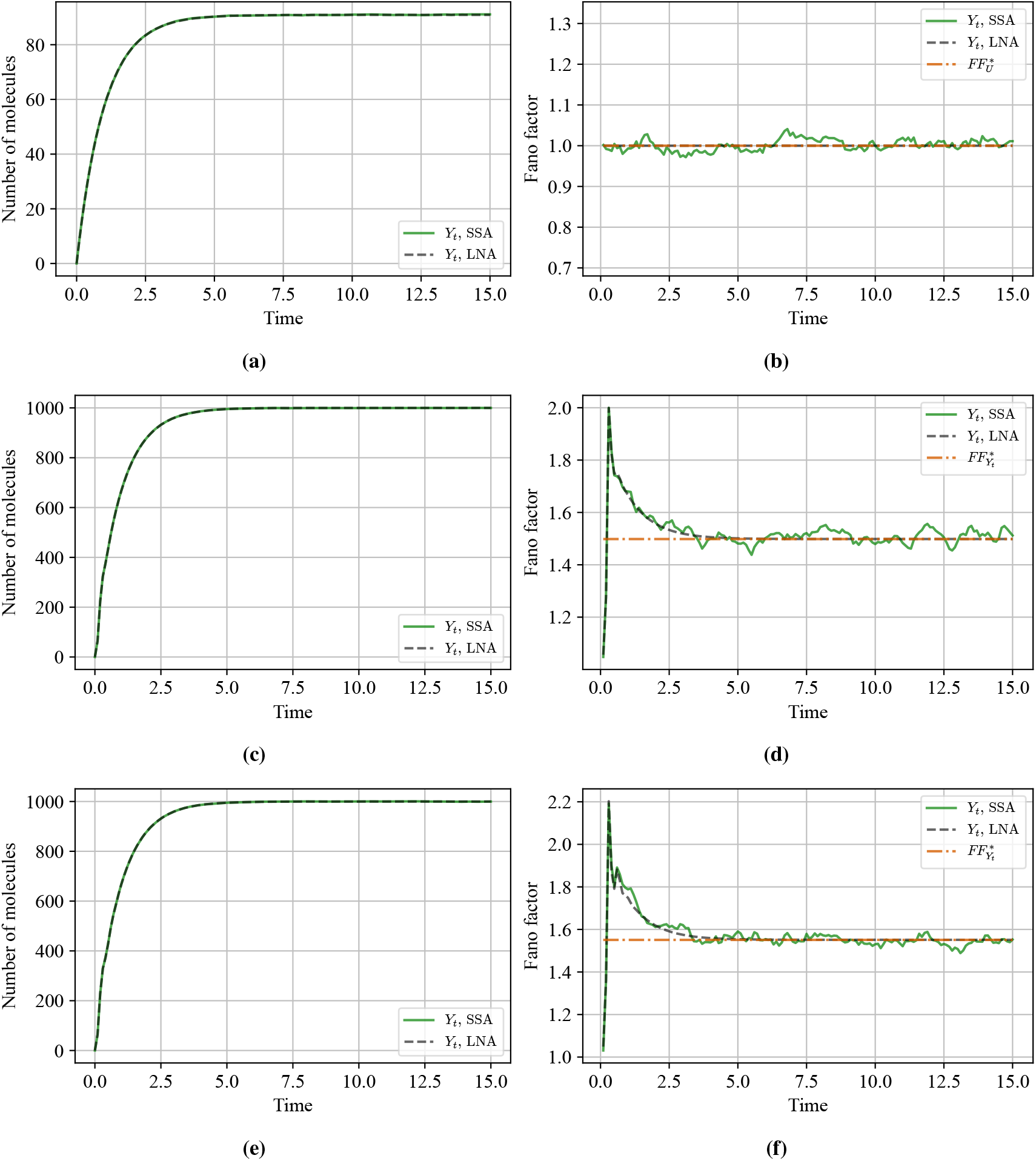
Positive-feedback, derivative control. Time evolution of the mean molecular copy number and the Fano factor (together with the corresponding analytically derived steady-state approximation), obtained using SSA and LNA for: (a,b) a target species (*Y*_*t*_) participating in a general bursty birth–death biomolecular process (CRN in Eq. (3)) with *c* = 1, *p*_1_ = 1, and *p*_2_ = 1 (process to be controlled); (c,d) the target species in (a,b) regulated by a *BioSD*-*II*-based controller (CRNs in Eq. (14)) with *k* _*f*_ = 0.5, *b* = 250, *k*_1_ = 16, *k*_2_ = 1, *k*_3_ = 20, *n* = 1000 and *k*_*in*_ = 32. (e,f) the target species in (a,b) regulated by an *AC*-*BioSD*-*II*-based controller (CRNs in Eq. (17)) with *k*_*in*_ = 1.6 and all remaining parameters unchanged. The initial conditions in all cases are set to zero. Individual stochastic trajectories for the above simulation scenarios are presented in Fig. S14 of the Supplementary Material.

## 6 Discussion

In this work, we focus on the noise analysis of the *BioSD*-*II* module, initially introduced in [7], and of the *AC*-*BioSD*-*II* module, its newly proposed structural variant. Both topologies are capable of exhibiting filtered derivative action with respect to an input signal, which is applied through a catalytic production process in the former and an autocatalytic production process in the latter. As a test input signal, we consider a biomolecular species participating in a general bursty birth–death process under both nominal and non-ideal conditions while we also take into account the special case of no input stimulus. In addition, assuming abstract processes to be controlled, we employ these modules to construct several regulatory topologies based on negative and positive feedback control action. To gain more practical insight into their function, we further evaluate their regulatory capacity with respect to the aforementioned bursty birth–death process.

For each scenario under investigation, we examine key characteristics of the macroscopic system behaviour. In particular, for the standalone *BioSD*-*II* and *AC*-*BioSD*-*II* modules, we focus on their ability to differentiate signals, accompanied by low-pass filtering action, and maintain system stability. For the regulatory topologies, we investigate the form of the feedback control law they realize— various forms of (filtered) derivative control which, at times, are combined with proportional control action. We additionally assess closed-loop stability and, where applicable, establish sufficient stability conditions expressed solely in terms of the properties of the controlled process around relevant equilibria.

Within the macroscopic settings of interest (based on the features mentioned above), we perform noise analysis using LNA. This allows us to generate a closed set of ODEs for each case, describing the time evolution of the mean vector and the covariance matrix of the stochastic fluctuations around the corresponding deterministic solutions. As a noise metric, we use the Fano factor (variance normalized to the mean), which can be computed directly from these equations. In most cases, we are able to derive analytical expressions for the steady-state Fano factor of the system output (while, under some additional assumptions, we discuss a simple method of obtaining analytical approximations of the transient dynamics). We further characterize noise levels by comparing them with Poisson noise, for which the value of the Fano factor is equal to one. In parallel, we employ SSA to computationally assess the accuracy of our LNA-based results.

We believe that our work contributes to a better understanding of the noise profile associated with derivative-action–based biomolecular topologies. In many technological engineering applications, the operation of signal differentiators has been linked to disruptive noise amplification. Consequently, analyzing the behaviour of their biomolecular counterparts in the presence of (intrinsic and extrinsic) stochastic noise is crucial for guiding potential experimental efforts. Our results suggest that the *BioSD*-*II* and *AC*-*BioSD*-*II* modules can operate reliably in noisy environments. Specifically, when considered in isolation, their output mean varies approximately proportionally with the input mean around an equilibrium of interest, provided that the latter evolves sufficiently slowly over time. Although the steady-state output noise is always super-Poisson, appropriate parameter tuning may reduce it below the corresponding input noise level. We further show that introducing an additional degradation mechanism acting solely on the output species leaves this behaviour practically unaffected, in contrast to the case where such degradation impacts all species within the *BioSD* topologies. Moreover, we observe that decreasing the common annihilation rate between the two auxiliary species leads to lower output noise levels, albeit with a reduction in the accuracy of signal differentiation. When embedded within feedback control architectures and under suitable tuning, both *BioSD* modules might be able to reduce the steady-state output noise of the regulated process relative to the unregulated case. Interestingly, negative feedback can suppress noise to sub-Poisson levels, whereas under positive feedback the noise remains strictly super-Poisson. Note that these conclusions, though insightful, are largely based on specific test input signals and controlled processes, and thus no general performance guarantees can be provided.

Finally, although the LNA is a convenient approach for studying the behaviour (up to second-order statistical moments) of biomolecular systems in the stochastic setting, it provides only approximate descriptions whose accuracy is not always guaranteed, except for certain classes of systems [32, 51]. For example, in regimes with sufficiently low molecular counts, the LNA-based analysis presented herein may not yield a realistic description. In such cases, more suitable methods—such as (tailored) moment-closure techniques [52, 53]—should be employed to obtain analytical results. Exploring these alternatives constitutes an interesting direction for future work.

## Code availability

The programming codes supporting this work can be found at: https://github.com/sespirio/noise_paper_sim.

## Authors’ contributions

Conceptualization and methodology: E.A.; Formal analysis and numerical simulations: E.A, S.E.R., L.L., C.W.R., Visualization: E.A., S.E.R. ; Writing: E.A., S.E.R, L.L., L.C., I.G.K., C.W.R., J.L.A.; Supervision: L.L., L.C., I.G.K., C.W.R., J.L.A. ; Funding: J.L.A.

## Acknowledgements

Figures 1, 11, 12, S11, and S12 were generated using BioRender.com.

