## Supplementary Material for "Noise analysis of derivative-action biomolecular topologies"

Emmanouil Alexis[1,2,*], Sebastián Espinel-Ríos[3], Luca Laurenti[4,5], Luca Cardelli[6], Ioannis G. Kevrekidis[7,8], Clarence W. Rowley[9], and José L. Avalos[1,2,10,11,12]

[1]*Omenn-Darling Bioengineering Institute, Princeton University, New Jersey, USA*

[2]*Department of Chemical and Biological Engineering, Princeton University, New Jersey, USA*

[3]*School of Chemical and Bioprocess Engineering, University College Dublin, Dublin, Ireland*

[4]*Delft Center for Systems and Control, Delft University of Technology, Delft, Netherlands*

[5]*The Italian Institute of Artificial Intelligence (AI4I), Turin, Italy*

[6]*Department of Computer Science, University of Oxford, Oxford, UK*

[7]*Department of Chemical and Biomolecular Engineering, Johns Hopkins University, Baltimore, USA*

[8]*Department of Applied Mathematics and Statistics, Johns Hopkins University, Baltimore, USA*

[9]*Department of Mechanical and Aerospace Engineering, Princeton University, New Jersey, USA*

[10]*High Meadows Environmental Institute, Princeton University, New Jersey, USA*

[11]*The Andlinger Center for Energy and the Environment, Princeton University, New Jersey, USA*

[12]*Department of Molecular Biology, Princeton University, New Jersey, USA*

**S2 Numerical Simulations** **7**

**S3 *BioSD-II* - Fano factor** **8**

**S4 *AC-BioSD-II*** **10**

**S11 Supplementary figures** **48**

**References** **61**

### S1 Modeling principles

#### S1.1 Chemical Reaction Networks

A Chemical Reaction Network (CRN) is defined as a pair of finite sets $\mathcal{C} = (\mathcal{L}, \mathcal{R})$, where $\mathcal{L}$ denotes the set of chemical species with size $|\mathcal{L}|$ and $\mathcal{R}$ denotes the set of reactions. A reaction $\tau \in \mathcal{R}$ is represented by a triple $\tau = (r_\tau, p_\tau, k_\tau)$, with $r_\tau, p_\tau \in \mathbb{N}^{\mathcal{L}}$ denoting the stoichiometry of reactants and products, respectively, and $k_\tau \in \mathbb{R}_+$ the reaction rate coefficient. Moreover, the net change corresponding to a reaction $\tau$ is defined as $u_\tau = p_\tau - r_\tau$. Consider, for instance, a reaction $\tau_a = ([1,0,1]^T, [0,2,0]^T, k_a)$ with $\mathcal{L} = (X,Y,Z)$ which yields $u_{\tau_a} = [-1,-2,-1]^T$ (where $^T$ denotes the transpose of a matrix). This can be also expressed in the conventional form: $\tau_a$: $X + Y \xrightarrow{k_a} 2Z$.

#### S1.2 Reaction Rate Equations

The time evolution of a given CRN $\mathcal{C} = (\mathcal{L}, \mathcal{R})$ in the thermodynamic (large-system) limit can be described by the solution of the following set of Ordinary Differential Equation (ODEs), also known as Reaction Rate Equations (RREs):

$$\frac{d\Phi(t)}{dt} = F(\Phi(t)) \tag{S1}$$

where $F(\Phi(t)) = \sum_{\tau \in \mathcal{R}} u_\tau \alpha_{c,\tau}(\Phi(t))$ and $\alpha_{c,\tau}(\Phi(t)) = k_\tau \prod_{i=1}^{|\mathcal{L}|} \Phi_i(t)^{r_{\tau,i}}$. $\Phi(t) \in \mathbb{R}^{|\mathcal{L}|}$ represents the vector of the concentrations of all species at time $t$. Note that here, we are concerned with mass-action kinetics.

#### S1.3 Linear Noise Approximation

In the stochastic setting, a CRN can be modeled as a continuous-time discrete-space Markov process (CTMC) $(\Theta(t), t \geq 0)$, whose time evolution is described by the Chemical Master Equation (CME). The Linear Noise Approximation (LNA) provides a continuous state-space approximation of this CTMC based on a Gaussian process - it can be obtained by truncating the solution of the CME at the second moment. More specifically, for a CRN $\mathcal{C} = (\mathcal{L}, \mathcal{R})$, the probability distribution of $\Theta(t)$ at time $t$ can be approximated under the LNA by the probability distribution of a random vector $\Theta_a(t)$, such that

$$\Theta(t) \approx \Theta_a(t) = \Omega\Phi(t) + \sqrt{\Omega}G(t) \tag{S2}$$

where $\Omega = V^{\Omega}N_A$ denotes the volumetric factor with $V^{\Omega}$ and $N_A$ representing the system volume and Avogadro's number, respectively; $\Phi(t)$ denotes the solution of the RREs; and $G(t) = (G_1(t), G_2(t), \ldots, G_{|\mathcal{L}|}(t))$ is a random vector independent of $\Omega$, capturing the stochastic fluctuations at time $t$. The probability distribution of $G(t)$ can be determined by the solution of a linear Fokker-Plank equation. Therefore, at each time instant $t$, $G(t)$ has a multivariate normal distribution, with expected value ($E[G(t)]$) and covariance matrix ($C[G(t)]$) given by the following ODEs:

$$\frac{dE[G(t)]}{dt} = J_F(\Phi(t))E[G(t)] \tag{S3}$$

$$\frac{dC[G(t)]}{dt} = J_F(\Phi(t))C[G(t)] + C[G(t)]J_F^T(\Phi(t)) + W(\Phi(t)) \tag{S4}$$

where $J_F(\Phi(t))$ represents the Jacobian of $F(\Phi(t))$ and $W(\Phi(t)) = \sum_{\tau \in \mathcal{R}} u_\tau u_\tau^T \alpha_{c,\tau}(\Phi(t))$.

Note that, assuming zero initial conditions, Eq. (S3) yields $E[G(t)] = 0$ for all $t$. Moreover, the following relations hold:

$$E[\Theta(t)] \approx E[\Theta_a(t)] = \Omega\Phi(t)$$

$$C[\Theta(t)] \approx C[\Theta_a(t)] = \Omega C[G(t)]$$

Note that although the LNA is only exact in the large-population limit (i.e., for $\Omega \to \infty$), it generally provides accurate approximations even at relatively small molecule numbers [1]. In practice, when the species of interest exhibit unimodal distributions and molecular abundances allow for a continuous description, the LNA is generally found to be reliable.

For a more comprehensive discussion of the above concepts, the reader is referred to [2].

In what follows, we adopt a simplified notation to streamline the analysis. Specifically, we denote the components of $\Phi(t)$ by the corresponding species names, and the components of $C[G(t)]$ by $V_q$, where $q$ denotes either a single species (for variances) or a pair of species (for covariances). For example, given a CRN with two species $(X, Y)$, we will have:

$$\Phi(t) = \begin{bmatrix} X \\ Y \end{bmatrix}$$

$$C[G(t)] = \begin{bmatrix} V_X & V_{XY} \\ V_{XY} & V_Y \end{bmatrix}$$

Finally, the steady state of a variable is denoted by $^*$; for example, the steady states of $X$ and $Y$ are $X^*$ and $Y^*$, respectively. Note that here we are interested in hyperbolic steady states–the corresponding system's Jacobian has no eigenvalues on the imaginary axis.

### S2 Numerical Simulations

In all stochastic simulations, we used Gillespie's stochastic simulation algorithm (SSA) [3]. Specifically, we generated 10,000 stochastic trajectories per case and computed the mean, variance, and Fano factor (defined as the ratio of variance to mean). Each trajectory was simulated in Python using the BasiCO library [4], which interfaces with COPASI (COmplex PAthway SImulator) [5]. To convert concentrations to particle numbers, we used $n_i = V_c c_i N_A$, where $n_i$ is the number of particles of species $i$, $c_i$ is the concentration of species $i$, $V_c$ is the compartment volume, and $N_A$ is Avogadro's constant. LNA trajectories were generated by numerically integrating the corresponding system of ODEs in Python using SciPy [6], employing the LSODA solver. For both stochastic and LNA simulations, we used an output sampling interval of 0.1.

#### S3 *BioSD-II* - Fano factor

The *BioSD-II* topology is given by the CRN in Eq. (1).

Assuming $U = 0$, Eqs. (S1), (S4) yield:

$$\begin{aligned}
\frac{dX}{dt} &= b - k_1 X Z_1 \\
\frac{dZ_1}{dt} &= k_2 X - n Z_1 Z_2 \\
\frac{dZ_2}{dt} &= k_3 - n Z_1 Z_2 \\
\frac{dV_X}{dt} &= b + k_1 X Z_1 - 2k_1 X V_{XZ_1} - 2k_1 Z_1 V_X \\
\frac{dV_{XZ_1}}{dt} &= k_2 V_X - k_1 X V_{Z_1} - k_1 Z_1 V_{XZ_1} - n Z_1 V_{XZ_2} - n Z_2 V_{XZ_1} \\
\frac{dV_{XZ_2}}{dt} &= -k_1 X V_{Z_1 Z_2} - k_1 Z_1 V_{XZ_2} - n Z_1 V_{XZ_2} - n Z_2 V_{XZ_1} \\
\frac{dV_{Z_1}}{dt} &= k_2 X + 2k_2 V_{XZ_1} + n Z_1 Z_2 - 2n Z_2 V_{Z_1} - 2n Z_1 V_{Z_1 Z_2} \\
\frac{dV_{Z_1 Z_2}}{dt} &= k_2 V_{XZ_2} + n Z_1 Z_2 - n Z_1 V_{Z_2} - n Z_2 V_{Z_1} - n Z_1 V_{Z_1 Z_2} - n Z_2 V_{Z_1 Z_2} \\
\frac{dV_{Z_2}}{dt} &= k_3 + n Z_1 Z_2 - 2n Z_1 V_{Z_2} - 2n Z_2 V_{Z_1 Z_2}
\end{aligned}$$

For $E[X(t)] > 0$, the output Fano factor, $FF_X(t)$, is given as:

$$FF_X(t) = \frac{Var[X(t)]}{E[X(t)]} = \frac{V_X}{X} \tag{S5}$$

We now consider an input signal given by the CRN in Eq. (3). Taking into account Eqs. (1), (3), (S1), (S4), we get:

$$
\begin{aligned}
\frac{dU}{dt} &= cp_1 - p_2U \\
\frac{dX}{dt} &= b + k_{in}U - k_1XZ_1 \\
\frac{dZ_1}{dt} &= k_2X - nZ_1Z_2 \\
\frac{dZ_2}{dt} &= k_3 - nZ_1Z_2 \\
\frac{dV_U}{dt} &= p_1c^2 + p_2U - 2p_2V_U \\
\frac{dV_{UX}}{dt} &= k_{in}V_U - p_2V_{UX} - k_1XV_{UZ_1} - k_1Z_1V_{UX} \\
\frac{dV_{UZ_1}}{dt} &= k_2V_{UX} - p_2V_{UZ_1} - nZ_1V_{UZ_2} - nZ_2V_{UZ_1} \\
\frac{dV_{UZ_2}}{dt} &= -p_2V_{UZ_2} - nZ_1V_{UZ_2} - nZ_2V_{UZ_1} \\
\frac{dV_X}{dt} &= b + k_{in}U + 2k_{in}V_{UX} + k_1XZ_1 - 2k_1XV_{XZ_1} - 2k_1Z_1V_X \\
\frac{dV_{XZ_1}}{dt} &= k_2V_X + k_{in}V_{UZ_1} - k_1XV_{Z_1} - k_1Z_1V_{XZ_1} - nZ_1V_{XZ_2} - nZ_2V_{XZ_1} \\
\frac{dV_{XZ_2}}{dt} &= k_{in}V_{UZ_2} - k_1XV_{Z_1Z_2} - k_1Z_1V_{XZ_2} - nZ_1V_{XZ_2} - nZ_2V_{XZ_1} \\
\frac{dV_{Z_1}}{dt} &= k_2X + 2k_2V_{XZ_1} + nZ_1Z_2 - 2nZ_2V_{Z_1} - 2nZ_1V_{Z_1Z_2} \\
\frac{dV_{Z_1Z_2}}{dt} &= k_2V_{XZ_2} + nZ_1Z_2 - nZ_1V_{Z_2} - nZ_2V_{Z_1} - nZ_1V_{Z_1Z_2} - nZ_2V_{Z_1Z_2} \\
\frac{dV_{Z_2}}{dt} &= k_3 + nZ_1Z_2 - 2nZ_1V_{Z_2} - 2nZ_2V_{Z_1Z_2}
\end{aligned}
$$

and, for $E[X(t)] > 0$, $FF_X(t) = \frac{V_X}{X}$.

### S4 *AC-BioSD-II*

The *BioSD-II* topology is given by the CRN in Eq. (6) and can be described by the following RREs (see Eq. (S1)):

$$\frac{dX}{dt} = b + k_{in}UX - k_1XZ_1 \tag{S6}$$
$$\frac{dZ_1}{dt} = k_2X - nZ_1Z_2 \tag{S7}$$
$$\frac{dZ_2}{dt} = k_3 - nZ_1Z_2 \tag{S8}$$

#### S4.1 Equilibrium and stability

**Theorem 1**

For any non-negative $U^*$, the system defined by Eqs.(S6)-(S8) has a unique and positive steady state which is locally exponentially stable.

*Proof.* By setting the time derivatives in Eqs.(S6)-(S8) to zero, we obtain:

$$X^* = \frac{k_3}{k_2}$$
$$Z_1^* = \frac{k_{in}k_3U^* + bk_2}{k_1k_3}$$
$$Z_2^* = \frac{k_1k_3^2}{n(k_{in}k_3U^* + bk_2)}$$

Clearly, $X^*, Z_1^*, Z_2^* > 0$ since $k_{in}, k_1, k_2, k_3, b, n > 0$.

Jacobian linearization of Eqs.(S6)-(S8) around the fixed point $(X^*, Z_1^*, Z_2^*)$ gives:

$$\begin{bmatrix} \dot{X} \\ \dot{Z}_1 \\ \dot{Z}_2 \end{bmatrix} = \underbrace{\begin{bmatrix} -\frac{k_2b}{k_3} & -k_1X^* & 0 \\ k_2 & -nZ_2^* & -nZ_1^* \\ 0 & -nZ_2^* & -nZ_1^* \end{bmatrix}}_{M_1} \begin{bmatrix} X \\ Z_1 \\ Z_2 \end{bmatrix}$$

The characteristic polynomial of $M_1$ is:

$$P_{M_1}(s) = \det(sI - M_1) = s^3 + \zeta_2s^2 + \zeta_1s + \zeta_0$$

where $\zeta_2 = n(Z_1^* + Z_2^*) + \frac{k_2 b}{k_3}$, $\zeta_1 = \frac{nk_2 b}{k_3}(Z_1^* + Z_2^*) + k_1 k_2 X^*$ and $\zeta_0 = nk_1 k_2 X^* Z_1^*$. According to the Routh–Hurwitz criterion, $P_{M_1}(s)$ has all its roots in the open left half-plane if and only if $\zeta_2, \zeta_1, \zeta_0 > 0$ and $\zeta_2 \zeta_1 > \zeta_0$. It is evident that the positivity condition is satisfied.

The second inequality is also satisfied:

$$\left(n(Z_1^* + Z_2^*) + \frac{k_2 b}{k_3}\right)\left(\frac{nk_2 b}{k_3}(Z_1^* + Z_2^*) + k_1 k_2 X^*\right) > nk_1 k_2 X^* Z_1^*$$

or

$$n^2 \frac{k_2 b}{k_3}(Z_1^* + Z_2^*)^2 + n\frac{k_2^2 b^2}{k_3^2}(Z_1^* + Z_2^*) + nk_1 k_2 X^* Z_2^* + \frac{k_1 k_2^2 b}{k_3} X^* > 0$$

which is always true.

Consequently, $M_1$ is Hurwitz and the steady state ($X^*$, $Z_1^*$, $Z_2^*$) is locally exponentially stable. □

#### S4.2 Filtered derivative action

**Theorem 2**

In the neighbourhood of ($U^*$, $X^*$, $Z_1^*$, $Z_2^*$) and in the limit as $n \to \infty$, the input/output relation of the system defined by Eqs.(S6)-(S8) in the Laplace domain can be described as:

$$\tilde{T}_{AB}(s) = \frac{\tilde{x}(s)}{\tilde{u}(s)} = \frac{k_{in}}{k_1 k_2} \frac{s}{\varepsilon_{AB}(s^2 + s) + 1} \quad \text{with} \quad \varepsilon_{AB} = \frac{b^2 k_2^2}{k_1 k_3^3}$$

where $\tilde{u}(s)$, $\tilde{x}(s)$ are the Laplace transform of $u = U - U^*$, $x = X - X^*$, respectively and $s$ is the Laplace variable (complex frequency).

*Proof.* We adopt the coordinate transformations: $u = U - U^*$, $x = X - X^*$, $z_1 = Z_1 - Z_1^*$, $z_2 = Z_2 - Z_2^*$ representing sufficiently small perturbations around ($U^*$, $X^*$, $Z_1^*$, $Z_2^*$) and we obtain via Jacobian linearization of Eqs.(S6)-(S8):

$$\begin{bmatrix} \dot{x} \\ \dot{z}_1 \\ \dot{z}_2 \end{bmatrix} = \underbrace{\begin{bmatrix} -\frac{k_2 b}{k_3} & -k_1 X^* & 0 \\ k_2 & -nZ_2^* & -nZ_1^* \\ 0 & -nZ_2^* & -nZ_1^* \end{bmatrix}}_{M_1} \begin{bmatrix} x \\ z_1 \\ z_2 \end{bmatrix} + \begin{bmatrix} k_{in} X^* \\ 0 \\ 0 \end{bmatrix} u$$

Subsequently, by applying the transformation $g = z_1 - z_2$ we get:

$$\begin{bmatrix} \dot{x} \\ \dot{g} \\ \dot{z}_2 \end{bmatrix} = \begin{bmatrix} -\frac{k_2 b}{k_3} & -k_1 X^* & -k_1 X^* \\ k_2 & 0 & 0 \\ 0 & -nZ_2^* & -n(Z_1^* + Z_2^*) \end{bmatrix} \begin{bmatrix} x \\ g \\ z_2 \end{bmatrix} + \begin{bmatrix} k_{in} X^* \\ 0 \\ 0 \end{bmatrix} u \tag{S9}$$

The idea is to show that $z_2$ goes to zero rapidly as $n \to \infty$. This is intuitively plausible from (S9), but one needs to be careful, because $Z_2^*$ depends on $n$ (as shown in the proof of Theorem 1).

To show this, we introduce the nondimensional variables $t_n = \frac{k_3}{Z_1^*} t$, $x_n = \frac{x}{\beta_1}$ (where $\beta_1$ is an arbitrary scaling parameter with the same units as $x$), $g_n = \frac{k_3}{k_2 \beta_1 Z_1^*} g$, $z_{2n} = \frac{k_3}{k_2 \beta_1 Z_1^*} z_2$, $u_n = \frac{k_{in} X^* Z_1^*}{k_3 \beta_1} u$ and the nondimensional parameters $\gamma_1 = \frac{k_3}{n Z_1^{*2}}$, $\gamma_2 = \frac{k_2 b Z_1^*}{k_3^2}$, $\gamma_3 = \frac{k_1 k_2 X^* Z_1^{*2}}{k_3^2}$.

Eq. (S9) can thus be written as:

$$\frac{dx_n}{dt} = u_n - \gamma_2 x_n - \gamma_3 g_n - \gamma_3 z_{2n} \tag{S10}$$

$$\frac{dg_n}{dt} = x_n \tag{S11}$$

$$\gamma_1 \frac{dz_{2n}}{dt} = -\gamma_1 g_n - (1 + \gamma_1) z_{2n} \tag{S12}$$

We now treat $\gamma_1$ as a singular perturbation parameter and invoke Tikhonov's theorem (Theorem 11.1 in [7]). Specifically, as $\gamma_1 \to 0$, we get the reduced model:

$$\frac{dx_n}{dt} = u_n - \gamma_2 x_n - \gamma_3 g_n \tag{S13}$$

$$\frac{dg_n}{dt} = x_n \tag{S14}$$

since $z_{2n} \to 0$. Also, given a finite time interval $[0, t_f]$ and some initial conditions of interest, the reduced problem given by Eqs. (S15), (S16) has a unique solution, $\bar{x}_n(t)$, $\bar{g}_n(t)$. In addition, the origin is an exponentially stable equilibrium point of the boundary-layer model $\frac{dz_{2n}}{d\tau} = -z_{2n}$, where $\tau = \frac{t_n}{\gamma_1}$.

Consequently, there exists a positive constant $\gamma_1^*$ such that for $0 < \gamma_1^* < \gamma_1$ the singular perturbation problem given by Eqs. (S10)–(S12) has a unique solution $x_n(t,\gamma_1)$, $g_n(t,\gamma_1)$, $z_{2n}(t,\gamma_1)$ on $[0,t_f]$ and:

$$\begin{aligned} x_n(t,\gamma_1) - \bar{x}_n(t) &= \mathscr{O}(\gamma_1) \\ g_n(t,\gamma_1) - \bar{g}_n(t) &= \mathscr{O}(\gamma_1) \end{aligned}$$

Moreover, given any $t_b > 0$, there is $\gamma_1^{**} \leq \gamma_1*$ such that

$$z_{2,n}(t,\gamma 1) = \mathscr{O}(\gamma_1)$$

whenever $\gamma_1 < \gamma_1^{**}$. Note that $\gamma_1, Z_2^* \to 0$ as $n \to \infty$.

Finally, we consider the nondimensional variables $\breve{t}_n = \gamma_2 t_n$, $\breve{x}_n = \dfrac{x_n}{\beta_2}$ (where $\beta_2$ is an arbitrary nondimensial scaling parameter), $\breve{g}_n = \dfrac{\gamma_2}{\beta_2} g_n$, $\breve{u}_n = \dfrac{\gamma_2}{\gamma_3\beta_2} u_n$ and the nondimensional parameter $\varepsilon_{AB} = \dfrac{b^2 k_2^2}{k_1 k_3^3}$.

Eqs. (S15), (S16) can thus be written as:

$$\varepsilon_{AB} \frac{d\breve{x}_n}{dt} = -\varepsilon_{AB}\breve{x}_n - \breve{g}_n + \breve{u}_n \tag{S15}$$

$$\frac{d\breve{g}_n}{dt} = \breve{x}_n \tag{S16}$$

which results in the following transfer function in the Laplace domain:

$$\tilde{T}_{AB}(s) = \frac{\tilde{\breve{x}}_n(s)}{\tilde{\breve{u}}_n(s)} = \frac{s}{\varepsilon_{AB}(s^2+s)+1}$$

or, equivalently:

$$\tilde{T}_{AB}(s) = \frac{\tilde{x}(s)}{\tilde{u}(s)} = \frac{k_{in}}{k_1 k_2} \frac{s}{\varepsilon_{AB}(s^2+s)+1}$$

□

**Remark 1**

In practice, to ensure that $\gamma_1$ is sufficiently small, we can impose the constraint $\gamma_1 \ll 1$, or equivalently,

$$n \gg \frac{k_1^2 k_3^3}{(k_{in} k_3 U^* + b k_2)^2}$$

**Remark 2**

For a sufficiently slow input signal and after a sufficiently long time period, the output of the *BioSD-II* topology can be approximated in the time domain as:

$$X = \frac{k_{in}}{k_1 k_2}\frac{dU}{dt} + \frac{k_3}{k_2}$$

#### S4.3 Fano factor

In the special case where $U = 0$, the output Fano factor is given by Eq. (S5). If we now consider the input signal described by the CRN in Eq. (3) and assume $E[X(t)] > 0$, we have $FF_X(t) = \frac{V_X}{X}$ where (see Eqs. (S1), (S4)):

$$\begin{aligned}
\frac{dU}{dt} &= cp_1 - p_2 U \\
\frac{dX}{dt} &= b + k_{in}UX - k_1 X Z_1 \\
\frac{dZ_1}{dt} &= k_2 X - nZ_1 Z_2 \\
\frac{dZ_2}{dt} &= k_3 - nZ_1 Z_2 \\
\frac{dV_U}{dt} &= p_1 c^2 + p_2 U - 2p_2 V_U \\
\frac{dV_{UX}}{dt} &= V_{UX}(k_{in}U - k_1 Z_1) - p_2 V_{UX} - k_1 X V_{UZ_1} + k_{in} X V_U \\
\frac{dV_{UZ_1}}{dt} &= k_2 V_{UX} - p_2 V_{UZ_1} - nZ_1 V_{UZ_2} - nZ_2 V_{UZ_1} \\
\frac{dV_{UZ_2}}{dt} &= -p_2 V_{UZ_2} - nZ_1 V_{UZ_2} - nZ_2 V_{UZ_1} \\
\frac{dV_X}{dt} &= b + 2V_X(k_{in}U - k_1 Z_1) + k_{in}UX + k_1 X Z_1 - 2k_1 X V_{XZ_1} + 2k_{in} X V_{UX} \\
\frac{dV_{XZ_1}}{dt} &= k_2 V_X + V_{XZ_1}(k_{in}U - k_1 Z_1) - k_1 X V_{Z_1} + k_{in} X V_{UZ_1} - nZ_1 V_{XZ_2} - nZ_2 V_{XZ_1} \\
\frac{dV_{XZ_2}}{dt} &= V_{XZ_2}(k_{in}U - k_1 Z_1) - k_1 X V_{Z_1 Z_2} + k_{in} X V_{UZ_2} - nZ_1 V_{XZ_2} - nZ_2 V_{XZ_1} \\
\frac{dV_{Z_1}}{dt} &= k_2 X + 2k_2 V_{XZ_1} + nZ_1 Z_2 - 2nZ_2 V_{Z_1} - 2nZ_1 V_{Z_1 Z_2} \\
\frac{dV_{Z_1 Z_2}}{dt} &= k_2 V_{XZ_2} + nZ_1 Z_2 - nZ_1 V_{Z_2} - nZ_2 V_{Z_1} - nZ_1 V_{Z_1 Z_2} - nZ_2 V_{Z_1 Z_2} \\
\frac{dV_{Z_2}}{dt} &= k_3 + nZ_1 Z_2 - 2nZ_1 V_{Z_2} - 2nZ_2 V_{Z_1 Z_2}
\end{aligned}$$

### S5 Approximate analytical description of the Fano factor

To study the output noise levels in the CRNs under investigation, we use the Fano factor, which is defined as the variance normalized by the mean number of molecules. Both the mean and the variance can be determined by the solution of an ODE system obtained through LNA. Because this ODE system is generally nonlinear, deriving a full closed-form solution for the Fano factor is highly challenging. In cases where the system evolves sufficiently close to its steady state, this obstacle can be circumvented by applying Jacobian linearization to approximate the local dynamics. Assuming $m$ differential equations, we can therefore obtain an ODE system of the form:

$$\frac{dy}{dt} = Ay \tag{S17}$$

where $y \in \mathbb{R}^m$ and $A \in \mathbb{R}^{m\times m}$. For some initial condition $y_0$ at time $t = 0$, we have the following solution:

$$y = e^{At} y_0 \quad \text{with} \quad e^{At} = I + \sum_{m=1}^{\infty} \frac{A^m t^m}{m!} \tag{S18}$$

where $I$ is an $m \times m$ identity matrix. In principle, the matrix exponential in Eq. (S18) can be computed in various ways; a survey of the most important techniques of practical interest can be found in [8].

As a demonstration example, we consider the (LNA-derived) nonlinear system of ODEs describing the behaviour of *BioSD-II* in the presence of a bursty birth-death process as an input signal (see Section S3). Assuming an equilibrium of interest, ($X^*$, $Z_1^*$, $Z_2^*$, $V_{UZ_1}^*$, $V_{UX}^*$, $V_{UZ_2}^*$, $V_{XZ_1}^*$, $V_x^*$, $V_{Z_1}^*$, $V_{XZ_2}^*$, $V_{Z_1Z_2}^*$, $V_{Z_2}^*$), we apply Jacobian linearization to obtain:

$$
\begin{aligned}
\frac{du}{dt} &= -p_2 u \\
\frac{dx}{dt} &= -k_1 Z_1^* x - k_1 X^* z_1 + k_{in} u \\
\frac{dz_1}{dt} &= k_2 x - n Z_2^* z_1 - n Z_1^* z_2 \\
\frac{dz_2}{dt} &= -n Z_2^* z_1 - n Z_1^* z_2 \\
\frac{dv_u}{dt} &= p_2 u - 2 p_2 v_u \\
\frac{dv_{ux}}{dt} &= k_{in} v_u - p_2 v_{ux} - k_1 X^* v_{uz_1} - k_1 V_{UZ_1}^* x - k_1 Z_1^* v_{ux} - k_1 V_{UX}^* z_1 \\
\frac{dv_{uz_1}}{dt} &= k_2 v_{ux} - p_2 v_{uz_1} - n Z_1^* v_{uz_2} - n V_{UZ_2}^* z_1 - n Z_2^* v_{uz_1} - n V_{UZ_1}^* z_2 \\
\frac{dv_{uz_2}}{dt} &= -p_2 v_{uz_2} - n Z_1^* v_{uz_2} - n V_{UZ_2}^* z_1 - n Z_2^* v_{uz_1} - n V_{UZ_1}^* z_2 \\
\frac{dv_x}{dt} &= k_{in} u + 2 k_{in} v_{ux} + k_1 Z_1^* x + k_1 X^* z_1 - 2 k_1 X^* v_{xz_1} - 2 k_1 V_{XZ_1}^* x - 2 k_1 Z_1^* v_x - 2 k_1 V_x^* z_1 \\
\frac{dv_{xz_1}}{dt} &= k_2 v_x + k_{in} v_{uz_1} - k_1 Z_1^* v_{xz_1} - k_1 V_{XZ_1}^* z_1 - k_1 X^* v_{z_1} - k_1 V_{Z1}^* x \\
&\quad - n Z_1^* v_{xz_2} - n V_{XZ_2}^* z_1 - n Z_2^* v_{xz_1} - n V_{XZ_1}^* z_2 \\
\frac{dv_{xz_2}}{dt} &= k_{in} v_{uz_2} - k_1 X^* v_{z_1 z_2} - k_1 V_{Z_1 Z_2}^* x - (k_1 + n) Z_1^* v_{xz_2} - (k_1 + n) V_{XZ_2}^* z_1 \\
&\quad - n Z_2^* v_{xz_1} - n V_{XZ_1}^* z_2 \\
\frac{dv_{z_1}}{dt} &= k_2 x + 2 k_2 v_{xz_1} + n Z_2^* z_1 + n Z_1^* z_2 - 2 n Z_2^* v_{z_1} - 2 n V_{Z_1}^* z_2 - 2 n Z_1^* v_{z_1 z_2} - 2 n V_{Z_1 Z_2}^* z_1 \\
\frac{dv_{z_1 z_2}}{dt} &= k_2 v_{xz_2} + n Z_2^* z_1 + n Z_1^* z_2 - n Z_1^* v_{z_2} - n V_{Z_2}^* z_1 - n Z_2^* v_{z_1} - n V_{Z_1}^* z_2 \\
&\quad - n Z_1^* v_{z_1 z_2} - n V_{Z_1 Z_2}^* z_1 - n Z_2^* v_{z_1 z_2} - n V_{Z_1 Z_2}^* z_2 \\
\frac{dv_{z_2}}{dt} &= n Z_2^* z_1 + n Z_1^* z_2 - 2 n Z_1^* v_{z_2} - 2 n V_{Z_2}^* z_1 - 2 n Z_2^* v_{z_1 z_2} - 2 n V_{Z_1 Z_2}^* z_2
\end{aligned}
$$

which can be expressed in the form of Eq. (S17).

In Fig. S2, we provide a numerical comparison between the nonlinear model and its linearization with respect to the output mean, variance, and Fano factor, assuming also a perturbation of 15% in the initial conditions. The results show that the linearized model offers a satisfactory approximation of the local dynamics of the nonlinear system.

### S6 External degradation

#### S6.1 *AC-BioSD-II*: Equilibrium and stability

In the presence of output degradation, the *AC-BioSD-II* topology is described by the CRN in Eq. (6), supplemented with the reaction $X \xrightarrow{d} \varnothing$. The corresponding RREs are:

$$\frac{dX}{dt} = b + k_{in}UX - k_1XZ_1 - dX \tag{S19}$$

$$\frac{dZ_1}{dt} = k_2X - nZ_1Z_2 \tag{S20}$$

$$\frac{dZ_2}{dt} = k_3 - nZ_1Z_2 \tag{S21}$$

**Theorem 3**

For any $d < \frac{bk_2}{k_3} + k_{in}U^*$ with $U^* \geq 0$, the system defined by Eqs.(S19)-(S21) has a unique and positive steady state which is locally exponentially stable.

*Proof.* By setting the time derivatives in Eqs.(S19)-(S21) to zero, we obtain:

$$X^* = \frac{k_3}{k_2}$$

$$Z_1^* = \frac{k_3k_{in}U^* + bk_2 - dk_3}{k_1k_3}$$

$$Z_2^* = \frac{k_1k_3^2}{n(k_3k_{in}U^* + bk_2 - dk_3)}.$$

Since $k_{in}$, $k_1$, $k_2$, $k_3$, $b$, $n > 0$, for any $d < \frac{bk_2}{k_3} + k_{in}U^*$ with $U^* \geq 0$, we have $X^*$, $Z_1^*$, $Z_2^* > 0$.

Jacobian linearization of Eqs.(S19)-(S21) around the fixed point ($X^*$, $Z_1^*$, $Z_2^*$) gives:

$$\begin{bmatrix} \dot{X} \\ \dot{Z}_1 \\ \dot{Z}_2 \end{bmatrix} = \underbrace{\begin{bmatrix} -\frac{k_2b}{k_3} & -k_1X^* & 0 \\ k_2 & -nZ_2^* & -nZ_1^* \\ 0 & -nZ_2^* & -nZ_1^* \end{bmatrix}}_{M_2} \begin{bmatrix} X \\ Z_1 \\ Z_2 \end{bmatrix}$$

Observe that $M_2$ coincides with $M_1$, which has been shown to be Hurwitz (see **Theorem 1**). Consequently, the steady state $(X^*, Z_1^*, Z_2^*)$ is locally exponentially stable.

Note that, assuming $d < \frac{bk_2}{k_3} + k_{in}U^*$ with $U^* \geq 0$, **Theorem 2** also applies to the system given by Eqs. (S19)–(S21). □

#### S6.2 Fano factor

For $E[X(t)] > 0$, the output Fano factor is given by $FF_X(t) = \frac{V_X}{X}$ where $V_X$, $X$ can be determined by an ODE system based on the CRN under consideration (see Eqs. (S1), (S4)). Specifically, we have the following cases:

##### S6.2.1 Degradation of the output species

No input signal: CRN in Eq. (1) and $X \xrightarrow{d} \varnothing$ assuming $U = 0$

$$
\begin{aligned}
\frac{dX}{dt} &= b - k_1 X Z_1 - dX \\
\frac{dZ_1}{dt} &= k_2 X - n Z_1 Z_2 \\
\frac{dZ_2}{dt} &= k_3 - n Z_1 Z_2 \\
\frac{dV_X}{dt} &= dX + b + k_1 X Z_1 - 2k_1 X V_{XZ_1} - 2(k_1 Z_1 + d) V_X \\
\frac{dV_{XZ_1}}{dt} &= k_2 V_X - k_1 X V_{Z_1} - (k_1 Z_1 + d) V_{XZ_1} - n Z_1 V_{XZ_2} - n Z_2 V_{XZ_1} \\
\frac{dV_{XZ_2}}{dt} &= -k_1 X V_{Z_1 Z_2} - (k_1 Z_1 + d) V_{XZ_2} - n Z_1 V_{XZ_2} - n Z_2 V_{XZ_1} \\
\frac{dV_{Z_1}}{dt} &= k_2 X + 2k_2 V_{XZ_1} + n Z_1 Z_2 - 2n Z_2 V_{Z_1} - 2n Z_1 V_{Z_1 Z_2} \\
\frac{dV_{Z_1 Z_2}}{dt} &= k_2 V_{XZ_2} + n Z_1 Z_2 - n Z_1 V_{Z_2} - n Z_2 V_{Z_1} - n Z_1 V_{Z_1 Z_2} - n Z_2 V_{Z_1 Z_2} \\
\frac{dV_{Z_2}}{dt} &= k_3 + n Z_1 Z_2 - 2n Z_1 V_{Z_2} - 2n Z_2 V_{Z_1 Z_2}
\end{aligned}
$$

*BioSD-II* with input signal: CRNs in Eqs. (1), (3) and $X \xrightarrow{d} \varnothing$

$$
\begin{aligned}
\frac{dU}{dt} &= cp_1 - p_2U \\
\frac{dX}{dt} &= b + k_{in}U - k_1XZ_1 - dX \\
\frac{dZ_1}{dt} &= k_2X - nZ_1Z_2 \\
\frac{dZ_2}{dt} &= k_3 - nZ_1Z_2 \\
\frac{dV_U}{dt} &= p_1c^2 + p_2U - 2p_2V_U \\
\frac{dV_{UX}}{dt} &= k_{in}V_U - p_2V_{UX} - k_1XV_{UZ_1} - (k_1Z_1 + d)V_{UX} \\
\frac{dV_{UZ_1}}{dt} &= k_2V_{UX} - p_2V_{UZ_1} - nZ_1V_{UZ_2} - nZ_2V_{UZ_1} \\
\frac{dV_{UZ_2}}{dt} &= -p_2V_{UZ_2} - nZ_1V_{UZ_2} - nZ_2V_{UZ_1} \\
\frac{dV_X}{dt} &= b + dX + k_{in}U + 2k_{in}V_{UX} + k_1XZ_1 - 2k_1XV_{XZ_1} - 2(d + k_1Z_1)V_X \\
\frac{dV_{XZ_1}}{dt} &= k_2V_X + k_{in}V_{UZ_1} - k_1XV_{Z_1} - (d + k_1Z_1)V_{XZ_1} - nZ_1V_{XZ_2} - nZ_2V_{XZ_1} \\
\frac{dV_{XZ_2}}{dt} &= k_{in}V_{UZ_2} - k_1XV_{Z_1Z_2} - (d + k_1Z_1)V_{XZ_2} - nZ_1V_{XZ_2} - nZ_2V_{XZ_1} \\
\frac{dV_{Z_1}}{dt} &= k_2X + 2k_2V_{XZ_1} + nZ_1Z_2 - 2nZ_2V_{Z_1} - 2nZ_1V_{Z_1Z_2} \\
\frac{dV_{Z_1Z_2}}{dt} &= k_2V_{XZ_2} + nZ_1Z_2 - nZ_1V_{Z_2} - nZ_2V_{Z_1} - nZ_1V_{Z_1Z_2} - nZ_2V_{Z_1Z_2} \\
\frac{dV_{Z_2}}{dt} &= k_3 + nZ_1Z_2 - 2nZ_1V_{Z_2} - 2nZ_2V_{Z_1Z_2}
\end{aligned}
$$

*AC-BioSD-II* with input signal: CRNs in Eqs. (3), (6) and $X \xrightarrow{d} \varnothing$

$$
\begin{aligned}
\frac{dU}{dt} &= cp_1 - p_2U \\
\frac{dX}{dt} &= b + k_{in}UX - k_1XZ_1 - dX \\
\frac{dZ_1}{dt} &= k_2X - nZ_1Z_2 \\
\frac{dZ_2}{dt} &= k_3 - nZ_1Z_2 \\
\frac{dV_U}{dt} &= p_1c^2 + p_2U - 2p_2V_U \\
\frac{dV_{UX}}{dt} &= -V_{UX}(d - k_{in}U + k_1Z_1) - p_2V_{UX} - k_1XV_{UZ_1} + k_{in}XV_U \\
\frac{dV_{UZ_1}}{dt} &= k_2V_{UX} - p_2V_{UZ_1} - nZ_1V_{UZ_2} - nZ_2V_{UZ_1} \\
\frac{dV_{UZ_2}}{dt} &= -p_2V_{UZ_2} - nZ_1V_{UZ_2} - nZ_2V_{UZ_1} \\
\frac{dV_X}{dt} &= dX + b - 2V_X(d - k_{in}U + k_1Z_1) + k_{in}UX + k_1XZ_1 - 2k_1XV_{XZ_1} + 2k_{in}XV_{UX} \\
\frac{dV_{XZ_1}}{dt} &= k_2V_X - V_{XZ_1}(d - k_{in}U + k_1Z_1) - k_1XV_{Z_1} + k_{in}XV_{UZ_1} - nZ_1V_{XZ_2} - nZ_2V_{XZ_1} \\
\frac{dV_{XZ_2}}{dt} &= -V_{XZ_2}(d - k_{in}U + k_1Z_1) - k_1XV_{Z_1Z_2} + k_{in}XV_{UZ_2} - nZ_1V_{XZ_2} - nZ_2V_{XZ_1} \\
\frac{dV_{Z_1}}{dt} &= k_2X + 2k_2V_{XZ_1} + nZ_1Z_2 - 2nZ_2V_{Z_1} - 2nZ_1V_{Z_1Z_2} \\
\frac{dV_{Z_1Z_2}}{dt} &= k_2V_{XZ_2} + nZ_1Z_2 - nZ_1V_{Z_2} - nZ_2V_{Z_1} - nZ_1V_{Z_1Z_2} - nZ_2V_{Z_1Z_2} \\
\frac{dV_{Z_2}}{dt} &= k_3 + nZ_1Z_2 - 2nZ_1V_{Z_2} - 2nZ_2V_{Z_1Z_2}
\end{aligned}
$$

228 ### S6.2.2 Degradation of all species

229 No input signal: CRN in Eq. (1) and $X, Z_1, Z_2 \xrightarrow{d} \varnothing$ assuming $U = 0$

$$\frac{dX}{dt} = b - k_1 Z_1 X - d X$$
$$\frac{dZ_1}{dt} = k_2 X - n Z_1 Z_2 - d Z_1$$
$$\frac{dZ_2}{dt} = k_3 - n Z_1 Z_2 - d Z_2$$
$$\frac{dV_X}{dt} = b + d X - 2 (d + k_1 Z_1) V_X + k_1 Z_1 X - 2 k_1 X V_{XZ_1}$$
$$\frac{dV_{XZ_1}}{dt} = k_2 V_X - (d + k_1 Z_1) V_{XZ_1} - (d + n Z_2) V_{XZ_1} - k_1 X V_{Z_1} - n Z_1 V_{XZ_2}$$
$$\frac{dV_{XZ_2}}{dt} = -(d + k_1 Z_1) V_{XZ_2} - (d + n Z_1) V_{XZ_2} - k_1 X V_{Z_1 Z_2} - n Z_2 V_{XZ_1}$$
$$\frac{dV_{Z_1}}{dt} = d Z_1 + k_2 X + 2 k_2 V_{XZ_1} - 2 (d + n Z_2) V_{Z_1} + n Z_1 Z_2 - 2 n Z_1 V_{Z_1 Z_2}$$
$$\frac{dV_{Z_1 Z_2}}{dt} = k_2 V_{XZ_2} - (d + n Z_1) V_{Z_1 Z_2} - (d + n Z_2) V_{Z_1 Z_2} + n Z_1 Z_2 - n Z_1 V_{Z_2} - n Z_2 V_{Z_1}$$
$$\frac{dV_{Z_2}}{dt} = k_3 + d Z_2 - 2 (d + n Z_1) V_{Z_2} + n Z_1 Z_2 - 2 n Z_2 V_{Z_1 Z_2}$$

*BioSD-II* with input signal: CRNs in Eqs. (1), (3) and $X, Z_1, Z_2 \xrightarrow{d} \varnothing$

$$
\begin{aligned}
\frac{dU}{dt} &= c\,p_1 - p_2\,U \\
\frac{dX}{dt} &= b + k_{in}\,U - k_1\,Z_1\,X - d\,X \\
\frac{dZ_1}{dt} &= k_2\,X - n\,Z_2\,Z_1 - d\,Z_1 \\
\frac{dZ_2}{dt} &= k_3 - n\,Z_1\,Z_2 - d\,Z_2 \\
\frac{dV_U}{dt} &= p_1\,c^2 + p_2\,U - 2\,p_2\,V_U \\
\frac{dV_{UX}}{dt} &= k_{in}\,V_U - p_2\,V_{UX} - (d + k_1\,Z_1)\,V_{UX} - k_1\,X\,V_{UZ_1} \\
\frac{dV_{UZ_1}}{dt} &= k_2\,V_{UX} - p_2\,V_{UZ_1} - (d + n\,Z_2)\,V_{UZ_1} - n\,Z_1\,V_{UZ_2} \\
\frac{dV_{UZ_2}}{dt} &= -p_2\,V_{UZ_2} - (d + n\,Z_1)\,V_{UZ_2} - n\,Z_2\,V_{UZ_1} \\
\frac{dV_X}{dt} &= b + d\,X + k_{in}\,U + 2\,k_{in}\,V_{UX} - 2\,(d + k_1\,Z_1)\,V_X + k_1\,Z_1\,X - 2\,k_1\,X\,V_{XZ_1} \\
\frac{dV_{XZ_1}}{dt} &= k_2\,V_X + k_{in}\,V_{UZ_1} - (d + k_1\,Z_1)\,V_{XZ_1} - (d + n\,Z_2)\,V_{XZ_1} - k_1\,X\,V_{Z_1} - n\,Z_1\,V_{XZ_2} \\
\frac{dV_{XZ_2}}{dt} &= k_{in}\,V_{UZ_2} - (d + k_1\,Z_1)\,V_{XZ_2} - (d + n\,Z_1)\,V_{XZ_2} - k_1\,X\,V_{Z_1Z_2} - n\,Z_2\,V_{XZ_1} \\
\frac{dV_{Z_1}}{dt} &= d\,Z_1 + k_2\,X + 2\,k_2\,V_{XZ_1} - 2\,(d + n\,Z_2)\,V_{Z_1} + n\,Z_1\,Z_2 - 2\,n\,Z_1\,V_{Z_1Z_2} \\
\frac{dV_{Z_1Z_2}}{dt} &= k_2\,V_{XZ_2} - (d + n\,Z_1)\,V_{Z_1Z_2} - (d + n\,Z_2)\,V_{Z_1Z_2} + n\,Z_1\,Z_2 - n\,Z_1\,V_{Z_2} - n\,Z_2\,V_{Z_1} \\
\frac{dV_{Z_2}}{dt} &= k_3 + d\,Z_2 - 2\,(d + n\,Z_1)\,V_{Z_2} + n\,Z_1\,Z_2 - 2\,n\,Z_2\,V_{Z_1Z_2}
\end{aligned}
$$

<u>*AC-BioSD-II* with input signal: CRNs in Eqs. (3), (6) and $X, Z_1, Z_2 \xrightarrow{d} \varnothing$</u>

$$
\begin{aligned}
\frac{dU}{dt} &= p_1\, c - p_2\, U \\
\frac{dX}{dt} &= b + k_{in}\, U\, X - k_1\, Z_1\, X - d\, X \\
\frac{dZ_1}{dt} &= k_2 X - n\, Z_1\, Z_2 - d\, Z_1 \\
\frac{dZ_2}{dt} &= k_3 - n\, Z_1\, Z_2 - d\, Z_2 \\
\frac{dV_U}{dt} &= p_1\, c^2 + p_2\, U - 2\, p_2\, V_U \\
\frac{dV_{UX}}{dt} &= k_{in} X\, V_U - (d - k_{in}\, U + k_1\, Z_1)\, V_{UX} - k_1\, X\, V_{UZ_1} - p_2\, V_{UX} \\
\frac{dV_{UZ_1}}{dt} &= k_2\, V_{UX} - p_2\, V_{UZ_1} - (d + n\, Z_2)\, V_{UZ_1} - n\, Z_1\, V_{UZ_2} \\
\frac{dV_{UZ_2}}{dt} &= -p_2\, V_{UZ_2} - (d + n\, Z_1)\, V_{UZ_2} - n\, Z_2\, V_{UZ_1} \\
\frac{dV_X}{dt} &= b + d\, X - 2\, (d - k_{in}\, U + k_1\, Z_1)\, V_X + k_{in}\, U\, X + k_1\, Z_1\, X - 2\, k_1\, X\, V_{XZ_1} + 2\, k_{in} X\, V_{UX} \\
\frac{dV_{XZ_1}}{dt} &= k_2\, V_X - (d + n\, Z_2)\, V_{XZ_1} - (d - k_{in}\, U + k_1\, Z_1)\, V_{XZ_1} - k_1\, X\, V_{Z_1} + k_{in} X\, V_{UZ_1} - n\, Z_1\, V_{XZ_2} \\
\frac{dV_{XZ_2}}{dt} &= k_{in} X\, V_{UZ_2} - (d - k_{in}\, U + k_1\, Z_1)\, V_{XZ_2} - k_1\, X\, V_{Z_1 Z_2} - (d + n\, Z_1)\, V_{XZ_2} - n\, Z_2\, V_{XZ_1} \\
\frac{dV_{Z_1}}{dt} &= d\, Z_1 + k_2 X + 2\, k_2\, V_{XZ_1} - 2\, (d + n\, Z_2)\, V_{Z_1} + n\, Z_1\, Z_2 - 2\, n\, Z_1\, V_{Z_1 Z_2} \\
\frac{dV_{Z_1 Z_2}}{dt} &= k_2\, V_{XZ_2} - (d + n\, Z_1)\, V_{Z_1 Z_2} - (d + n\, Z_2)\, V_{Z_1 Z_2} + n\, Z_1\, Z_2 - n\, Z_1\, V_{Z_2} - n\, Z_2\, V_{Z_1} \\
\frac{dV_{Z_2}}{dt} &= k_3 + d\, Z_2 - 2\, (d + n\, Z_1)\, V_{Z_2} + n\, Z_1\, Z_2 - 2\, n\, Z_2\, V_{Z_1 Z_2}
\end{aligned}
$$

### S7 Negative feedback control based on *BioSD-II*

#### S7.1 Derivative control

We consider the CRN in Eq. (8) and a process to be controlled with $v$ species, $Y_1$, $Y_2$,..., $Y_{v-1}$, $Y_t$. By using RREs, the dynamics of the resulting closed-loop topology can be described as:

$$\frac{dY}{dt} = f(Y) + \xi \underbrace{(-k_f X Y_t)}_{\substack{\text{control}\\ \text{input signal}}} \tag{S22}$$

$$\frac{dX}{dt} = b + k_{in} Y_t - k_1 X Z_1 \tag{S23}$$

$$\frac{dZ_1}{dt} = k_2 X - n Z_1 Z_2 \tag{S24}$$

$$\frac{dZ_2}{dt} = k_3 - n Z_1 Z_2 \tag{S25}$$

where $f(Y)$ represents the open-loop process dynamics, $Y = [Y_1 \;\; Y_2 \;\; \dots \;\; Y_{v-1} \;\; Y_t]^T$ and $\xi = [0 \;\; 0 \;\; \dots \;\; 1]^T \in \mathbb{Z}^v$.

Assuming an equilibrium of interest, denoted as $E$, by applying Jacobian linearization on Eqs. (S22)-(S25) we get:

$$\frac{dy}{dt} = Ay + \xi u_{CL} \tag{S26}$$

$$\frac{dx}{dt} = k_{in} y_t - k_1 Z_1^* x - k_1 X^* z_1 \tag{S27}$$

$$\frac{dz_1}{dt} = k_2 x - n Z_2^* z_1 - n Z_1^* z_2 \tag{S28}$$

$$\frac{dz_2}{dt} = -n Z_2^* z_1 - n Z_1^* z_2 \tag{S29}$$

where $y = [y_1 \;\; y_2 \;\; \dots \;\; y_{v-1} \;\; y_t]^T$ and $A = \left.\frac{\partial f(Y)}{\partial Y}\right|_E$. The coordinate transformations $y = [y_1 \;\; y_2 \;\; \dots \;\; y_{v-1} \;\; y_t]^T = [Y_1 \;\; Y_2 \;\; \dots \;\; Y_{v-1} \;\; Y_t]^T - [Y_1^* \;\; Y_2^* \;\; \dots \;\; Y_{v-1}^* \;\; Y_t^*]^T$, $x = X - X^*$, $z_1 = Z_1 - Z_1^*$, $z_2 = Z_2 - Z_2^*$ represent sufficiently small perturbations around the fixed point $E$, ($Y_1^*$, $Y_2^*$,..., $Y_{v-1}^*$, $Y_t^*$, $X^*$, $Z_1^*$, $Z_2^*$). We also have $u_{CL} = -k_f X^* y_t - k_f Y_t^* x$ or, equivalently, in the Laplace domain:

$$\tilde{u}_{CL}(s) = \underbrace{-k_p \tilde{y}_t(s)}_{\substack{\text{Proportional}\\ \text{control}}} \underbrace{-k_d \frac{s\tilde{y}_t(s)}{\varepsilon_B(s^2+s)+1}}_{\substack{\text{Filtered derivative}\\ \text{control}}} \tag{S30}$$

with $k_p = k_f X^*$, $k_d = \dfrac{k_{in} k_f Y_t^*}{k_1 k_3}$.

Considering the same open-loop process as before, we now examine a more general case in which the feedback control action is applied to a different, arbitrarily selected species, $Y_1$ (see Fig. S11a). We further assume that an increase in $Y_1$ results in a corresponding increase in $Y_t$ (positive process gain).

The CRN describing the *BioSD-II*-based controller in this case is:

$$
\begin{aligned}
&\varnothing \xrightarrow{b} X\,, \quad X \xrightarrow{k_2} X + Z_1\,, \quad Z_1 + X \xrightarrow{k_1} Z_1\,, \quad Z_1 + Z_2 \xrightarrow{n} \varnothing\,, \\
&\varnothing \xrightarrow{k_3} Z_2\,, \quad Y_t \xrightarrow{k_{in}} Y_t + X\,, \quad Y_1 + X \xrightarrow{k_f} X
\end{aligned} \tag{S31}
$$

and the RREs capturing the closed-loop dynamics are:

$$\frac{dY}{dt} = f(Y) + \xi \underbrace{(-k_f X Y_1)}_{\substack{\text{control} \\ \text{input signal}}} \tag{S32}$$

$$\frac{dX}{dt} = b + k_{in} Y_t - k_1 X Z_1 \tag{S33}$$

$$\frac{dZ_1}{dt} = k_2 X - n Z_1 Z_2 \tag{S34}$$

$$\frac{dZ_2}{dt} = k_3 - n Z_1 Z_2 \tag{S35}$$

where $\xi = [1 \;\; 0 \;\; \dots \;\; 0]^T \in \mathbb{Z}^\nu$.

Using the same assumptions and notation as before, we apply Jacobian linearization to Eqs. (S32)-(S35) and obtain:

$$\frac{dy}{dt} = Ay + \xi u_{CL} \tag{S36}$$

$$\frac{dx}{dt} = k_{in} y_t - k_1 Z_1^* x - k_1 X^* z_1 \tag{S37}$$

$$\frac{dz_1}{dt} = k_2 x - n Z_2^* z_1 - n Z_1^* z_2 \tag{S38}$$

$$\frac{dz_2}{dt} = -n Z_2^* z_1 - n Z_1^* z_2 \tag{S39}$$

where $A = \left.\frac{\partial f(Y)}{\partial Y}\right|_E - k_f X^* \xi \xi^T$. We also have $u_{CL} = -k_f Y_1^* x$ or, equivalently, in the Laplace domain:

$$\tilde{u}_{CL}(s) = \underbrace{-k_d \frac{s\tilde{y}_t(s)}{\varepsilon_B(s^2+s)+1}}_{\text{Filtered derivative control}} \tag{S40}$$

with $k_d = \frac{k_{in} k_f Y_1^*}{k_1 k_3}$.

#### S7.2 Sufficient conditions for closed-loop stability

The systems defined by Eqs. (S26)–(S29) and (S74)–(S77) can be interpreted as the negative-feedback interconnection of two subsystems:

$$\frac{dy}{dt} = A_p y + B_p u_p \tag{S41}$$

$$w_p = C_p y + D_p u_p \tag{S42}$$

and

$$\frac{dm}{dt} = A_{bs} m + B_{bs} u_{bs} \tag{S43}$$

$$w_{bs} = C_{bs} m + D_{bs} u_{bs} \tag{S44}$$

where $m = [x \;\; z_1 \;\; z_2]^T$, $A_p = \left.\frac{\partial f(Y)}{\partial Y}\right|_E - k_f X^* \xi \xi^T$, $A_{bs} = \begin{bmatrix} -k_1 Z_1^* & -k_1 X^* & 0 \\ k_2 & -nZ_2^* & -nZ_1^* \\ 0 & -nZ_2^* & -nZ_1^* \end{bmatrix}$, $B_p = \xi \xi^T y^*$, $B_{bs} = [k_{in} \;\; 0 \;\; 0]^T$, $C_p = [0 \;\; 0 \;\; \ldots \;\; 1] \in \mathbb{Z}^\nu$, $C_{bs} = [kf \;\; 0 \;\; 0]$, $D_p = 0$, $D_{bs} = 0$ and $u_p = -w_{bs}$, $u_{bs} = w_p$.

To facilitate the forthcoming analysis, we now state a couple of useful definitions:

- Definition 2.33 in [9]: A transfer function $H(s)$ is said to be positive real (PR) if:
  - $H(s)$ is analytic in $\mathrm{Re}[s] > 0$.
  - $H(s)$ is real for positive real $s$.

– $\mathrm{Re}[H(s)] \geq 0$, for all $\mathrm{Re}[s] > 0$.

- Definition 2.77 in [9]: A rational function $H(s) \in \mathbb{C}$ is weakly strictly positive real (WSPR) if:
  – $H(s)$ is analytic in $\mathrm{Re}[s] \geq 0$.
  – $\mathrm{Re}[H(j\omega)] > 0$, for all $\omega \in (-\infty, \infty)$.

We now present a theorem that establishes sufficient conditions for the closed-loop stability of the system under consideration near an equilibrium of practical interest.

**Theorem 4**

If the subsystem given by Eqs. (S41)–(S42) is stabilizable and detectable and $H_p(s) = C_p(sI - A_p)^{-1}B_p + D_p$ is WSPR, then the closed-loop system given by Eqs. (S41)–(S44) is asymptotically stable.

*Proof.* The system given by Eqs. (S43)–(S44) is asymptotically stable as $A_{bs}$ is Hurwitz (see [10] for a detailed proof). We also calculate the associated transfer function as:

$$H_{bs} = C_{bs}(sI - A_{bs})^{-1}B_{bs} + D_{bs} = \frac{k_f k_{in} s(s+\mu_1)}{s^3 + \mu_2 s^2 + \mu_3 s + \mu_4}$$

where $\mu_1 = n(Z_1^* + Z_2^*)$, $\mu_2 = k_1 Z_1^* + \mu_1$, $\mu_3 = \mu_1 k_1 Z_1^* + k_1 k_2 X^*$, $\mu_4 = nk_1 k_2 X^* Z_1^*$.

We also have:

$$\mathrm{Re}[H_{bs}(j\omega)] = \frac{(\mu_2 - \mu_1)k_f k_{in}\omega^4 + (\mu_1\mu_3 - \mu_4)k_f k_{in}\omega^2}{(-\mu_2\omega^2 + \mu_4)^2 + \omega^2(\mu_3 - \omega^2)^2}$$

Observe that $\mu_2 - \mu_1 = k_1 Z_1^* > 0$ and $\mu_1\mu_3 - \mu_4 = \mu_1^2 k_1 Z_1^* + nk_1 k_2 X^* Z_2^* > 0$. As a result, $\mathrm{Re}[H_{bs}(j\omega)] \geq 0$ for all $[\omega \in [-\infty, \infty]$. Therefore, based on Theorem 2.45 in [9], we conclude that $H_{bs}$ is PR.

Now, consider that $H_p(s) = C_p(sI - A_p)^{-1}B_p + D_p$ is WSPR. It then follows from Corollary 1.1 in [11] that the negative feedback interconnection of a minimal realization of $H_p(s)$ with a minimal realization of $H_{bs}(s)$ is asymptotically stable.

Subsequently, using the Kalman decomposition [12], the subsystem given by Eqs. (S41)–(S42) can be rewritten as:

$$\frac{d}{dt}\begin{bmatrix} y_I \\ y_{II} \\ y_{III} \\ y_{IV} \end{bmatrix} = \begin{bmatrix} A_1 & 0 & A_6 & 0 \\ A_2 & A_3 & A_4 & A_5 \\ 0 & 0 & A_7 & 0 \\ 0 & 0 & A_8 & A_9 \end{bmatrix} \begin{bmatrix} y_I \\ y_{II} \\ y_{III} \\ y_{IV} \end{bmatrix} + \begin{bmatrix} B_1 \\ B_2 \\ 0 \\ 0 \end{bmatrix} u_p \tag{S45}$$

$$w_p = \begin{bmatrix} C_1 & 0 & C_2 & 0 \end{bmatrix} \begin{bmatrix} y_I \\ y_{II} \\ y_{III} \\ y_{IV} \end{bmatrix} \tag{S46}$$

where the states $y_I$ are both controllable and observable, the states $y_{II}$ are controllable but not observable, the states $y_{III}$ are observable but not controllable and the states $y_{IV}$ are neither controllable nor observable. Note that a minimal realization is then:

$$\frac{dy_I}{dt} = A_1 y_I + B_1 u_p \tag{S47}$$

$$w_p = C_1 y_I \tag{S48}$$

The poles of the realization given by Eqs. (S45)–(S46) are the eigenvalues of $A_1$ (both controllable and observable), together with those of $A_3$ (controllable but not observable), $A_7$ (observable but not controllable), and $A_9$ (neither controllable nor observable).

Let us now write the state vector for the closed-loop system as $y_{cl} = (y_I, m, y_{II}, y_{III}, y_{IV})$. Then the closed-loop system has the form $\dot{y}_{cl} = Ay_{cl}$ with:

$$A = \begin{bmatrix} A_1 & -B_1 C_{bs} & 0 & A_6 & 0 \\ B_{bs} C_1 & A_{bs} & 0 & 0 & 0 \\ A_2 & -B_2 C_{bs} & A_3 & A_4 & A_5 \\ 0 & 0 & 0 & A_7 & 0 \\ 0 & 0 & 0 & A_8 & A_9 \end{bmatrix}$$

which is a block upper triangular matrix and, thus, its eigenvalues are those of the matrix

$$\begin{bmatrix} A_1 & -B_1C_{bs} & 0 \\ B_{bs}C_1 & A_{bs} & 0 \\ A_2 & -B_2C_{bs} & A_3 \end{bmatrix} \tag{S49}$$

together with those of the matrix

$$\begin{bmatrix} A_7 & 0 \\ A_8 & A_9 \end{bmatrix} \tag{S50}$$

The matrix in Eq. (S49) is block lower triangular, so its eigenvalues are those of $A_7$, together with those of $A_9$. Similarly, the matrix in Eq. (S50) is block lower triangular, so its eigenvalues are those of $A_3$, together with those of

$$\begin{bmatrix} A_1 & -B_1C_{bs} \\ B_{bs}C_1 & A_{bs} \end{bmatrix} \tag{S51}$$

Note that the closed-loop dynamics of the negative feedback interconnection of the minimal realization in Eqs.(S47)-(S48) and the subsystem given by Eqs.(S41)–(S42) can be described as $\dot{\hat{y}}_{\mathrm{cl}} = \hat{A}\hat{y}$, with $\hat{y} = (y_I, m)$ and $\hat{A}$ given by Eq.(S51). The eigenvalues of $\hat{A}$ are therefore in the open left half-plane. At the same time, due to the stabilizability and detectability of the subsystem given by Eqs. (S41)–(S42), the eigenvalues of $A_3$, $A_7$, $A_9$ are also in the open left half-plane. Consequently, the overall closed-loop system given by Eqs. (S41)–(S44) is asymptotically stable. □

In case the process to be controlled is a bursty birth-death process, i.e. it is composed by species $Y_t$ following the reactions in CRN described by Eq. (3), then Eqs. (S41)–(S42) become:

$$\frac{dy_t}{dt} = -(p_2 + k_f X^*)y + Y^* u_p \tag{S52}$$

$$w_p = y \tag{S53}$$

which yields:

$$H_p(s) = \frac{Y^*}{s + p_2 + k_f X^*}$$

and

$$\mathrm{Re}[H_p(j\omega)] = \frac{Y^*(p_2 + k_f X^*)}{(p_2 + k_f X^*)^2 + \omega^2}$$

As can be seen, $H_p(s)$ (a minimal realization of which is the system given by Eqs. (S52)–(S53)) is analytic in $\mathrm{Re}[s] \geq 0$ and $\mathrm{Re}[H_p(j\omega)] > 0$, for all $\omega \in (-\infty, \infty)$. Thus, $H_p(s)$ is WSPR (see Definition 2.77 in [9]).

Note also that the corresponding nonlinear closed-loop system admits the following unique steady state, which can be obtained by setting the time derivatives in Eqs. (S22)–(S25) to zero:

$$Y_t^* = \frac{ck_2p_1}{k_3k_f + k_2p_2} \tag{S54}$$

$$X^* = \frac{k_3}{k_2} \tag{S55}$$

$$Z_1^* = \frac{k_2(bk_3k_f + ck_2k_{in}p_1 + bk_2p_2)}{k_1k_3(k_3k_f + k_2p_2)} \tag{S56}$$

$$Z_2^* = \frac{k_1k_3^2(k_3k_f + k_2p_2)}{nk_2(bk_3k_f + ck_2k_{in}p_1 + bk_2p_2)} \tag{S57}$$

According to Theorem 4, the steady state described by Eqs. (S55)–(S57) is locally asymptotically stable.

#### S7.3 Fano factor

Assuming that the process to be controlled is a bursty birth-death process (see above) and for $E[Y_t(t)] > 0$, the output Fano factor is given by $FF_{Y_t}(t) = \frac{V_{Y_t}}{Y_t}$ where $V_{Y_t}$, $Y_t$ can be determined by the ODE system (see Eqs. (S1), (S4)):

$$
\begin{aligned}
\frac{dY_t}{dt} &= cp_1 - p_2Y_t - k_fXY_t, \\
\frac{dX}{dt} &= b + k_{in}Y_t - k_1XZ_1, \\
\frac{dZ_1}{dt} &= k_2X - nZ_1Z_2, \\
\frac{dZ_2}{dt} &= k_3 - nZ_1Z_2, \\
\frac{dV_{Y_t}}{dt} &= p_1c^2 + p_2Y_t - 2(p_2 + k_fX)V_{Y_t} + k_fXY_t - 2k_fV_{Y_tX}Y_t, \\
\frac{dV_{Y_tX}}{dt} &= k_{in}V_{Y_t} - (p_2 + k_fX)V_{Y_tX} - k_fV_XY_t - k_1XV_{Y_tZ_1} - k_1Z_1V_{Y_tX}, \\
\frac{dV_{Y_tZ_1}}{dt} &= k_2V_{Y_tX} - (p_2 + k_fX)V_{Y_tZ_1} - k_fV_{XZ_1}Y_t - nZ_1V_{Y_tZ_2} - nZ_2V_{Y_tZ_1}, \\
\frac{dV_{Y_tZ_2}}{dt} &= -(p_2 + k_fX)V_{Y_tZ_2} - k_fV_{XZ_2}Y_t - nZ_1V_{Y_tZ_2} - nZ_2V_{Y_tZ_1}, \\
\frac{dV_X}{dt} &= b + k_{in}Y_t + 2k_{in}V_{Y_tX} + k_1XZ_1 - 2k_1XV_{XZ_1} - 2k_1Z_1V_X, \\
\frac{dV_{XZ_1}}{dt} &= k_2V_X + k_{in}V_{Y_tZ_1} - k_1XV_{Z_1} - k_1Z_1V_{XZ_1} - nZ_1V_{XZ_2} - nZ_2V_{XZ_1}, \\
\frac{dV_{XZ_2}}{dt} &= k_{in}V_{Y_tZ_2} - k_1XV_{Z_1Z_2} - k_1Z_1V_{XZ_2} - nZ_1V_{XZ_2} - nZ_2V_{XZ_1}, \\
\frac{dV_{Z_1}}{dt} &= k_2X + 2k_2V_{XZ_1} + nZ_1Z_2 - 2nZ_2V_{Z_1} - 2nZ_1V_{Z_1Z_2}, \\
\frac{dV_{Z_1Z_2}}{dt} &= k_2V_{XZ_2} + nZ_1Z_2 - nZ_1V_{Z_2} - nZ_2V_{Z_1} - nZ_1V_{Z_1Z_2} - nZ_2V_{Z_1Z_2}, \\
\frac{dV_{Z_2}}{dt} &= k_3 + nZ_1Z_2 - 2nZ_1V_{Z_2} - 2nZ_2V_{Z_1Z_2}.
\end{aligned}
$$

At steady state and for $n \to \infty$, we have:

$$
FF_{Y_t}^* = 1 + \frac{\lambda_1}{\lambda_2}
$$

where

$$
\begin{aligned}
\lambda_1 = {} & b^2(-1+c)k_2^2(k_3k_f+k_2p_2)^4 \\
& + b(k_3k_f+k_2p_2)\Big(k_1k_2^2k_3^2\big(k_3k_f^2((-1+c)k_3+2cp_1)+2(-1+c)k_2k_3k_fp_2+(-1+c)k_2^2p_2^2\big) \\
& \quad + (k_3k_f+k_2p_2)\big((-1+c)k_3^4k_f^3+2ck_2^2k_3k_f(k_3k_f+(-2+c)k_2k_{in})p_1 \\
& \qquad + (-1+c)k_2(3k_3^3k_f^2+2ck_2^3k_{in}p_1)p_2+3(-1+c)k_2^2k_3^2k_fp_2^2+(-1+c)k_2^3k_3p_2^3\big)\Big) \\
& + cp_1\Big(k_2k_{in}(k_3k_f+k_2p_2)\big(2(-1+c)k_3^4k_f^3+ck_2^2k_3k_f(2k_3k_f+(-3+c)k_2k_{in})p_1 \\
& \quad + (-1+c)k_2(5k_3^3k_f^2+ck_2^3k_{in}p_1)p_2+4(-1+c)k_2^2k_3^2k_fp_2^2+(-1+c)k_2^3k_3p_2^3\big) \\
& \quad + k_1k_3^2\big(2k_3^4k_f^4+4k_2k_3^3k_f^3p_2+(-1+c)k_2^5k_{in}p_2^2 \\
& \qquad + 2k_2^3k_3k_fk_{in}(ck_fp_1+(-1+c)k_2p_2)+k_2^2k_3^2k_f^2((-1+c)k_2k_{in}+2p_2^2)\big)\Big) \in \mathbb{R}
\end{aligned}
$$

$$
\begin{aligned}
\lambda_2 = {} & 2(k_3k_f+k_2p_2)\Big(b^2k_2^2(k_3k_f+k_2p_2)^3 \\
& + b(k_3k_f+k_2p_2)\big(k_3^4k_f^3+3k_2k_3^3k_f^2p_2+2ck_2^4k_{in}p_1p_2+3k_2^2k_3^2k_fp_2^2 \\
& \quad + k_1k_2^2k_3^2(k_3k_f+k_2p_2)+k_2^3k_3(3ck_fk_{in}p_1+p_2^3)\big) \\
& + ck_2k_{in}p_1\big(k_1k_2^2k_3^2(k_3k_f+k_2p_2) \\
& \quad + (2k_3k_f+k_2p_2)(ck_2^3k_{in}p_1+k_3(k_3k_f+k_2p_2)^2)\big)\Big) \in \mathbb{R}^+
\end{aligned}
$$

### S8 Negative feedback control based on *AC-BioSD-II*

#### S8.1 Derivative control

Here, we employ the same methodology as in Section S7.1. Accordingly, we summarize only the key differences without reproducing the detailed derivations.

First, we consider the CRN in Eq. (11) together with a process to be controlled, comprising $\nu$ species $Y_1, Y_2, \ldots, Y_{\nu-1}, Y_t$. This leads to the ODE system in Eqs. (S22)–(S25), differing only in Eq. (S23). The latter now is:

$$\frac{dX}{dt} = b + k_{in} Y_t X - k_1 X Z_1 \tag{S58}$$

This results in a similar control law as in Eq. (S30), namely:

$$\tilde{u}_{CL}(s) = \underbrace{-k_p \tilde{y}_t(s)}_{\substack{\text{Proportional}\\ \text{control}}} \underbrace{-k_d \frac{s\tilde{y}_t(s)}{\varepsilon_{AB}(s^2+s)+1}}_{\substack{\text{Filtered derivative}\\ \text{control}}}$$

with $k_p = k_f X^*$, $k_d = \dfrac{k_{in} k_f Y_t^*}{k_1 k_2}$.

Subsequently, we consider the more general case (see Fig. S11b) where the feedback control action is applied on an arbitrarily chosen species, $Y_1$, interacting with the target species $Y_t$ (assuming positive process gain). For the resulting *AC-BioSD-II*-based controller, we obtain the same CRN as in Eq. (S31), except for the following structural difference: Instead of $Y_t \xrightarrow{k_{in}} Y_t + X$ we now have $Y_t + X \xrightarrow{k_{in}} Y_t + X + X$. This yields the ODE system given by Eqs. (S22)–(S25) where Eq. (S33) has now been replaced by Eq.(S58). We therefore get a similar control law as in Eq. (S40), namely:

$$\tilde{u}_{CL}(s) = \underbrace{-k_d \frac{s\tilde{y}_t(s)}{\varepsilon_{AB}(s^2+s)+1}}_{\substack{\text{Filtered derivative}\\ \text{control}}}$$

with $k_d = \dfrac{k_{in} k_f Y_1^*}{k_1 k_2}$.

#### S8.2 Sufficient conditions for closed-loop stability

Here, we adopt the same methodology as in Section S7.2. Once again, we summarize only the key differences and omit the detailed derivations.

We can use the same closed-loop system decomposition as in Eqs. (S41)–(S44), with modifications to two matrices. Specifically, we have:

$$A_{bs} = \begin{bmatrix} -\frac{k_2 b}{k_3} & -k_1 X^* & 0 \\ k_2 & -nZ_2^* & -nZ_1^* \\ 0 & -nZ_2^* & -nZ_1^* \end{bmatrix}, \quad B_{bs} = \begin{bmatrix} k_{in} X^* \\ 0 \\ 0 \end{bmatrix} \tag{S59}$$

In addition, $A_{bs}$ is Hurwitz (see Section S4 for a detailed proof). We also have:

$$H_{bs} = C_{bs}(sI - A_{bs})^{-1} B_{bs} + D_{bs} = \frac{k_f k_{in} X^* s(s+\mu_1)}{s^3 + \mu_2 s^2 + \mu_3 s + \mu_4}$$

where $\mu_1 = n(Z_1^* + Z_2^*)$, $\mu_2 = \frac{k_2 b}{k_3} + \mu_1$, $\mu_3 = \mu_1 \frac{k_2 b}{k_3} + k_1 k_2 X^*$, $\mu_4 = nk_1 k_2 X^* Z_1^*$. Moreover:

$$\text{Re}[H_{bs}(j\omega)] = \frac{(\mu_2 - \mu_1) k_f k_{in} X^* \omega^4 + (\mu_1 \mu_3 - \mu_4) k_f k_{in} X^* \omega^2}{(-\mu_2 \omega^2 + \mu_4)^2 + \omega^2 (\mu_3 - \omega^2)^2}$$

with $\mu_2 - \mu_1 = \frac{k_2 b}{k_3} > 0$ and $\mu_1 \mu_3 - \mu_4 = \mu_1^2 \frac{k_2 b}{k_3} + nk_1 k_2 X^* Z_2^* > 0$. Consequently, Theorem 4 holds.

Finally, if the process to be controlled is a bursty birth-death process—where $Y_t$ follows the reactions in the CRN described by Eq. (3)—we obtain the following unique steady state:

$$\begin{aligned} Y_t^* &= \frac{ck_2 p_1}{k_3 k_f + k_2 p_2} \\ X^* &= \frac{k_3}{k_2} \\ Z_1^* &= \frac{k_2(bk_3 k_f + ck_3 k_{in} p_1 + bk_2 p_2)}{k_1 k_3 (k_3 k_f + k_2 p_2)} \\ Z_2^* &= \frac{k_1 k_3^3 k_f + k_1 k_2 k_3^2 p_2}{k_2 n(bk_3 k_f + ck_3 k_{in} p_1 + bk_2 p_2)} \end{aligned}$$

which is locally asymptotically stable, according to Theorem 4.

#### S8.3 Fano factor

Assuming that the process to be controlled is a bursty birth-death process (see above) and for $E[Y_t(t)] > 0$, the output Fano factor is given by $FF_{Y_t}(t) = \frac{V_{Y_t}}{Y_t}$ where $V_{Y_t}$, $Y_t$ can be determined by the ODE system (see Eqs. (S1), (S4)):

$$
\begin{aligned}
\frac{dY_t}{dt} &= cp_1 - p_2Y_t - k_fXY_t, \\
\frac{dX}{dt} &= b + k_{in}Y_tX - k_1XZ_1, \\
\frac{dZ_1}{dt} &= k_2X - nZ_1Z_2, \\
\frac{dZ_2}{dt} &= k_3 - nZ_1Z_2, \\
\frac{dV_{Y_t}}{dt} &= p_1c^2 + p_2Y_t - 2(p_2 + k_fX)V_{Y_t} + k_fXY_t - 2k_fY_tV_{Y_tX}, \\
\frac{dV_{Y_tX}}{dt} &= (Y_tk_{in} - k_1Z_1)V_{Y_tX} - (p_2 + k_fX)V_{Y_tX} - k_fY_tV_X - k_1XV_{Y_tZ_1} + k_{in}XV_{Y_t}, \\
\frac{dV_{Y_tZ_1}}{dt} &= k_2V_{Y_tX} - (p_2 + k_fX)V_{Y_tZ_1} - k_fY_tV_{XZ_1} - nZ_1V_{Y_tZ_2} - nZ_2V_{Y_tZ_1}, \\
\frac{dV_{Y_tZ_2}}{dt} &= -(p_2 + k_fX)V_{Y_tZ_2} - k_fY_tV_{XZ_2} - nZ_1V_{Y_tZ_2} - nZ_2V_{Y_tZ_1}, \\
\frac{dV_X}{dt} &= b + 2(Y_tk_{in} - Z_1k_1)V_X + k_{in}Y_tX + k_1XZ_1 - 2k_1XV_{XZ_1} + 2k_{in}XV_{Y_tX}, \\
\frac{dV_{XZ_1}}{dt} &= k_2V_X + (Y_tk_{in} - Z_1k_1)V_{XZ_1} - k_1XV_{Z_1} + k_{in}XV_{Y_tZ_1} - nZ_1V_{XZ_2} - nZ_2V_{XZ_1}, \\
\frac{dV_{XZ_2}}{dt} &= (Y_tk_{in} - Z_1k_1)V_{XZ_2} - k_1XV_{Z_1Z_2} + k_{in}XV_{Y_tZ_2} - nZ_1V_{XZ_2} - nZ_2V_{XZ_1}, \\
\frac{dV_{Z_1}}{dt} &= k_2X + 2k_2V_{XZ_1} + nZ_1Z_2 - 2nZ_2V_{Z_1} - 2nZ_1V_{Z_1Z_2}, \\
\frac{dV_{Z_1Z_2}}{dt} &= k_2V_{XZ_2} + nZ_1Z_2 - nZ_1V_{Z_2} - nZ_2V_{Z_1} - nZ_1V_{Z_1Z_2} - nZ_2V_{Z_1Z_2}, \\
\frac{dV_{Z_2}}{dt} &= k_3 + nZ_1Z_2 - 2nZ_1V_{Z_2} - 2nZ_2V_{Z_1Z_2}.
\end{aligned}
$$

At steady state and for $n \to \infty$, we have:

$$FF^*_{Y_t} = 1 + \frac{\lambda_1}{\lambda_2}$$

where

$$
\begin{aligned}
\lambda_1 = {} & b^2(-1+c)k_2^2(k_3k_f+k_2p_2)^3 \\
& + ck_3^3k_fp_1\Big(k_f\big(k_3^2k_f(2k_1+(-1+c)k_{in})+2ck_2^2k_{in}p_1\big) \\
& \quad + 2k_2k_3k_f(k_1+(-1+c)k_{in})p_2+(-1+c)k_2^2k_{in}p_2^2\Big) \\
& + bk_3\Big(k_1k_2^2k_3\big(k_3k_f^2((-1+c)k_3+2cp_1)+2(-1+c)k_2k_3k_fp_2 \\
& \quad + (-1+c)k_2^2p_2^2\big)+(k_3k_f+k_2p_2)\big((-1+c)k_3^3k_f^3+3(-1+c)k_2k_3^2k_f^2p_2 \\
& \quad + (-1+c)k_2^3p_2^3+k_2^2k_3k_f(2c(k_f-k_{in})p_1+3(-1+c)p_2^2)\big)\Big) \in \mathbb{R}
\end{aligned}
$$

$$
\begin{aligned}
\lambda_2 = {} & 2(k_3k_f+k_2p_2)\Big(ck_3^3k_fk_{in}p_1(k_3k_f+k_2p_2)+b^2k_2^2(k_3k_f+k_2p_2)^2 \\
& + bk_3\big(k_3^3k_f^3+3k_2k_3^2k_f^2p_2+k_2^3p_2^3+k_1k_2^2k_3(k_3k_f+k_2p_2) \\
& \quad + k_2^2k_3k_f(ck_{in}p_1+3p_2^2)\big)\Big) \in \mathbb{R}^+
\end{aligned}
$$

### S9 Positive feedback control based on *BioSD-II*

#### S9.1 Derivative control

We consider the CRN in Eq. (14) and a process to be controlled with $\nu$ species, $Y_1$, $Y_2$,..., $Y_{\nu-1}$, $Y_t$. Adopting the same notation as in Section S7.1, the dynamics of the closed-loop topology can be described by the following RRE-based model:

$$\frac{dY}{dt} = f(Y) + \xi \underbrace{(k_f X)}_{\substack{\text{control}\\ \text{input signal}}} \tag{S60}$$

$$\frac{dX}{dt} = b + k_{in}Y_t - k_1 X Z_1 \tag{S61}$$

$$\frac{dZ_1}{dt} = k_2 X - nZ_1 Z_2 \tag{S62}$$

$$\frac{dZ_2}{dt} = k_3 - nZ_1 Z_2 \tag{S63}$$

with $\xi = [0 \;\; 0 \;\; \dots \;\; 1]^T \in \mathbb{Z}^\nu$.

Applying Jacobian linearization on Eqs. (S60)-(S63) around an equilibrium of interest E yields:

$$\frac{dy}{dt} = Ay + \xi u_{CL} \tag{S64}$$

$$\frac{dx}{dt} = k_{in}y_t - k_1 Z_1^* x - k_1 X^* z_1 \tag{S65}$$

$$\frac{dz_1}{dt} = k_2 x - nZ_2^* z_1 - nZ_1^* z_2 \tag{S66}$$

$$\frac{dz_2}{dt} = -nZ_2^* z_1 - nZ_1^* z_2 \tag{S67}$$

where $A = \left.\frac{\partial f(Y)}{\partial Y}\right|_E$. Moreover, $u_{CL} = k_f x$, or equivalently, in the Laplace domain:

$$\tilde{u}_{CL}(s) = \underbrace{k_d \frac{s\tilde{y}_t(s)}{\varepsilon_B(s^2+s)+1}}_{\substack{\text{Filtered derivative}\\ \text{control}}} \tag{S68}$$

with $k_d = \frac{k_{in}k_f}{k_1 k_3}$.

Next, we consider the more general scenario where the feedback control action is exerted on a different, arbitrarily selected species, $Y_1$, that interacts with the target species $Y_t$ under the assumption of a positive process gain (see Fig. S12a).

The *BioSD-II*-based controller can now be described by the CRN:

$$\begin{aligned} &\varnothing \xrightarrow{\;b\;} X\,, \quad X \xrightarrow{\;k_2\;} X + Z_1\,, \quad Z_1 + X \xrightarrow{\;k_1\;} Z_1\,, \quad Z_1 + Z_2 \xrightarrow{\;n\;} \varnothing\,, \\ &\varnothing \xrightarrow{\;k_3\;} Z_2\,, \quad Y_t \xrightarrow{\;k_{in}\;} Y_t + X\,, \quad X \xrightarrow{\;k_f\;} X + Y_1 \end{aligned} \tag{S69}$$

which results in the following RREs:

$$\frac{dY}{dt} = f(Y) + \xi \underbrace{(k_f X)}_{\substack{\text{control} \\ \text{input signal}}} \tag{S70}$$

$$\frac{dX}{dt} = b + k_{in} Y_t - k_1 X Z_1 \tag{S71}$$

$$\frac{dZ_1}{dt} = k_2 X - n Z_1 Z_2 \tag{S72}$$

$$\frac{dZ_2}{dt} = k_3 - n Z_1 Z_2 \tag{S73}$$

where $\xi = [1 \ \ 0 \ \ \ldots \ \ 0]^T \in \mathbb{Z}^\nu$.

Through Jacobian linearization on Eqs. (S70)-(S73) around an equilibrium of interest E, we obtain:

$$\frac{dy}{dt} = Ay + \xi u_{CL} \tag{S74}$$

$$\frac{dx}{dt} = k_{in} y_t - k_1 Z_1^* x - k_1 X^* z_1 \tag{S75}$$

$$\frac{dz_1}{dt} = k_2 x - n Z_2^* z_1 - n Z_1^* z_2 \tag{S76}$$

$$\frac{dz_2}{dt} = -n Z_2^* z_1 - n Z_1^* z_2 \tag{S77}$$

where $A = \left.\dfrac{\partial f(Y)}{\partial Y}\right|_E$ and $u_{CL} = k_f x$. We therefore arrive at Eq.(S68).

#### S9.2 Regulating a bursty birth-death process

Here, we examine the scenario in which the process to be controlled is a bursty birth–death process; that is, it involves a single species, $Y_t$, obeying the reactions of the CRN in Eq. (3).

The closed-loop dynamics can be described by the RREs:

$$\frac{dY_t}{dt} = p_1 c - p_2 Y_t + k_f X \tag{S78}$$
$$\frac{dX}{dt} = b + k_{in} Y_t - k_1 X Z_1 \tag{S79}$$
$$\frac{dZ_1}{dt} = k_2 X - n Z_1 Z_2 \tag{S80}$$
$$\frac{dZ_2}{dt} = k_3 - n Z_1 Z_2 \tag{S81}$$

**Theorem 5**

The system defined by Eqs.(S78)-(S81) has a unique and positive steady state which is locally exponentially stable.

*Proof.* Setting the time derivatives in Eqs.(S78)-(S81 to zero yields:

$$\begin{aligned}
Y_t^* &= \frac{k_3 k_f + c k_2 p_1}{k_2 p_2} \\
X^* &= \frac{k_3}{k_2} \\
Z_1^* &= \frac{k_3 k_f k_{in} + c k_2 k_{in} p_1 + b k_2 p_2}{k_1 k_3 p_2} \\
Z_2^* &= \frac{k_1 k_3^2 p_2}{n\left(k_3 k_f k_{in} + c k_2 k_{in} p_1 + b k_2 p_2\right)}
\end{aligned}$$

Clearly, $Y_t^*, X^*, Z_1^*, Z_2^* > 0$ since $p_1, p_2, c, k_f, k_{in}, k_1, k_2, k_3, b, n > 0$.

We next apply Jacobian linearization on Eqs.(S78)-(S81 around the fix point $(Y_t^*, X^*, Z_1^*, Z_2^*)$ to obtain:

$$\begin{bmatrix} \dot{Y}_t \\ \dot{X} \\ \dot{Z}_1 \\ \dot{Z}_2 \end{bmatrix} = \underbrace{\begin{bmatrix} -p_2 & k_f & 0 & 0 \\ k_{in} & -k_1 Z_1^* & -k_1 X^* & 0 \\ 0 & k_2 & -n Z_2^* & -n Z_1^* \\ 0 & 0 & -n Z_2^* & -n Z_1^* \end{bmatrix}}_{M_3} \begin{bmatrix} Y_t \\ X \\ Z_1 \\ Z_2 \end{bmatrix}$$

The characteristic polynomial of $M_3$ is:

$$P_{M_3}(s) = \det(sI - M_3) = s^4 + \zeta_3 s^3 + \zeta_2 s^2 + \zeta_1 s + \zeta_0$$

where $\zeta_3 = n(Z_1^* + Z_2^*) + k_1 Z_1^* + p_2$, $\zeta_2 = k_1 k_3 + n k_1 Z_1^{*2} + n(Z_1^* + Z_2^*)(k_1 Z_1^* + p_2) + \dfrac{k_2(c k_{in} p_1 + b p_2)}{k_3}$,

$\zeta_1 = nk_1k_3Z_1^* + k_1k_3p_2 + (Z_1^* + Z_2^*)\dfrac{nk_2(ck_{in}p_1 + bp_2)}{k_3}$, $\zeta_0 = k_1k_3p_2nZ_1^*$.

According to the Routh–Hurwitz criterion, $P_{M_3}(s)$ has all its roots in the open left half-plane if and only if $\zeta_3, \zeta_2, \zeta_1, \zeta_0 > 0$, $\zeta_2\zeta_3 > \zeta_1$ and $\zeta_1\zeta_2\zeta_3 > \zeta_1^2 + \zeta_3^2\zeta_0$. The first condition clearly holds. The second condition holds because:

$$\zeta_2\zeta_3 - \zeta_1 > 0$$

or

$$\begin{aligned}\frac{1}{k_3}\Big(&ck_2k_{in}p_1(p_2 + k_1Z_1^*) + bk_2p_2(p_2 + k_1Z_1^*) + k_3\big(np_2(Z_1^* + Z_2^*)(p_2 + n(Z_1^* + Z_2^*))\\ &+ k_1^2Z_1^*(k_3 + nZ_1^*(2Z_1^* + Z_2^*)) + k_1n(k_3Z_2^* + nZ_1^*(Z_1^* + Z_2^*)(2Z_1^* + Z_2^*) + p_2Z_1^*(3Z_1^* + 2Z_2^*))\big)\Big) > 0\end{aligned} \tag{S82}$$

which is always true. The third condition also holds because:

$$\zeta_1\zeta_2\zeta_3 - \zeta_1^2 - \zeta_3^2\zeta_0 > 0$$

or

$$\begin{aligned}\frac{1}{k_3^2}\Bigg(&c^2k_2^2k_{in}^2np_1^2(p_2 + k_1Z_1^*)(Z_1^* + Z_2^*) + b^2k_2^2np_2^2(p_2 + k_1Z_1^*)(Z_1^* + Z_2^*)\\ &+ bk_2k_3p_2\Big(n^2p_2(Z_1^* + Z_2^*)^2(p_2 + n(Z_1^* + Z_2^*)) + k_1^2Z_1^*\big(n^2Z_1^*(Z_1^* + Z_2^*)(2Z_1^* + Z_2^*) + k_3(p_2 + 2nZ_1^* + nZ_2^*)\big)\\ &\quad + k_1\big(k_3(p_2^2 + np_2Z_1^* + n^2Z_2^*(Z_1^* + Z_2^*)) + n^2Z_1^*(Z_1^* + Z_2^*)\big(n(Z_1^* + Z_2^*)(2Z_1^* + Z_2^*) + p_2(3Z_1^* + 2Z_2^*)\big)\big)\Big)\\ &+ ck_2k_{in}p_1\Big(k_1^2k_3Z_1^*\big(n^2Z_1^*(Z_1^* + Z_2^*)(2Z_1^* + Z_2^*) + k_3(p_2 + 2nZ_1^* + nZ_2^*)\big)\\ &\quad + np_2(Z_1^* + Z_2^*)\big(2bk_2p_2 + k_3n(Z_1^* + Z_2^*)(p_2 + n(Z_1^* + Z_2^*))\big)\\ &\quad + k_1\big(2bk_2np_2Z_1^*(Z_1^* + Z_2^*) + k_3^2(p_2^2 + np_2Z_1^* + n^2Z_2^*(Z_1^* + Z_2^*))\\ &\qquad + k_3n^2Z_1^*(Z_1^* + Z_2^*)\big(n(Z_1^* + Z_2^*)(2Z_1^* + Z_2^*) + p_2(3Z_1^* + 2Z_2^*)\big)\big)\Big)\\ &+ k_1k_3^3\Big(np_2^2Z_2^*(p_2 + n(Z_1^* + Z_2^*)) + k_1^2Z_1^*\big(k_3(p_2 + nZ_1^*) + nZ_1^*(p_2(Z_1^* + Z_2^*) + nZ_1^*(2Z_1^* + Z_2^*))\big)\\ &\quad + k_1n\big(p_2^2Z_1^*(Z_1^* + 2Z_2^*) + nZ_1^*\big(k_3Z_2^* + nZ_1^*(Z_1^* + Z_2^*)(2Z_1^* + Z_2^*)\big)\\ &\qquad + p_2\big(k_3Z_2^* + nZ_1^*(3Z_1^{*2} + 3Z_1^*Z_2^* + Z_2^{*2})\big)\big)\Big)\Bigg) > 0\end{aligned}$$

which is always true.

Consequently, $M_3$ is Hurwitz and the steady state $(Y_t^*, X^*, Z_1^*, Z_2^*)$ is locally exponentially stable.

□

#### S9.3 Fano factor

For the closed-loop system under investigation in Section S9.2 and assuming $E[Y_t(t)] > 0$, the output Fano factor is given by $FF_{Y_t}(t) = \frac{V_{Y_t}}{Y_t}$ where $V_{Y_t}$, $Y_t$ can be determined by the ODE system (see Eqs. (S1), (S4)):

$$
\begin{aligned}
\frac{dY_t}{dt} &= cp_1 - p_2Y_t + k_fX,\\
\frac{dX}{dt} &= b + k_{in}Y_t - k_1XZ_1,\\
\frac{dZ_1}{dt} &= k_2X - nZ_1Z_2,\\
\frac{dZ_2}{dt} &= k_3 - nZ_1Z_2,\\
\frac{dV_{Y_t}}{dt} &= p_1c^2 + k_fX + p_2Y_t + 2k_fV_{Y_tX} - 2p_2V_{Y_t},\\
\frac{dV_{Y_tX}}{dt} &= k_fV_X + k_{in}V_{Y_t} - p_2V_{Y_tX} - k_1XV_{Y_tZ_1} - k_1Z_1V_{Y_tX},\\
\frac{dV_{Y_tZ_1}}{dt} &= k_2V_{Y_tX} + k_fV_{XZ_1} - p_2V_{Y_tZ_1} - nZ_1V_{Y_tZ_2} - nZ_2V_{Y_tZ_1},\\
\frac{dV_{Y_tZ_2}}{dt} &= k_fV_{XZ_2} - p_2V_{Y_tZ_2} - nZ_1V_{Y_tZ_2} - nZ_2V_{Y_tZ_1},\\
\frac{dV_X}{dt} &= b + k_{in}Y_t + 2k_{in}V_{Y_tX} + k_1XZ_1 - 2k_1XV_{XZ_1} - 2k_1Z_1V_X,\\
\frac{dV_{XZ_1}}{dt} &= k_2V_X + k_{in}V_{Y_tZ_1} - k_1XV_{Z_1} - k_1Z_1V_{XZ_1} - nZ_1V_{XZ_2} - nZ_2V_{XZ_1},\\
\frac{dV_{XZ_2}}{dt} &= k_{in}V_{Y_tZ_2} - k_1XV_{Z_1Z_2} - k_1Z_1V_{XZ_2} - nZ_1V_{XZ_2} - nZ_2V_{XZ_1},\\
\frac{dV_{Z_1}}{dt} &= k_2X + 2k_2V_{XZ_1} + nZ_1Z_2 - 2nZ_2V_{Z_1} - 2nZ_1V_{Z_1Z_2},\\
\frac{dV_{Z_1Z_2}}{dt} &= k_2V_{XZ_2} + nZ_1Z_2 - nZ_1V_{Z_2} - nZ_2V_{Z_1} - nZ_1V_{Z_1Z_2} - nZ_2V_{Z_1Z_2},\\
\frac{dV_{Z_2}}{dt} &= k_3 + nZ_1Z_2 - 2nZ_1V_{Z_2} - 2nZ_2V_{Z_1Z_2}.
\end{aligned}
$$

At steady state and for $n \to \infty$, we have:

$$FF_{Y_t}^* = 1 + \frac{\lambda_1}{\lambda_2} \tag{S83}$$

where

$$\lambda_1 = k_1 k_3^2 \Big( (-1+c) c k_2^3 p_1 (c k_{in} p_1 + b p_2) + k_2 k_3 k_f \big( c((-1+c)k_2 + 2k_f) k_{in} p_1 + 2 b k_f p_2 \big)$$

$$+ 2 k_3^2 k_f^2 (k_f k_{in} + p_2^2) \Big)$$

$$+ k_2 \Big( 2 k_3^3 k_f^3 k_{in} (k_{in} + p_2) + (-1+c) c k_2^3 p_1 (c k_{in} p_1 + b p_2)^2$$

$$+ k_2 k_3^2 k_f^2 \big( c(3+c) k_{in}^2 p_1 + 2 k_{in} (b + c p_1) p_2 + 2 b p_2^2 \big)$$

$$+ c k_2^2 k_3 p_1 (c k_{in} p_1 + b p_2)(2 c k_f k_{in} + (-1+c) p_2^2) \Big) \in \mathbb{R}^+$$

(Recall that $c \in \mathbb{N}^+$)

$$\lambda_2 = 2 k_2 (k_3 k_f + c k_2 p_1) \Big( k_1 k_3^2 (k_3 k_f k_{in} + c k_2 k_{in} p_1 + b k_2 p_2)$$

$$+ k_2 (c k_{in} p_1 + b p_2)(k_3 k_f k_{in} + c k_2 k_{in} p_1 + b k_2 p_2 + k_3 p_2^2) \Big) \in \mathbb{R}^+$$

### S10 Positive feedback control based on *AC-BioSD-II*

#### S10.1 Derivative control

Here, we follow the same methodology described in Section S9.1, and therefore present only the key differences here, omitting the detailed derivations.

Considering the CRN in Eq. (17) and a process to be controlled with $\nu$ species, $Y_1$, $Y_2$,..., $Y_{\nu-1}$, $Y_t$, yields the ODE system in Eqs. (S60)–(S63), with the only modification appearing in Eq. (S61), which now takes the form of Eq. (S58). Consequently, we obtain a similar control law as in Eq. (S68), namely:

$$\tilde{u}_{CL}(s) = \underbrace{k_d \frac{s\tilde{y}_t(s)}{\varepsilon_{AB}(s^2+s)+1}}_{\text{Filtered derivative control}} \tag{S84}$$

with $k_d = \dfrac{k_{in}k_f}{k_1k_2}$.

We now focus on the more general scenario (see Fig. S12b) where where the feedback control action is applied on an arbitrarily chosen species, $Y_1$, interacting with the target species Yt (assuming positive process gain). The *AC-BioSD-II*-based controller is therefore described by the CRN in Eq. (S69) where $Y_t \xrightarrow{k_{in}} Y_t + X$ has now been replaced by $Y_t + X \xrightarrow{k_{in}} Y_t + X + X$. We thus obtain the ODE system given by Eqs. (S70)–(S73) where Eq.(S71) takes the form of Eq. (S68). This results in the control law given by Eq. (S84).

#### S10.2 Regulating a bursty birth-death process

We now assume a bursty birth-death process (where $Y_t$ follows the reactions in the CRN described by Eq. (3)) as the process to be controlled. The resulting closed-loop dynamics can be described by the RREs:

$$\frac{dY_t}{dt} = p_1c - p_2Y_t + k_fX \tag{S85}$$

$$\frac{dX}{dt} = b + k_{in}Y_tX - k_1XZ_1 \tag{S86}$$

$$\frac{dZ_1}{dt} = k_2X - nZ_1Z_2 \tag{S87}$$

$$\frac{dZ_2}{dt} = k_3 - nZ_1Z_2 \tag{S88}$$

By setting the time derivatives in Eqs. (S85)–(S88) to zero, we get the following unique and positive steady state:

$$\begin{aligned}
Y_t^* &= \frac{k_3 k_f + c k_2 p_1}{k_2 p_2} \\
X^* &= \frac{k_3}{k_2} \\
Z_1^* &= \frac{k_3 k_f k_{in} + c k_2 k_{in} p_1 + b k_2 p_2}{k_1 k_3 p_2} \\
Z_2^* &= \frac{k_1 k_3^2 p_2}{n\left(k_3 k_f k_{in} + c k_2 k_{in} p_1 + b k_2 p_2\right)}
\end{aligned}$$

We are only interested in cases where the closed-loop dynamics is asymptotically stable, which is not always true here. To identify the parameter regimes of local asymptotic stability regarding the steady state $(Y_t^*, X^*, Z_1^*, Z_2^*)$, one could employ the Routh-Hurwitz criterion as in Section S9.2.

More specifically, by applying Jacobian linearization on Eqs. (S85)–(S88) around the fixed point $(Y_t^*, X^*, Z_1^*, Z_2^*)$, we have:

$$\begin{bmatrix} \dot{Y}_t \\ \dot{X} \\ \dot{Z}_1 \\ \dot{Z}_2 \end{bmatrix} = \underbrace{\begin{bmatrix} -p_2 & k_f & 0 & 0 \\ k_{in} X^* & k_{in} Y_t^* - k_1 Z_1^* & -k_1 X^* & 0 \\ 0 & k_2 & -n Z_2^* & -n Z_1^* \\ 0 & 0 & -n Z_2^* & -n Z_1^* \end{bmatrix}}_{M_4} \begin{bmatrix} Y_t \\ X \\ Z_1 \\ Z_2 \end{bmatrix}$$

The characteristic polynomial of $M_4$ is:

$$P_{M_4}(s) = \det(sI - M_4) = s^4 + \zeta_3 s^3 + \zeta_2 s^2 + \zeta_1 s + \zeta_0 \tag{S89}$$

where $\zeta_3 = n(Z_1^* + Z_2^*) - k_{in} Y_t^* + k_1 Z_1^* + p_2$, $\zeta_2 = -\frac{k_3 k_f k_{in}}{k_2} + n p_2 (Z_1^* + Z_2^*) - k_{in} Y_t^* (p_2 + n(Z_1^* + Z_2^*)) + k_1 (k_3 + Z_1^* (p_2 + n(Z_1^* + Z_2^*)))$, $\zeta_1 = k_1 k_3 (p_2 + n Z_1^*) - \frac{k_{in} n (k_3 k_f + k_2 p_2 Y_t^*)(Z_1^* + Z_2^*)}{k_2} + k_1 n p_2 Z_1^* (Z_1^* + Z_2^*)$, $\zeta_0 = k_1 k_3 p_2 n Z_1^*$. According to the Routh–Hurwitz criterion, $P_{M_4}(s)$ has all its roots in the open left half-plane if and only if $\zeta_3, \zeta_2, \zeta_1, \zeta_0 > 0$, $\zeta_2 \zeta_3 > \zeta_1$ and $\zeta_1 \zeta_2 \zeta_3 > \zeta_1^2 + \zeta_3^2 \zeta_0$. This entails that $M_4$ is Hurwitz and the steady state $(Y_t^*, X^*, Z_1^*, Z_2^*)$ is locally exponentially stable.

#### S10.3 Fano factor

For the closed-loop studied above (see Section S10.2) and assuming $E[Y_t(t)] > 0$, the output Fano factor is given by $FF_{Y_t}(t) = \frac{V_{Y_t}}{Y_t}$ where $V_{Y_t}$, $Y_t$ can be determined by the ODE system (see Eqs. (S1), (S4)):

$$
\begin{aligned}
\frac{dY_t}{dt} &= cp_1 - p_2Y_t + k_fX,\\
\frac{dX}{dt} &= b + k_{in}Y_tX - k_1XZ_1,\\
\frac{dZ_1}{dt} &= k_2X - nZ_1Z_2,\\
\frac{dZ_2}{dt} &= k_3 - nZ_1Z_2,\\
\frac{dV_{Y_t}}{dt} &= p_1c^2 + k_fX + p_2Y_t + 2k_fV_{Y_tX} - 2p_2V_{Y_t},\\
\frac{dV_{Y_tX}}{dt} &= k_fV_X - p_2V_{Y_tX} + (k_{in}Y_t - k_1Z_1)V_{Y_tX} - k_1XV_{Y_tZ_1} + k_{in}XV_{Y_t},\\
\frac{dV_{Y_tZ_1}}{dt} &= k_2V_{Y_tX} + k_fV_{XZ_1} - p_2V_{Y_tZ_1} - nZ_1V_{Y_tZ_2} - nZ_2V_{Y_tZ_1},\\
\frac{dV_{Y_tZ_2}}{dt} &= k_fV_{XZ_2} - p_2V_{Y_tZ_2} - nZ_1V_{Y_tZ_2} - nZ_2V_{Y_tZ_1},\\
\frac{dV_X}{dt} &= b + 2(k_{in}Y_t - k_1Z_1)V_X + k_{in}Y_tX + k_1XZ_1 - 2k_1XV_{XZ_1} + 2k_{in}XV_{Y_tX},\\
\frac{dV_{XZ_1}}{dt} &= k_2V_X + (k_{in}Y_t - k_1Z_1)V_{XZ_1} - k_1XV_{Z_1} + k_{in}XV_{Y_tZ_1} - nZ_1V_{XZ_2} - nZ_2V_{XZ_1},\\
\frac{dV_{XZ_2}}{dt} &= (k_{in}Y_t - k_1Z_1)V_{XZ_2} - k_1XV_{Z_1Z_2} + k_{in}XV_{Y_tZ_2} - nZ_1V_{XZ_2} - nZ_2V_{XZ_1},\\
\frac{dV_{Z_1}}{dt} &= k_2X + 2k_2V_{XZ_1} + nZ_1Z_2 - 2nZ_2V_{Z_1} - 2nZ_1V_{Z_1Z_2},\\
\frac{dV_{Z_1Z_2}}{dt} &= k_2V_{XZ_2} + nZ_1Z_2 - nZ_1V_{Z_2} - nZ_2V_{Z_1} - nZ_1V_{Z_1Z_2} - nZ_2V_{Z_1Z_2},\\
\frac{dV_{Z_2}}{dt} &= k_3 + nZ_1Z_2 - 2nZ_1V_{Z_2} - 2nZ_2V_{Z_1Z_2}.
\end{aligned}
$$

At steady state and for $n \to \infty$, we have:

$$FF^*_{Y_t} = 1 + \frac{\lambda_1}{\lambda_2} \tag{S90}$$

where

$$\lambda_1 = b^2(-1+c)\,c k_2^4 p_1 p_2 + b k_2 k_3 \Big(k_3\big(2k_3k_f^2(k_1+k_{in}) + ck_2\big((-1+c)k_1k_2 + 2k_fk_{in}\big)p_1\big) + 2k_2k_3k_f^2p_2 + (-1+c)\,c k_2^2 p_1 p_2^2\Big) + k_3^3 k_f \Big(ck_2k_{in}p_1\big(2k_f + p_2(1-c)\big) + 2k_3k_f\big(k_fk_{in} + k_1p_2\big)\Big)$$

$$\lambda_2 = 2\big(k_3k_f + ck_2p_1\big)\Big(b^2k_2^3p_2 - k_3^3k_fk_{in}p_2 + bk_2k_3\big(k_3(k_1k_2 - k_fk_{in}) + k_2p_2^2\big)\Big)$$

**Theorem 6**

If the matrix $M_4$ is Hurwitz in the limit as $n \to \infty$, then $FF_{Y_t}^*$ in Eq. (S90) is greater than unity.

*Proof.* Since the matrix $M_4$ is Hurwitz in the limit as $n \to \infty$, for sufficiently large $n$, we have $\zeta_1\zeta_2\zeta_3 - \zeta_1^2 - \zeta_3^2\zeta_0 > 0$, where $\zeta_0$, $\zeta_1$, $\zeta_2$, $\zeta_3$ are the coefficients of the characteristic polynomial of $M_4$, $P_{M_4}(s)$, given by Eq. (S89). Consequently, $\dfrac{\zeta_1\zeta_2\zeta_3 - \zeta_1^2 - \zeta_3^2\zeta_0}{n^3} > 0$ for any finite $n$, so in the limit as $n \to \infty$ we have

$$\frac{(k_3k_{in}(k_3k_f + ck_2p_1) + bk_2^2p_2)^3(b^2k_2^3p_2 - k_3^3k_fk_{in}p_2 + bk_2k_3(k_3(k_1k_2 - k_fk_{in}) + k_2p_2^2))}{k_1^3k_2^4k_3^5p_2^3} \geq 0$$

or

$$b^2k_2^3p_2 - k_3^3k_fk_{in}p_2 + bk_2k_3(k_3(k_1k_2 - k_fk_{in}) + k_2p_2^2) \geq 0 \tag{S91}$$

(recall that all the parameters involved are positive). It thus follows that $\lambda_2 \geq 0$. We now turn our attention to $\lambda_1$ and, under the assumption that $c \in [1, \infty)$, we compute:

$$\lambda_1|_{c=1} = bk_2k_3\big(k_3(2k_3k_f^2(k_1+k_{in}) + 2k_2k_fk_{in}p_1) + 2k_2k_3k_f^2p_2\big) + k_3^3k_f\big(2k_2k_fk_{in}p_1 + 2k_3k_f(k_fk_{in} + k_1p_2)\big),$$

$$\left.\frac{\partial \lambda_1}{\partial c}\right|_{c=1} = k_2p_1\Big(2k_3^3k_f^2k_{in} + b^2k_2^3p_2 - k_3^3k_fk_{in}p_2 + bk_2k_3(k_3(k_1k_2 + 2k_fk_{in}) + k_2p_2^2)\Big),$$

$$\frac{\partial^2 \lambda_1}{\partial c^2} = 2k_2p_1\Big(b^2k_2^3p_2 - k_3^3k_fk_{in}p_2 + bk_2^2k_3(k_1k_3 + p_2^2)\Big)$$

We can immediately see that $\lambda_1|_{c=1} > 0$. In addition, $\left.\dfrac{\partial \lambda_1}{\partial c}\right|_{c=1} > 0$ because:

$$\begin{aligned} &2k_3^3k_f^2k_{in} + b^2k_2^3p_2 - k_3^3k_fk_{in}p_2 + bk_2k_3\left(k_3(k_1k_2 + 2k_fk_{in}) + k_2p_2^2\right) \\ =&2k_3^3k_f^2k_{in} + 3bk_2k_3^2k_fk_{in} + b^2k_2^3p_2 - k_3^3k_fk_{in}p_2 + bk_2k_3\left(k_3(k_1k_2 - k_fk_{in}) + k_2p_2^2\right) > 0 \end{aligned} \tag{S92}$$

which is true due to the condition given by Eq. (S91). Finally, $\dfrac{\partial^2 \lambda_1}{\partial c^2} > 0$ because:

$$\begin{aligned} &b^2k_2^3p_2 - k_3^3k_fk_{in}p_2 + bk_2^2k_3(k_1k_3 + p_2^2) \\ =&bk_2k_3^2k_fk_{in} + b^2k_2^3p_2 - k_3^3k_fk_{in}p_2 + bk_2k_3\left(k_3(k_1k_2 - k_fk_{in}) + k_2p_2^2\right) > 0 \end{aligned}$$

which is also true due to the condition given by Eq. (S91).

Since $\lambda_1$ is shown to be strictly increasing in $c \in [1,\infty)$ with $\lambda_1|_{c=1} > 0$, it follows that $\lambda_1 > 0$ for $c \in \mathbb{N}^+$ (the latter represents the realistic parameter regime of interest). Consequently, $\dfrac{\lambda_1}{\lambda_2} \in (0,+\infty]$ and, therefore, $FF^*_{Y_t}$ is always greater than unity. $\square$

### S11 Supplementary figures

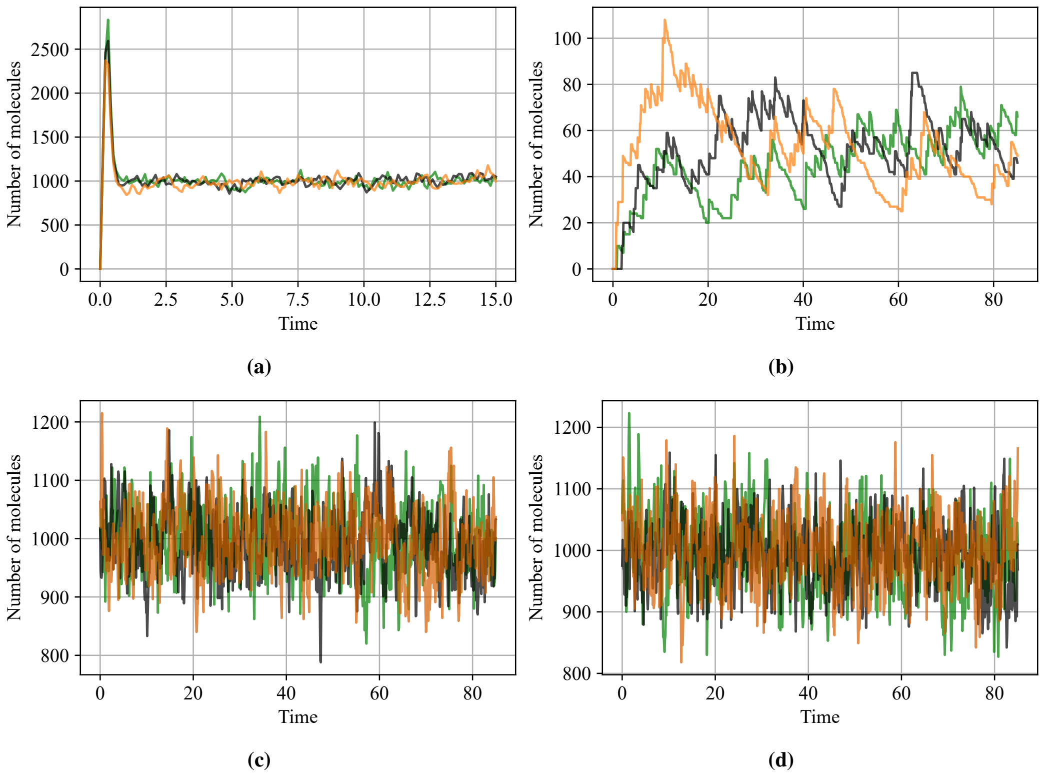

**Figure S1: Stochastic trajectories.** Panels (a)–(d) each display three individual stochastic trajectories for the simulation scenarios in Fig. 2(a,b), (c,d), (e,f), and (g,h), respectively

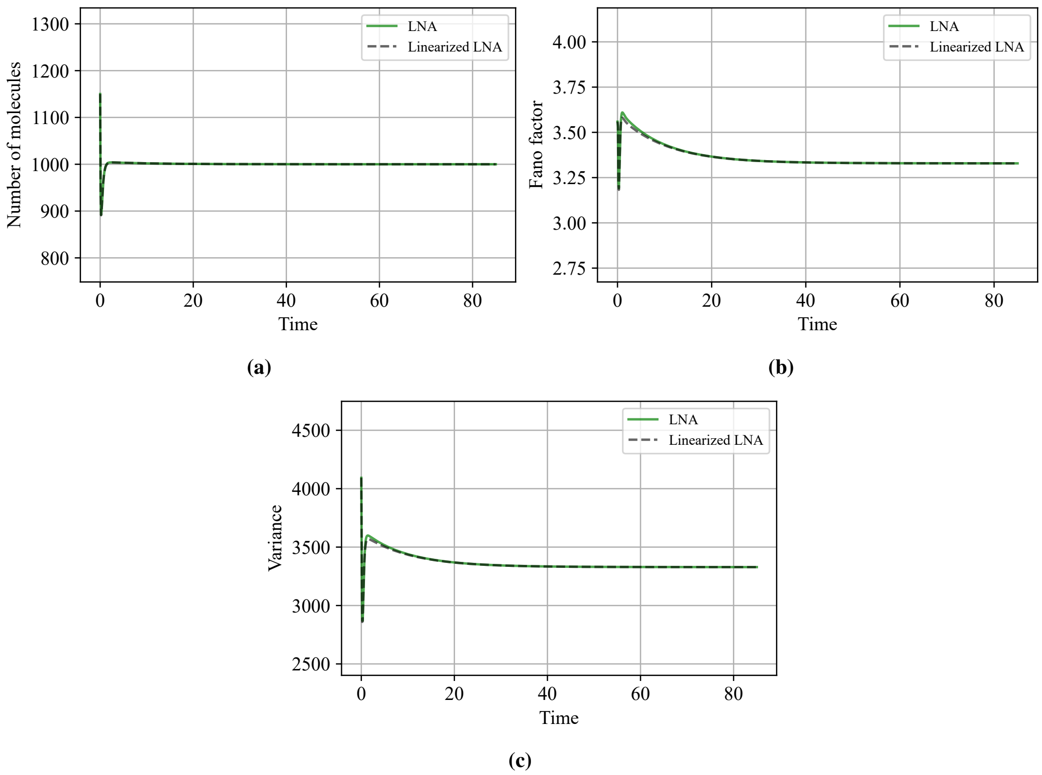

**Figure S2: Linear approximation of local dynamics.** Numerical comparison between an LNA-based nonlinear model and its linearized approximation about an equilibrium of interest, focusing on the output (a) mean, (b) variance, and (c) Fano factor of the *BioSD-II* module, as detailed in Section S5. This corresponds to the simulation scenario shown in Fig. 2(e,f), using the same parameter values except for $c = 1$ and initial conditions perturbed by 15% relative to those used in Fig. 2.

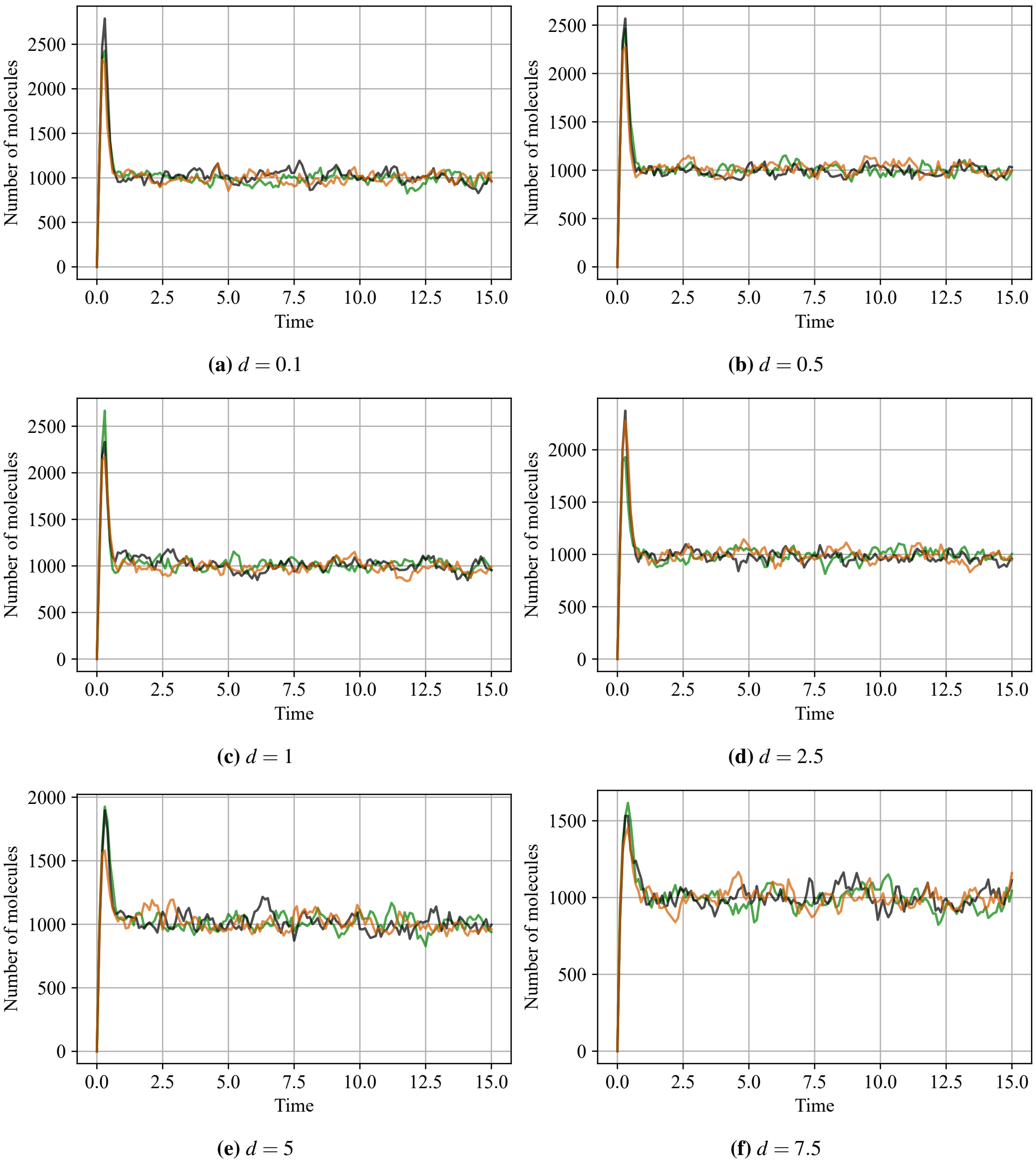

**(a)** $d = 0.1$ **(b)** $d = 0.5$

**(c)** $d = 1$ **(d)** $d = 2.5$

**(e)** $d = 5$ **(f)** $d = 7.5$

**Figure S3: Stochastic trajectories.** Panels (a)–(f) each display three individual stochastic trajectories for the simulation scenarios in Fig. 3(a)–(f), respectively.

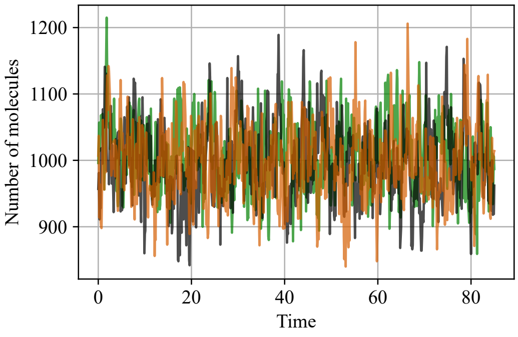

**(a)** $d = 0.1$

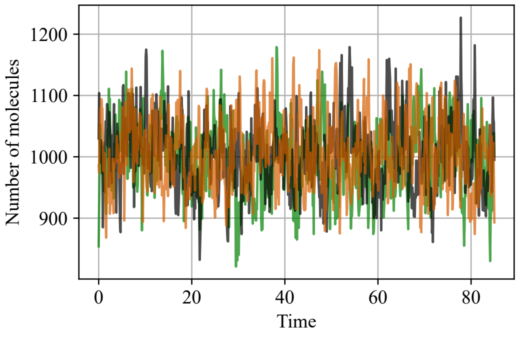

**(b)** $d = 0.5$

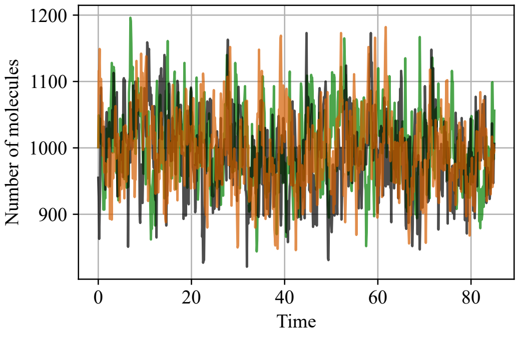

**(c)** $d = 1$

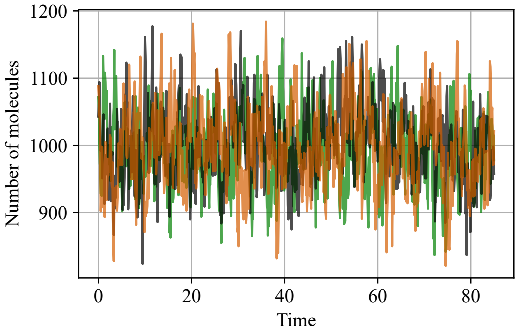

**(d)** $d = 2.5$

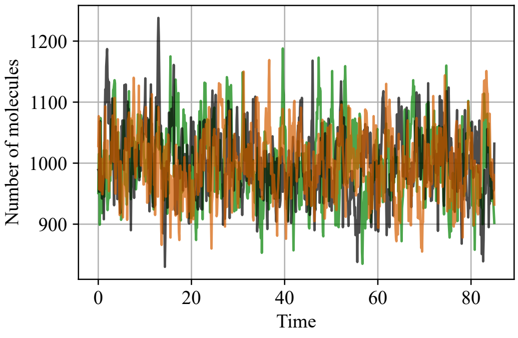

**(e)** $d = 5$

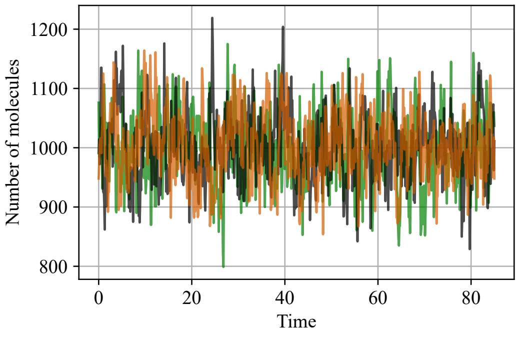

**(f)** $d = 7.5$

**Figure S4: Stochastic trajectories.** Panels (a)–(f) each display three individual stochastic trajectories for the simulation scenarios in Fig. 4(a)–(f), respectively.

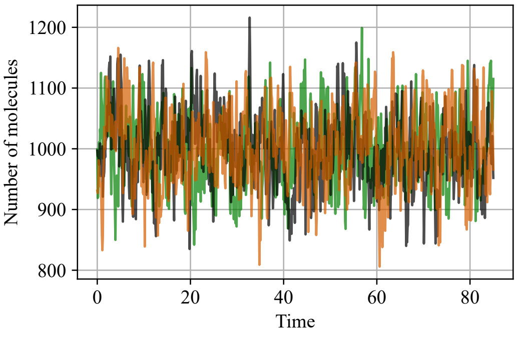

**(a)** $d = 0.1$

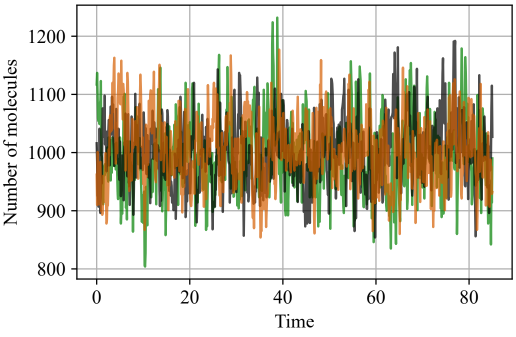

**(b)** $d = 0.5$

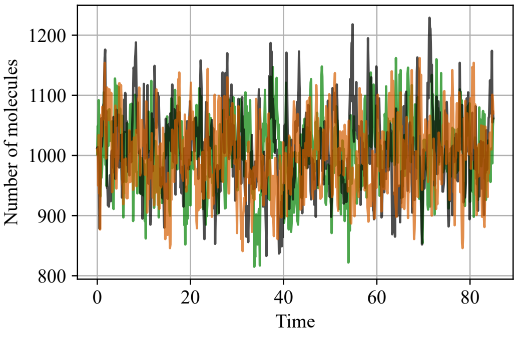

**(c)** $d = 1$

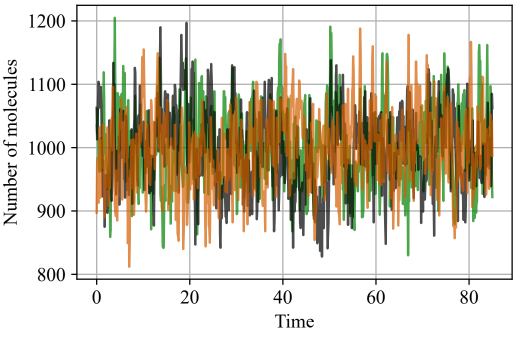

**(d)** $d = 2.5$

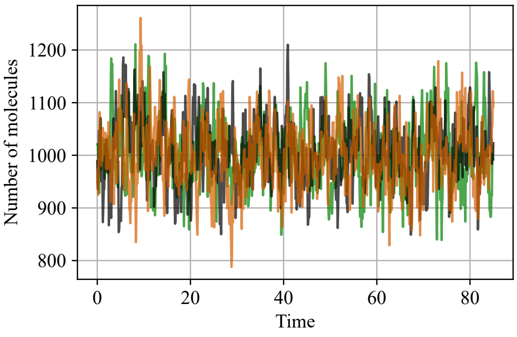

**(e)** $d = 5$

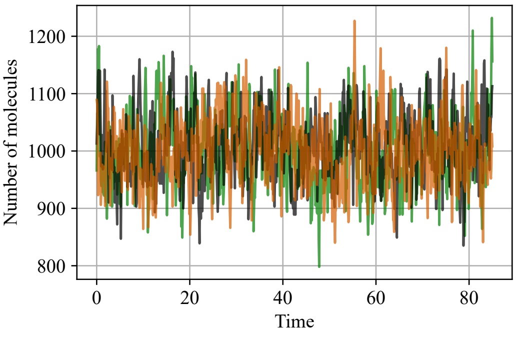

**(f)** $d = 7.5$

**Figure S5: Stochastic trajectories.** Panels (a)–(f) each display three individual stochastic trajectories for the simulation scenarios in Fig. 5(a)–(f), respectively.

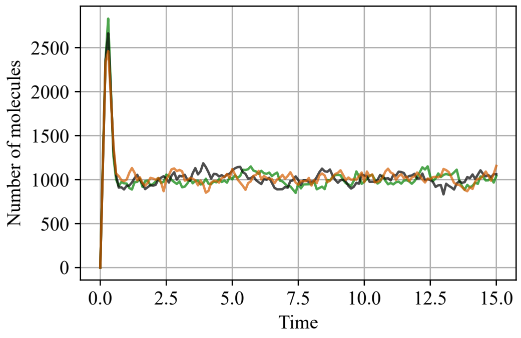

**(a)** $d = 0.1$

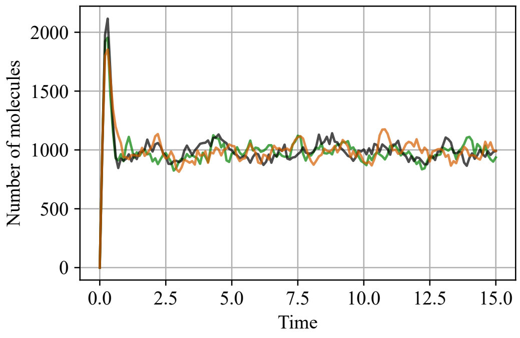

**(b)** $d = 0.5$

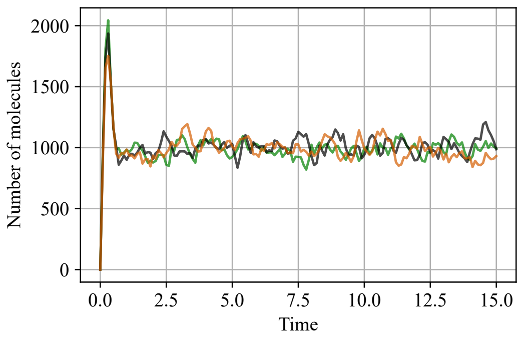

**(c)** $d = 1$

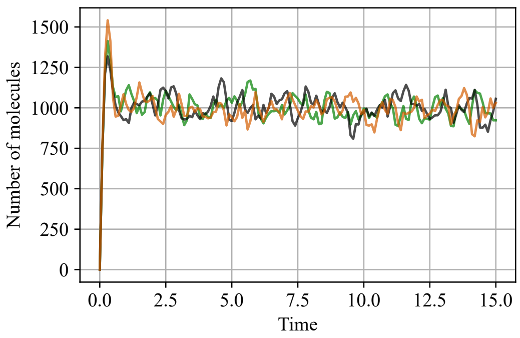

**(d)** $d = 2.5$

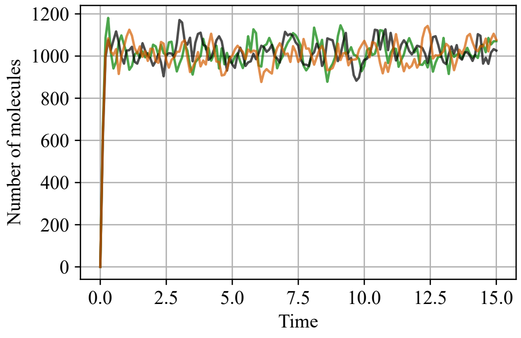

**(e)** $d = 5$

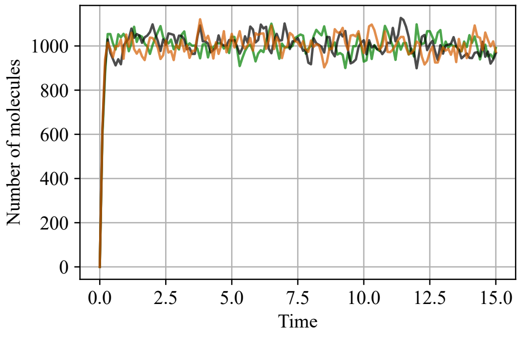

**(f)** $d = 7.5$

**Figure S6: Stochastic trajectories.** Panels (a)–(f) each display three individual stochastic trajectories for the simulation scenarios in Fig. 6(a)–(f), respectively.

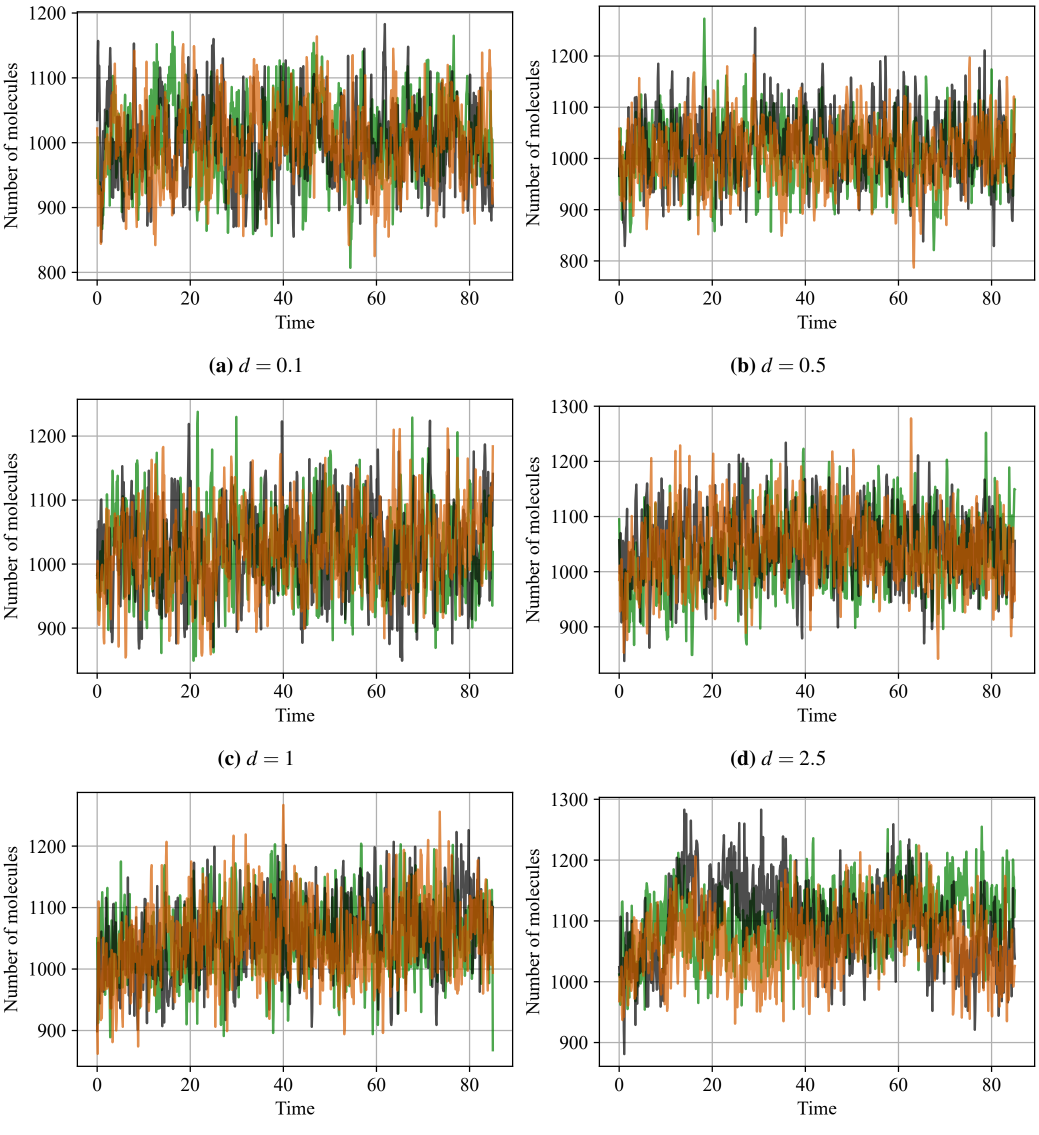

**(a)** $d = 0.1$

**(b)** $d = 0.5$

**(c)** $d = 1$

**(d)** $d = 2.5$

**(e)** $d = 5$

**(f)** $d = 7.5$

**Figure S7: Stochastic trajectories.** Panels (a)–(f) each display three individual stochastic trajectories for the simulation scenarios in Fig. 7(a)–(f), respectively.

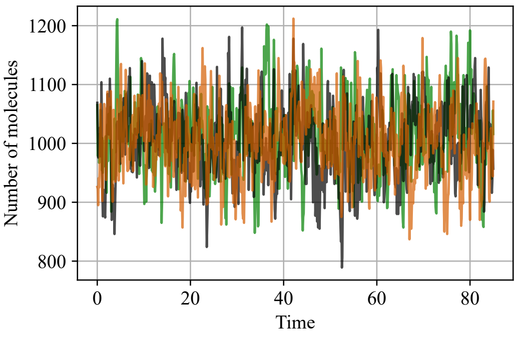

**(a)** $d = 0.1$

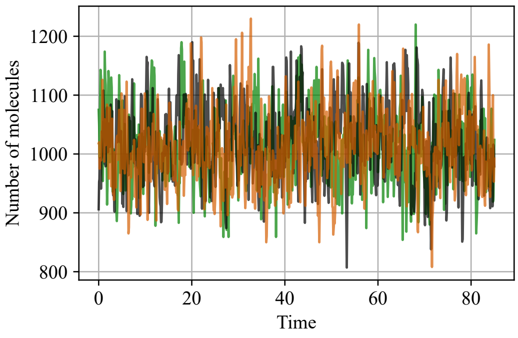

**(b)** $d = 0.5$

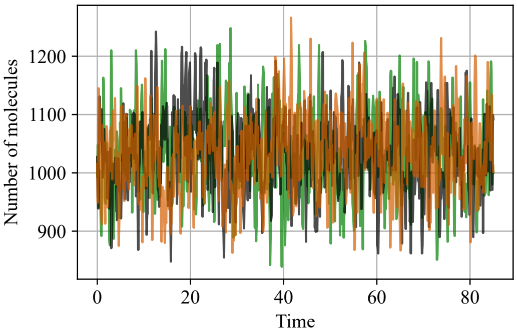

**(c)** $d = 1$

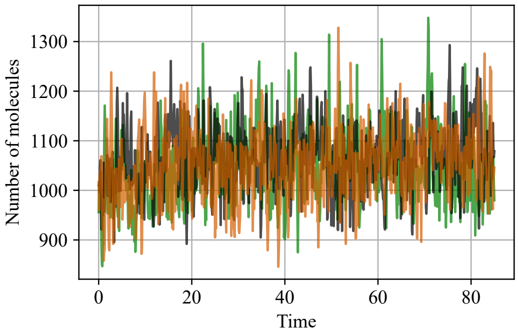

**(d)** $d = 2.5$

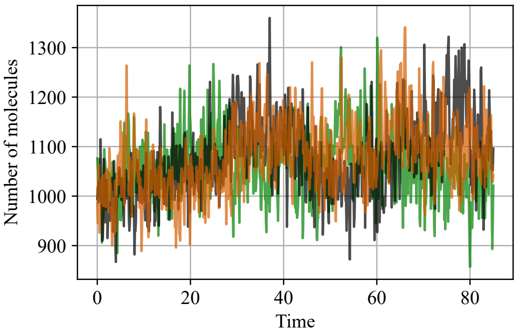

**(e)** $d = 5$

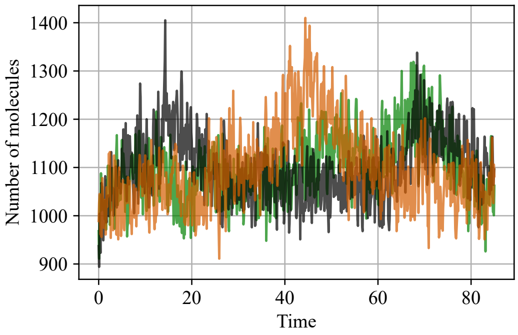

**(f)** $d = 7.5$

**Figure S8: Stochastic trajectories.** Panels (a)–(f) each display three individual stochastic trajectories for the simulation scenarios in Fig. 8(a)–(f), respectively.

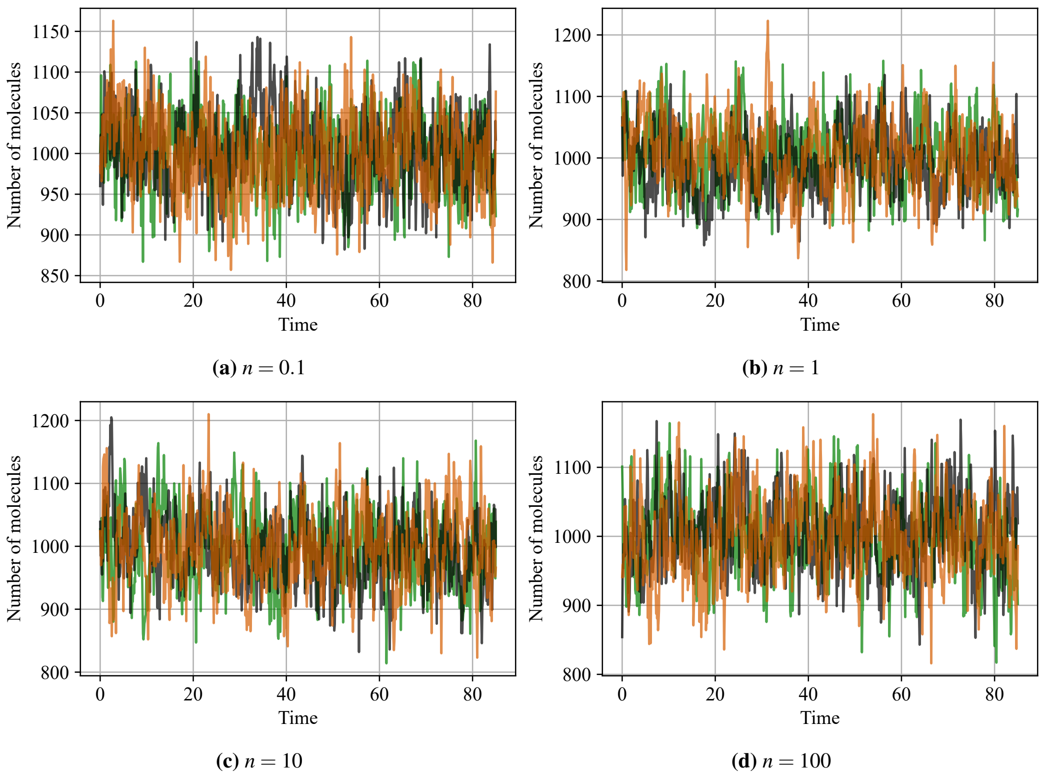

**(a)** $n = 0.1$ **(b)** $n = 1$

**(c)** $n = 10$ **(d)** $n = 100$

**Figure S9: Stochastic trajectories.** Panels (a)–(d) each display three individual stochastic trajectories for the simulation scenarios in Fig. 9(a)–(d), respectively.

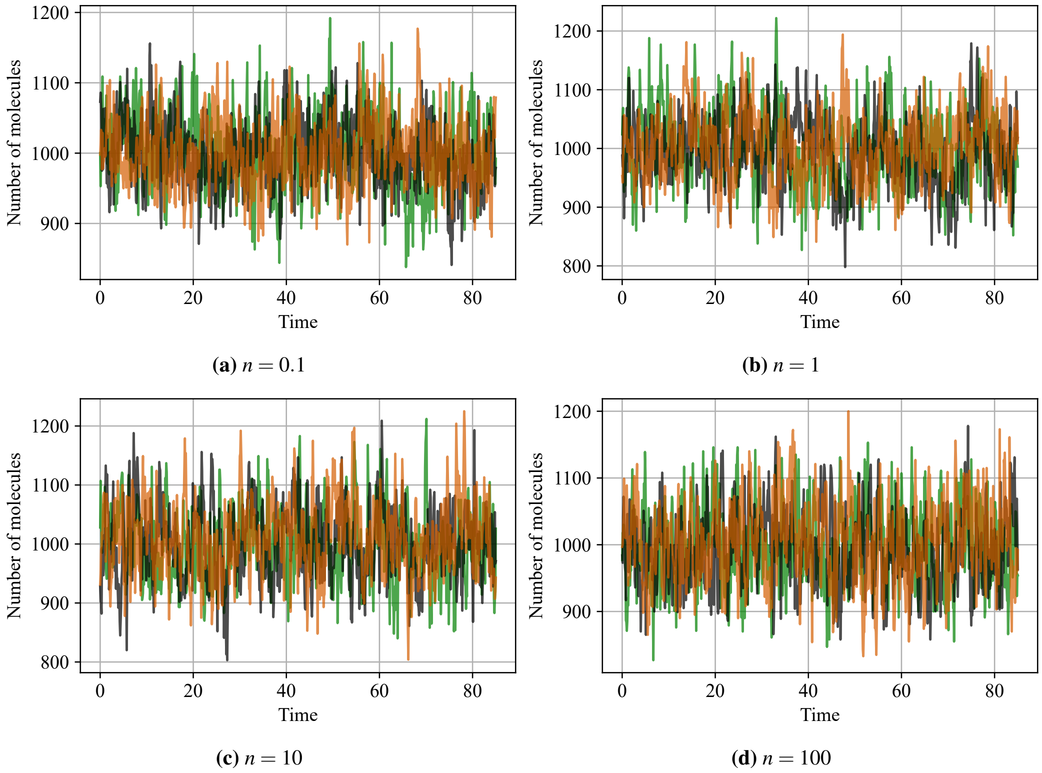

**(a)** $n = 0.1$ **(b)** $n = 1$

**(c)** $n = 10$ **(d)** $n = 100$

**Figure S10: Stochastic trajectories.** Panels (a)–(d) each display three individual stochastic trajectories for the simulation scenarios in Fig. 10(a)–(d), respectively.

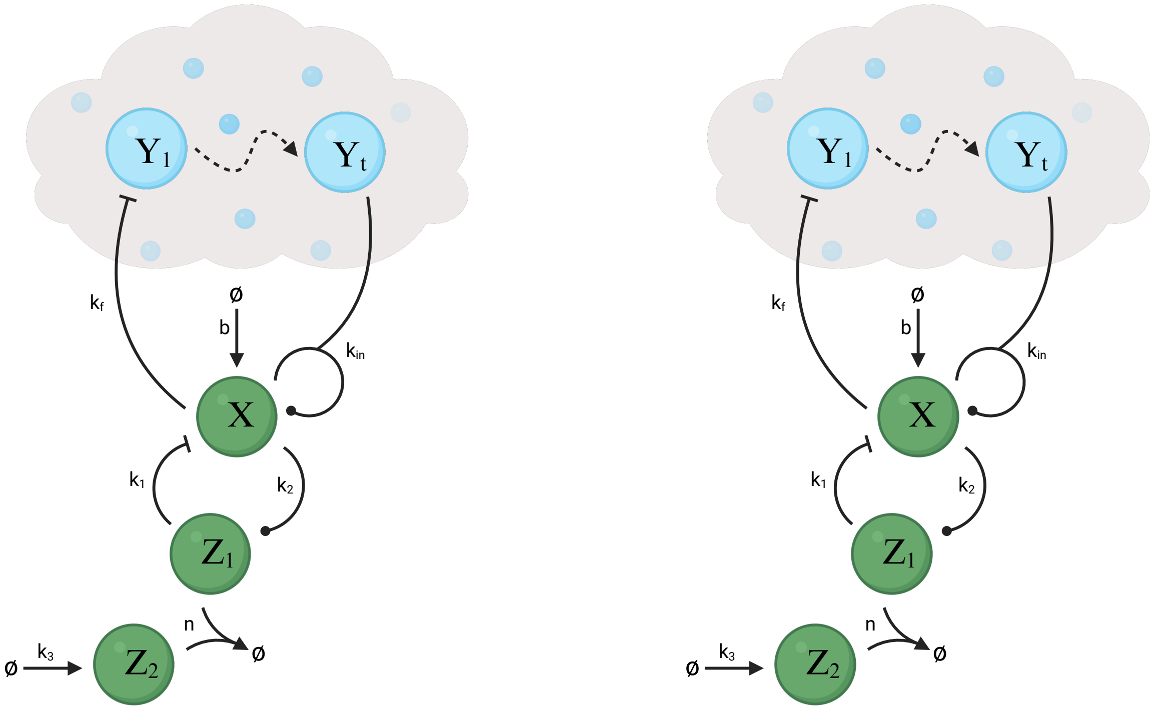

(a) (b)

**Figure S11: Biomolecular topologies exhibiting negative-feedback, derivative-control action.** Schematic of (a) a *BioSD-II*-based controller; (b) an *AC-BioSD-II*-based controller. The required measurement and actuation reactions are implemented via different species of the process being controlled. A positive process gain is assumed (represented by a dashed arrow).

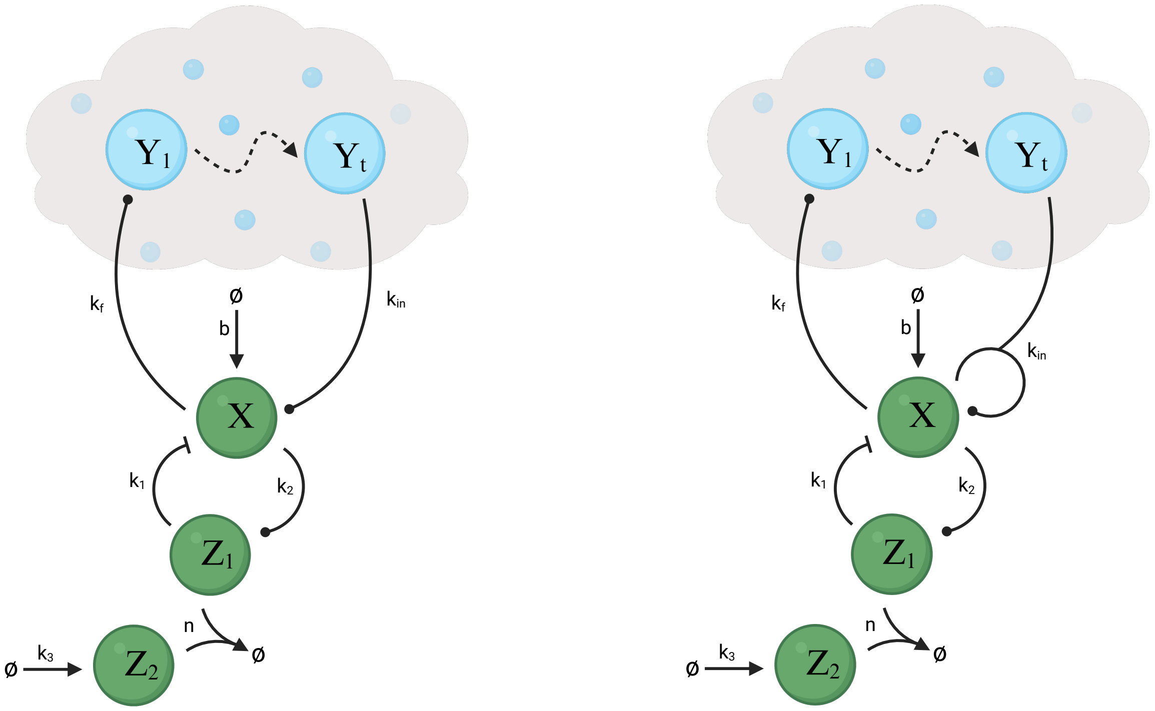

(a) (b)

**Figure S12: Biomolecular topologies exhibiting positive-feedback, derivative-control action.** Schematic of (a) a *BioSD-II*-based controller; (b) an *AC-BioSD-II*-based controller. The required measurement and actuation reactions are implemented via different species of the process being controlled. A positive process gain is assumed (represented by a dashed arrow).

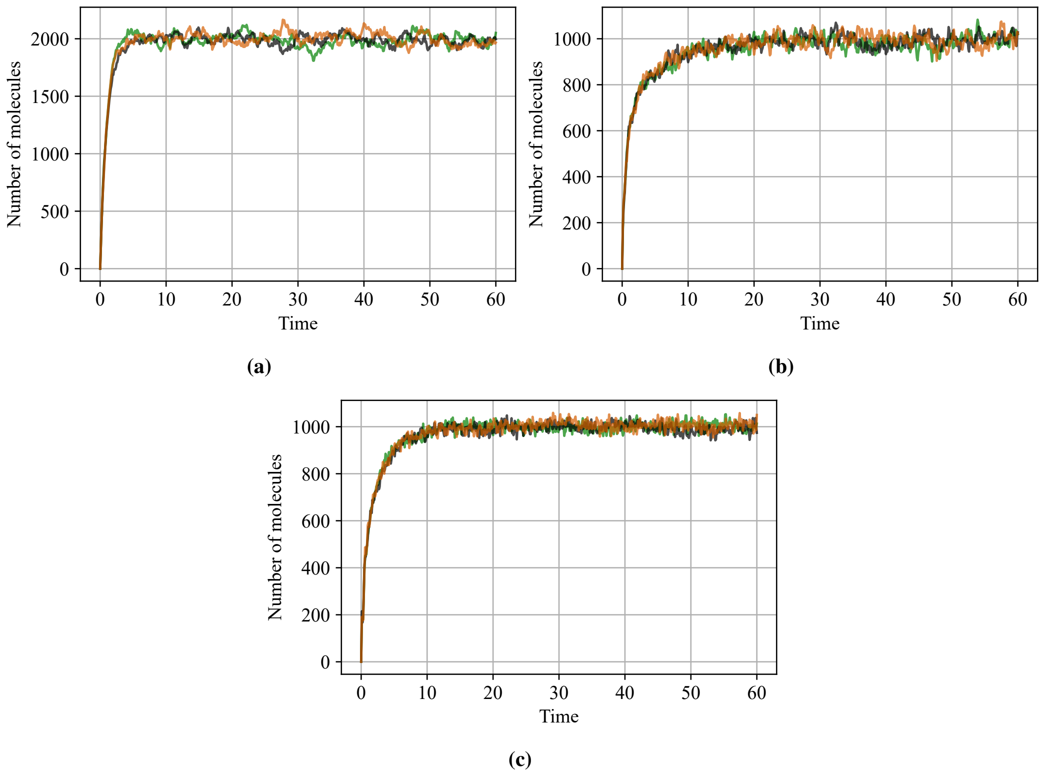

**Figure S13: Stochastic trajectories.** Panels (a)–(c) each display three individual stochastic trajectories for the simulation scenarios in Fig. 13(a,b), (c,d) and (e,f), respectively

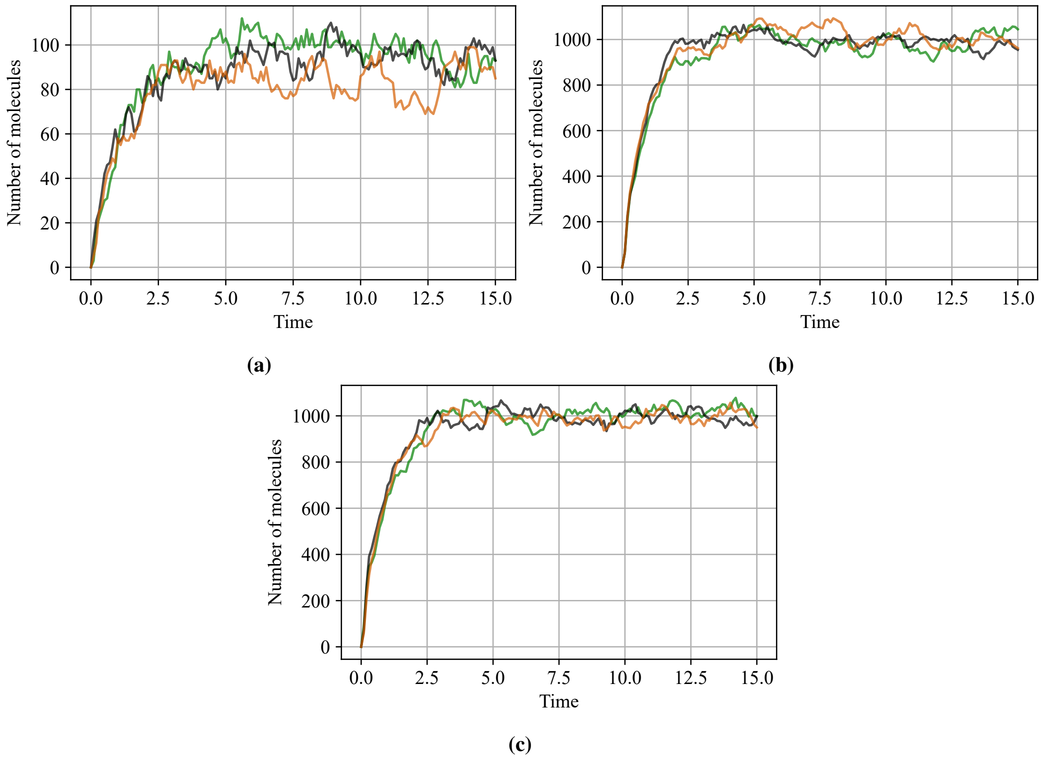

**Figure S14: Stochastic trajectories.** Panels (a)–(c) each display three individual stochastic trajectories for the simulation scenarios in Fig. 14(a,b), (c,d) and (e,f), respectively

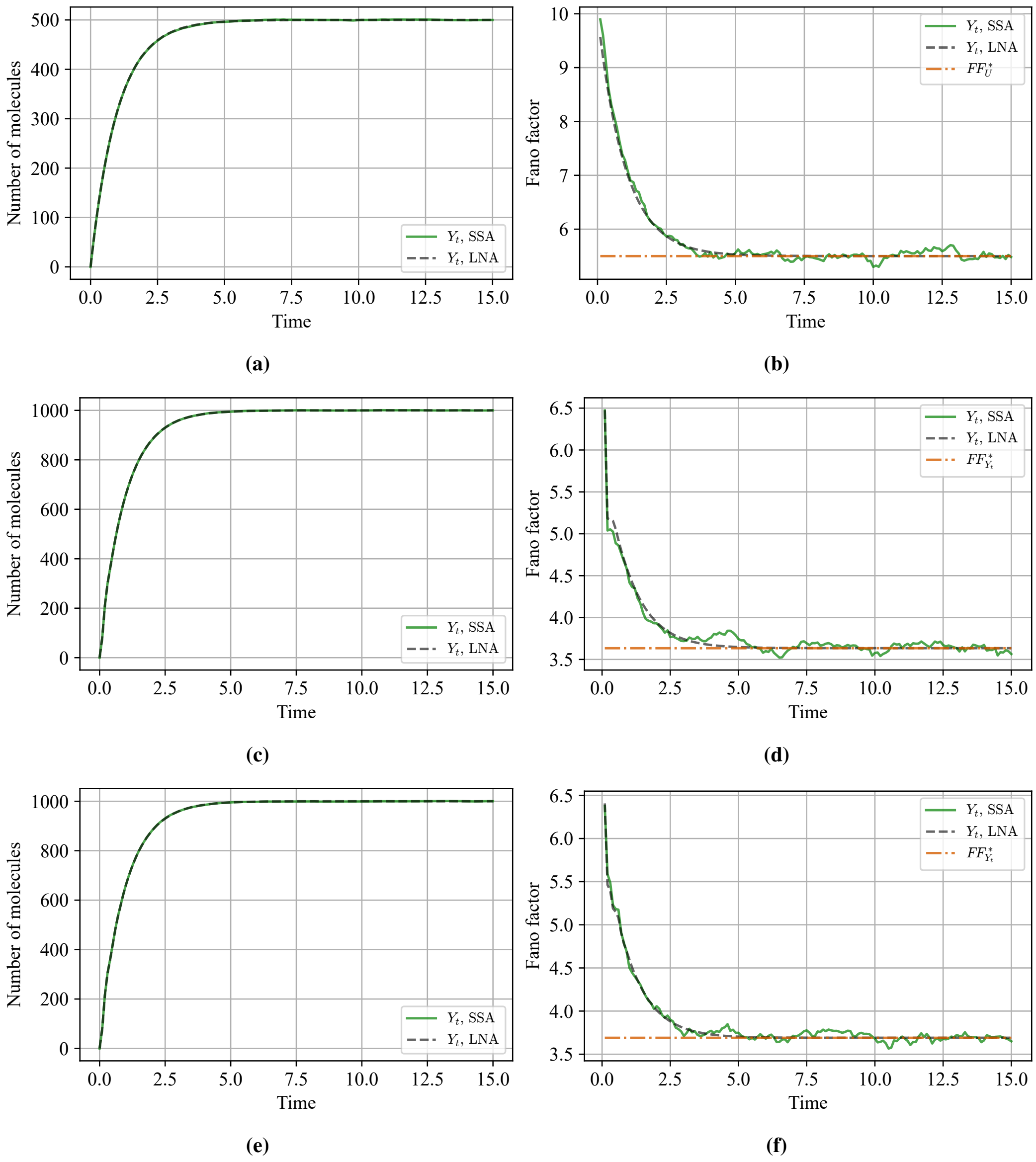

**Figure S15: Positive-feedback, derivative control.** Time evolution of the mean molecular copy number and the Fano factor (together with the corresponding analytically derived steady-state approximation), obtained using SSA and LNA for: (a,b) a target species ($Y_t$) participating in a general bursty birth–death biomolecular process (CRN in Eq. 3) with $c = 10$, $p_1 = 1$, and $p_2 = 1$ (process to be controlled); (c,d) the target species in (a,b) regulated by a *BioSD-II*-based controller (CRNs in Eq. 8) with $k_f = 0.5$, $b = 250$, $k_1 = 16$, $k_2 = 1$, $k_3 = 20$, $n = 1000$ and $k_{in} = 32$. (e,f) the target species in (a,b) regulated by an *AC-BioSD-II*-based controller (CRNs in Eq. 11) with $k_{in} = 1.6$ and all remaining parameters unchanged. The initial conditions in all cases are set to zero. Individual stochastic trajectories for the above simulation scenarios are presented in Fig. S16 of the Supplementary Material. Note that all parameters are identical to those in Fig. 14, except for $c = 10$.

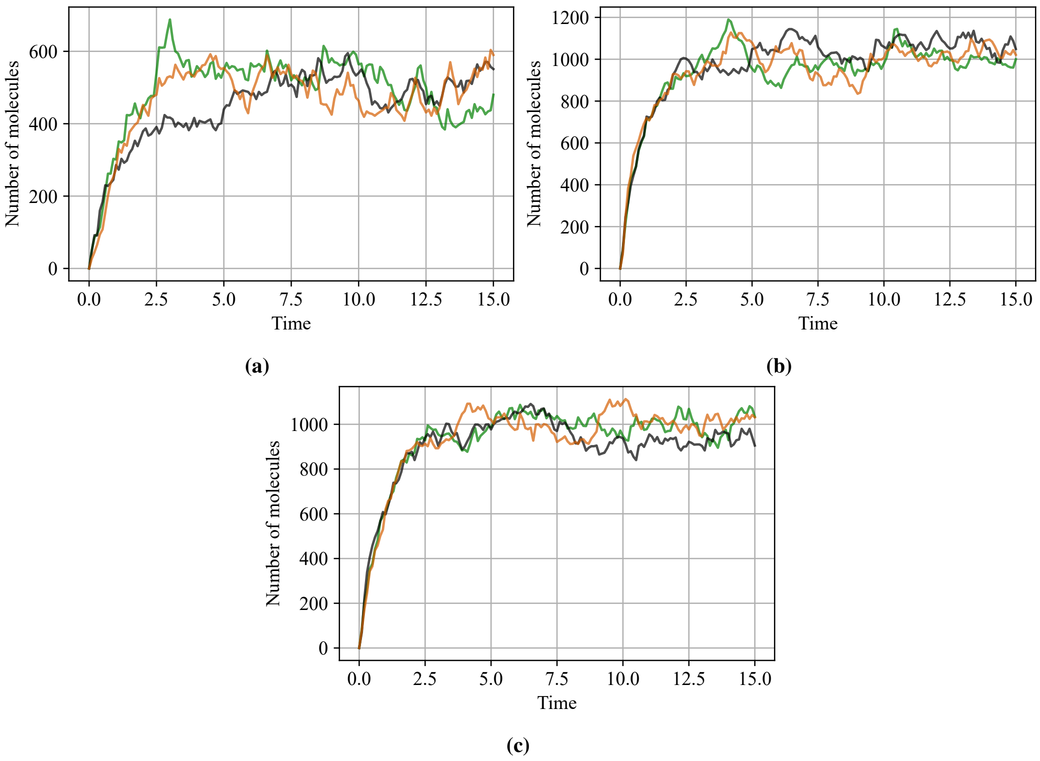

**Figure S16: Stochastic trajectories.** Panels (a)–(c) each display three individual stochastic trajectories for the simulation scenarios in Fig. S15(a,b), (c,d), and (e,f), respectively.
